# Disease-Causing Mutations Alter the Interplay Between Loop Dynamics and Catalysis in the Protein Tyrosine Phosphatases SHP-1 and SHP-2

**DOI:** 10.64898/2026.03.02.708844

**Authors:** Alfie-Louise R. Brownless, Yazdan Maghsoud, Michael Robinson, Shina C. L. Kamerlin

## Abstract

The protein tyrosine phosphatases (PTPs) SHP-1 and SHP-2 play complex roles in a variety of signaling pathways, including those involved in cancers and other diseases, making them important drug targets. These two PTPs have superimposable active sites, but different biological functions *in vivo*, including opposing roles in cancer development. Unique to these PTPs is the presence of two tandem Src homology 2 (SH2) domains, which regulate access to the phosphate-binding site in the catalytic domain through an autoinhibition mechanism. Studies of the allosteric regulation and dynamics of these PTPs, as well as associated drug discovery efforts, typically focus on autoinhibition rather than the dynamics of a catalytic loop in the phosphatase domain, the WPD-loop, which is essential for PTPase activity. However, recent deep mutational scanning data has demonstrated that oncogenic mutations also regulate WPD-loop motion in SHP-2. We provide here a detailed computational study of WPD-loop dynamics and catalysis in wild-type and mutant full-length and catalytic domain only SHP-1 and SHP-2, demonstrating that many oncogenic residues lie on the allosteric pathways regulating WPD-loop dynamics. Mutations at these positions alter WPD-loop dynamics, disrupting the active site and negatively impacting catalysis. Further, our simulations provide molecular insight into the link between the presence of the SH2 domains and loop motion in the catalytic domain, and, importantly, how it differs between the two PTPs. Taken together, our work showcases the impact of altered WPD-loop motion in disease-causing SHP-1 and SHP-2 variants, opening new strategies for selectively targeting these important therapeutic enzymes.

## Introduction

Protein tyrosine phosphatases (PTPs) are a large family of enzymes^1^ that regulate cellular signaling,^2^ thus frequently playing major roles in disease-related pathways.^3^ PTPs act in tandem with protein tyrosine kinases (PTKs) to phosphorylate and dephosphorylate their substrates,^4^ allowing for signal propagation across crucial molecular pathways. Both PTPs and PTKs play complex roles in regulating signaling. However, while wild-type PTKs exhibit positive effects on cell proliferation, PTPs can exhibit either positive or negative roles in proliferation, and are thus more variable in their roles in disease progression.^5^ Furthermore, due to their central roles in biology, aberrations in PTP function have been linked to human cancers, diabetes, autoimmune disorders, cardiovascular disorders, metabolic diseases, and neurological diseases.^3, 4, 6–10^ This makes PTPs crucial (but elusive) drug targets.^11–13^

A major challenge with targeting PTPs is the fact that their core catalytic (PTP) domain is highly conserved.^14^ Further, all PTPs share highly conserved active sites including a conserved phosphate-binding loop (P-loop, C(X)_5_R) motif,^15^ which makes inhibitor selectivity a particular challenge.^16, 17^ However, PTPs also carry a conserved mobile loop, the WPD-loop, which undergoes a substantive conformational change from catalytically “open” to a catalytically “closed” conformation.^18, 19^ In doing so, they position an (again) conserved aspartic acid side chain into the active site, that acts as a general acid/base in the two-step reaction catalyzed by PTPs (**Figure 1**).^19^ Importantly, despite the high conservation of the catalytic domain, active site structure, key catalytic loops, and chemical mechanism, the catalytic rates of PTPs vary by several orders of magnitude.^20^ This has been argued to be driven by differences in WPD-loop dynamics across PTPs.^21–23^ This makes the allosteric regulation of the WPD-loop of PTPs an attractive target for novel inhibitors that act by blocking proper closure of the WPD-loop into a catalytically competent position.^24–26^ Achieving this requires intimate understanding of the allosteric regulation of WPD-loop motion in the PTP of interest, and how this regulation is modified by disease-causing mutations.

**Figure 1.**
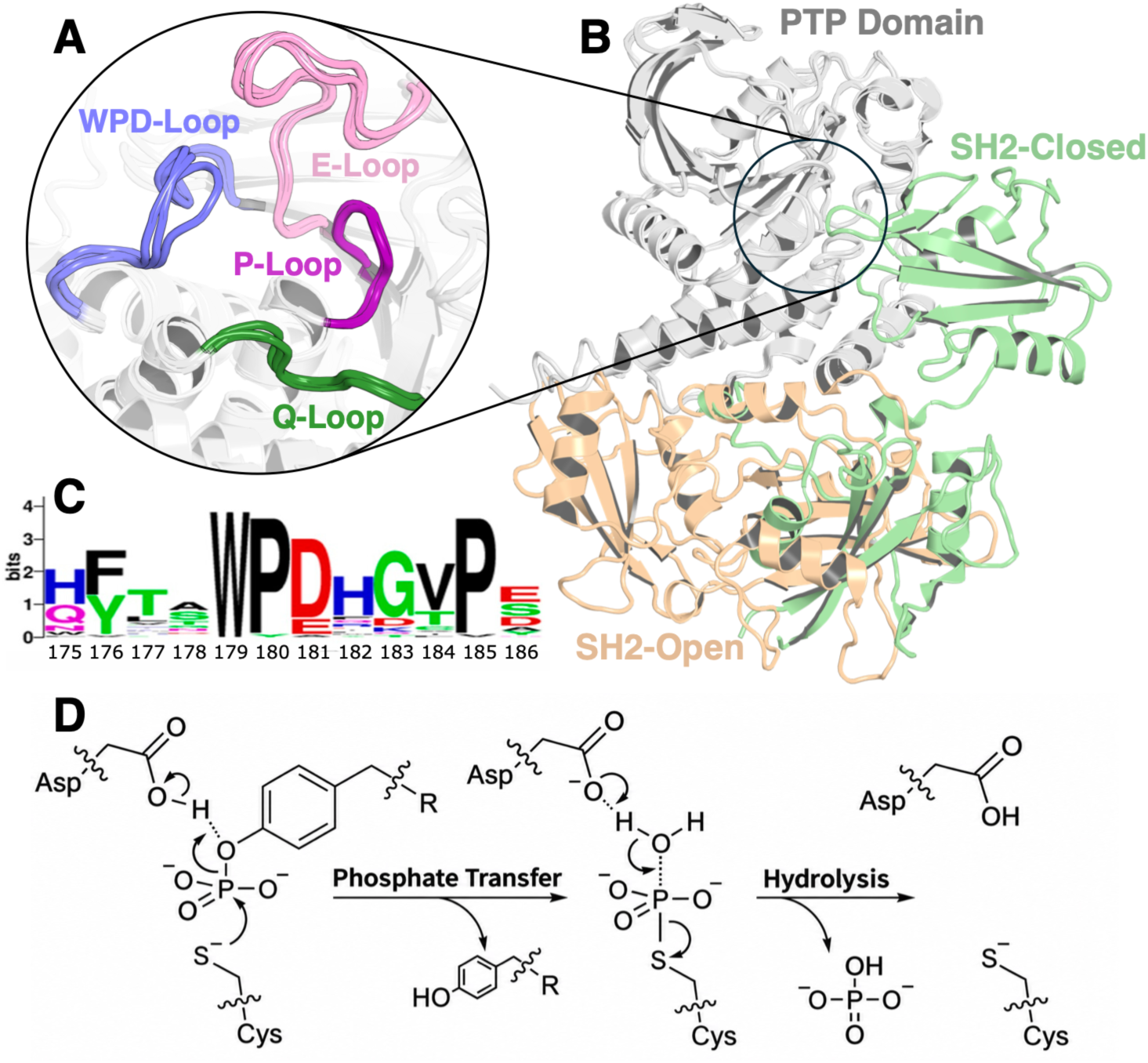
(**A**) Visual alignment of the PTP1B, SHP-1, and SHP-2 active sites with key active site loops emphasized as follows: WPD-loop (blue), P-loop (purple), Q-loop (green), E-loop (pink). (**B**) Overlay of the structure of SHP-1 with its Src homology 2 (SH2) domains in the catalytically active open conformation (tan) (PDB ID: 3PS5^27^), and in the autoinhibited closed conformation (green) (PDB ID: 2B3O^28^). (**C**) Sequence logo showing the amino acid conservation of the WPD-loop across a collection of PTPs, originally presented in ref. ^29^ under a CC-BY license. (**D**) Conserved two-step catalytic mechanism of PTPs, proceeding through acid-base catalysis involving an aspartic acid side chain located on the WPD-loop. The second (hydrolysis) step of this process has been shown or inferred to be rate-limiting in SHP-1, SHP-2 and analogous PTPs.^21, 30–33^ The WebLogo^34^ in panel (**C**) was originally presented in ref. ^29^ under a CC-BY 4.0 license. The catalytic mechanism in Panel (**D**) is adapted from ref. ^23^, again published under a CC-BY license. Both images copyright © The Authors. Published by the American Chemical Society.

The focus of this work is on the role of disease-causing mutations on regulating WPD-loop motion (and thus activity) in the protein tyrosine phosphatases SHP-1 and SHP-2. Both enzymes have complicated relationships to cancer signaling, as both SHP-1 and SHP-2 can act as both oncogenes and as tumor suppressors, depending on biological context.^7, 35–39^ However, their prominent role in oncogenesis makes them attractive anticancer drug targets.^10, 36, 38, 40, 41^ Further, gain-of-function mutations of SHP-2 have been shown to cause growth disorders^42^ and tumors.^43^ Interestingly, and uniquely to these enzymes, SHP-1 and SHP-2 are the only cytosolic PTPs to express two tandem Src homology 2 (SH2) domains,^44^ which are involved in autoinhibition of the enzyme, regulating access to the phosphate binding site in the catalytic domain (**Figures 1B** and **S1**).^27, 28, 45–47^ Curiously, despite the fact that SHP-1 and SHP-2 are an evolutionary pair of enzymes,^19^ with both high sequence conservation (59% overall, 60% across their catalytic domains) and structural conservation,^14^ the impact of the SH2 domains on the kinetics of each enzyme is very different. In SHP-1, the presence of the SH2 domains primarily impacts *K*_M_ rather than *k*_cat_ as shown by comparison to the activity of the truncated phosphatase domain.^30^ In contrast, in SHP-2, the SH2 domains also have a significant impact on *k*_cat_,^48^ suggesting a difference in active site loop dynamics. We note that in both SHP-1 and SHP-2, the C-terminal tail also plays an important role in regulation.^44^

Understanding the origins of these differences in catalytic loop dynamics, how they are regulated by the SH2 domains in the two enzymes, and how they are impacted by oncogenic mutations is an important step towards being able to develop selective inhibitors preferentially targeting only one of the two enzymes. To this end, we initiate our study with a molecular dynamics comparison of the intrinsic dynamics of the WPD-loop open forms of the catalytic domains of SHP-1 and SHP-2 to that of the canonical PTP, PTP1B,^3, 11, 22, 23, 32, 49–56^ and show that WPD-loop dynamics in SHP-1 and SHP-2 are different not just between the two enzymes but also different from that of PTP1B. We then demonstrate that, consistent with kinetic measurements,^30, 48^ from a conformational perspective, the SH2 domains have a differential impact on WPD-loop dynamics of full-length SHP-1 versus their impact in SHP-2, despite sequence similarities across both the catalytic and SH2 domains. Further, we present a detailed comparison of the dynamics of the catalytic domains of select pathogenic variants of SHP-1 and SHP-2. Here, we demonstrate that mutations at positions on the allosteric pathways regulating WPD-loop motion in these phosphatases alter the ensemble and conformational transitions of the WPD-loop, despite these positions being distal to the active site. Finally, we leverage empirical valence bond^57^ (EVB) simulations to characterize the impacts of chosen pathogenic variants’ chemical barriers in the rate-limiting step of the generalized PTP mechanism. We note that in both enzymes, both the distributions of loop conformations and reaction barriers are shifted in the pathogenic variants. Taken together, our work shows the sensitivity of WPD-loop motion across PTPs in particular to WPD-loop sequence, resulting in significant differences in activity and regulation of these enzymes. This shows how nature can use a simple structural motif to regulate catalysis in a broad range of biological contexts and provides an important target for future drug discovery efforts.

## Results and Discussion

### Contrasting WPD-loop dynamics across SHP-1, SHP-2, and PTP1B

SHP-1 and SHP-2 share a high degree of sequence similarity – 59% overall primary sequence identity, as well as 60% similarity in their catalytic domains,^14^ having evolved as an evolutionary pair.^19^ This is much higher than the sequence identity, for instance, between SHP-1 and PTP1B (38%) or between SHP-2 and PTP1B (39%) (all values calculated using MUSCLE,^58^ aligning on the catalytic domain). Given the high sequence similarity between SHP-1 and SHP-2, and the fact that PTPs tend to have very highly structurally conserved catalytic domains overall,^14^ it would stand to reason that SHP-1 and SHP-2 would have WPD-loop dynamics that are similar to that of each other, but different (potentially) to that of PTP1B. However, there are differences in both the sequences of the WPD-loop and adjacent residues in these PTPs (**Figure 1C**), which could lead to different WPD-loop dynamics, and perhaps should be expected from the observation that SHP-1 and SHP-2 have distinct non-redundant biological functions.^19^

To address this question, as our starting point, we have compared the dynamical behavior of the WPD-loop in the catalytic domains of SHP-1 and SHP-2, in the absence of any autoinhibition from the SHP-1/2 SH2 domains. To this end, we have performed conventional MD simulations of the open WPD-loop / unliganded states of each enzyme, as outlined in the **Materials and Methods**. This allows us to isolate the intrinsic conformational flexibility of the catalytic domains of these enzymes, including that of the WPD-loop, without confounding factors introduced by interdomain interactions with the SH2 domains. Further, prior work has indicated that while the WPD-closed conformation of PTPs is clearly defined, these enzymes sample a range of interconverting loop-open conformations,^23, 56, 59^ with a link between increased flexibility of the open conformation of the WPD-loop, and correspondingly increased PTPase activity.^21^ Thus, the relative conformational ensembles of the WPD-loops of the unliganded PTPs provide an important window into function.

Comparison of C_α_-atom root mean square fluctuations (RMSF) during MD simulations of SHP-1 and SHP-2 shows relatively similar overall local flexibility profiles for the two enzymes, with mobility differences confined to two loop regions that are distal from the active site (**Figure S2**). Despite this overall baseline similarity, differences emerge when comparing WPD-loop flexibility between the two PTPs. **Figure 2** shows joint probability distributions projecting trajectory frames onto predefined geometric coordinates (dRMSD *vs.* native contacts between the P- and WPD-loops, **Figure 2A**), and the first two time-lagged independent components (tIC1 *vs*. tIC2) derived from linear dimensionality reduction (**Figure 2B**, for details see the **Materials and Methods**, and for a list of pairwise distances used to calculate the dRMSD see Zenodo DOI: 10.5281/zenodo.18817309). A plot of implied timescales *vs*. lag time for our tICA analysis is shown in **Figure S3** (note that here, tICA is being used *solely* as a linear dimensionality reduction technique to identify slow coordinates for conformational space visualization, and *not* to extract kinetic data). Based on these data, we observe distinct dynamic behavior in the three enzymes: despite initiating all simulations from WPD-loop open states, unliganded SHP-1 exhibits extensive sampling of intermediate and closed-like WPD-loop conformations, whereas the SHP-2 ensemble remains tightly localized in more open WPD-loop conformations, with minimal sampling of loop-closed states. Finally, and in contrast, like SHP-1, PTP1B samples both open and closed WPD-loop conformational states, but remains more conservative in the space sampled, with sampling predominantly localized on the WPD-loop open confirmation. Thus, PTP1B serves as a conformational intermediate between the two SHPs. Representative spatial ensembles corresponding to these basins are shown visually in **Figure 2C**.

We note that the rotation of the WPD-loop tryptophan has been identified as a crucial step in WPD-loop closure, with differences in Trp behavior between species,^21, 60^ and with this residue acting as a hinge/pivot point for loop motions. **Figure S4** shows the joint probability distributions of each of the WPD-loop Trp *χ*_1_ and *χ*_2_ dihedral angles *vs.* the center of mass distance between the WPD-loop and P-loops of PTP1B, SHP-1 and SHP-2. In all three PTPs, the tryptophan side chain remains in a trans conformation on *χ*_1_ (∼180 - 200°), with PTP1B showing two clear basins corresponding to *χ*_2_ values of ∼ -90° (perpendicular) and 0° (flat), indicating that even in the unliganded form of the enzyme, the Trp sidechain is undergoing transitions between two distinct rotameric states, a stable t-90° state (χ_1_ = +180°, χ_2_ = -90°), with the indole ring perpendicular to the C_α_-C_β_ bond,^62^ and a sterically hindered t-0° conformation (χ_1_ = +180°, χ_2_ = 0°).^62^ In contrast, in unliganded SHP-1, we observe a single broad extended basin, with only subtle sub-state partitioning, and in unliganded SHP-2 we observed a narrower basin with sampling of the preferred t-90° state dominating. This is particularly interesting in the context of prior work which noted that the crystal structure of W179F PTP1B carries both open and closed states of the loop, and suggested that in PTP1B, the indole ring of the tryptophan and the WPD-loop move in the same direction.^21^ This motion appears to be dampened in SHP-1 and SHP-2 on the μs timescale, yielding a broader and less discrete rotameric ensemble in the unliganded state.

**Figure 2.**
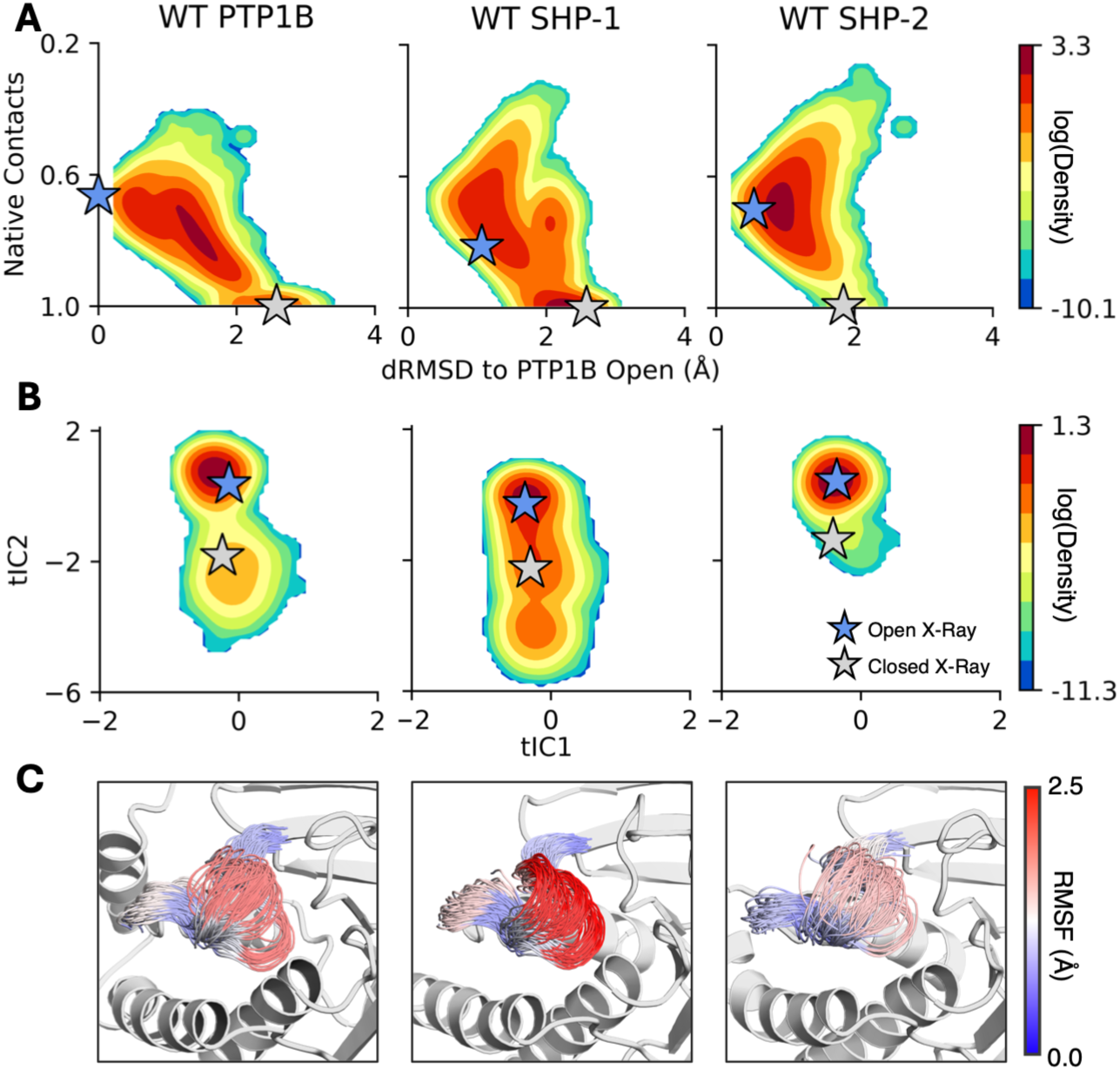
(**A**) Joint probability distributions of distance RMSD (dRMSD) to the open WPD-loop state of PTP1B (Å) and native contacts^61^ with relation to the closed WPD-loop state of PTP1B. (**B**) Joint probability distributions of the first two components generated from a time-lagged independent component analysis (TICA) fitted on all pairwise C_α_-atom distances between the WPD-loop and P-loops and sine and cosine values for the W417 (SHP-1 numbering) *χ*_1_ and *χ*_2_ dihedral angles. (**C**) Projection of WPD-loop positions evenly spaced across the range center of mass distances between the WPD-loop and P-loops. Residues colored according to root mean squared fluctuation (RMSF, Å) values. All results collected from 8 x 1.5µs molecular dynamics simulations initiated from the WPD-loop open unliganded state of PTP1B, and the SHP-1 and SHP-2 catalytic domains.

While the importance of the WPD-loop Trp is particularly pronounced in driving WPD-loop motion, several other structural features correlate with the motion of the WPD-loop, specifically (1) the E-loop, which carries a glutamic acid that forms a salt-bridge with the P-loop arginine,^23, 56, 63^ (2) a pre-organized “Q-loop” that carries a glutamine side chain that positions the nucleophilic water molecule for hydrolysis of the phosphocysteine intermediate,^64^ as well as (3) key helices including the α4-helix, which is connected to the P-loop, the α3-helix, which connects the WPD-loop to an allosteric site that can disrupt WPD-loop motion^25, 53^, the α5-helix, which flanks the Q-loop and is important to the binding of the allosteric inhibitor, BB3 to PTP1B,^25, 65^ and the allosterically important C-terminal α7-helix in PTP1B, which undergoes order-disorder transitions in correlation with WPD-loop motion.^53, 66, 67^ The position of these structural features is shown in **Figure 3**.

Dynamic cross-correlation maps^68^ (DCCM, calculated using CPPTRAJ^69^) and block-averaged linear mutual information analysis^70^ (**Figure 3A**) show distinct correlation profiles for the three enzymes. In all three PTPs, we observe extensive off-diagonal correlated motions, aligning with the expected correlated scaffold dynamics of the canonical PTP-fold (**Figure 3A**). Unexpectedly, however, SHP-1 uniquely shows prominent off-diagonal anti-correlation, in particular between the WPD-loop / P-loop and surrounding regions (**Figure 3B**), suggesting anti-phase motion between catalytic loops and distal surrounding helices. Further, in PTP1B, E-loop motion is strongly correlated with the α4-helix, in SHP-1 the E-loop and α4-helix show mixed correlated/anti-correlated behavior, and in SHP-2 the motion of these two regions is decoupled. In contrast, in both PTP1B and SHP-2, the motion of the WPD- and P-loops is strongly correlated, whereas in SHP-1 it is anti-correlated.

**Figure 3.**
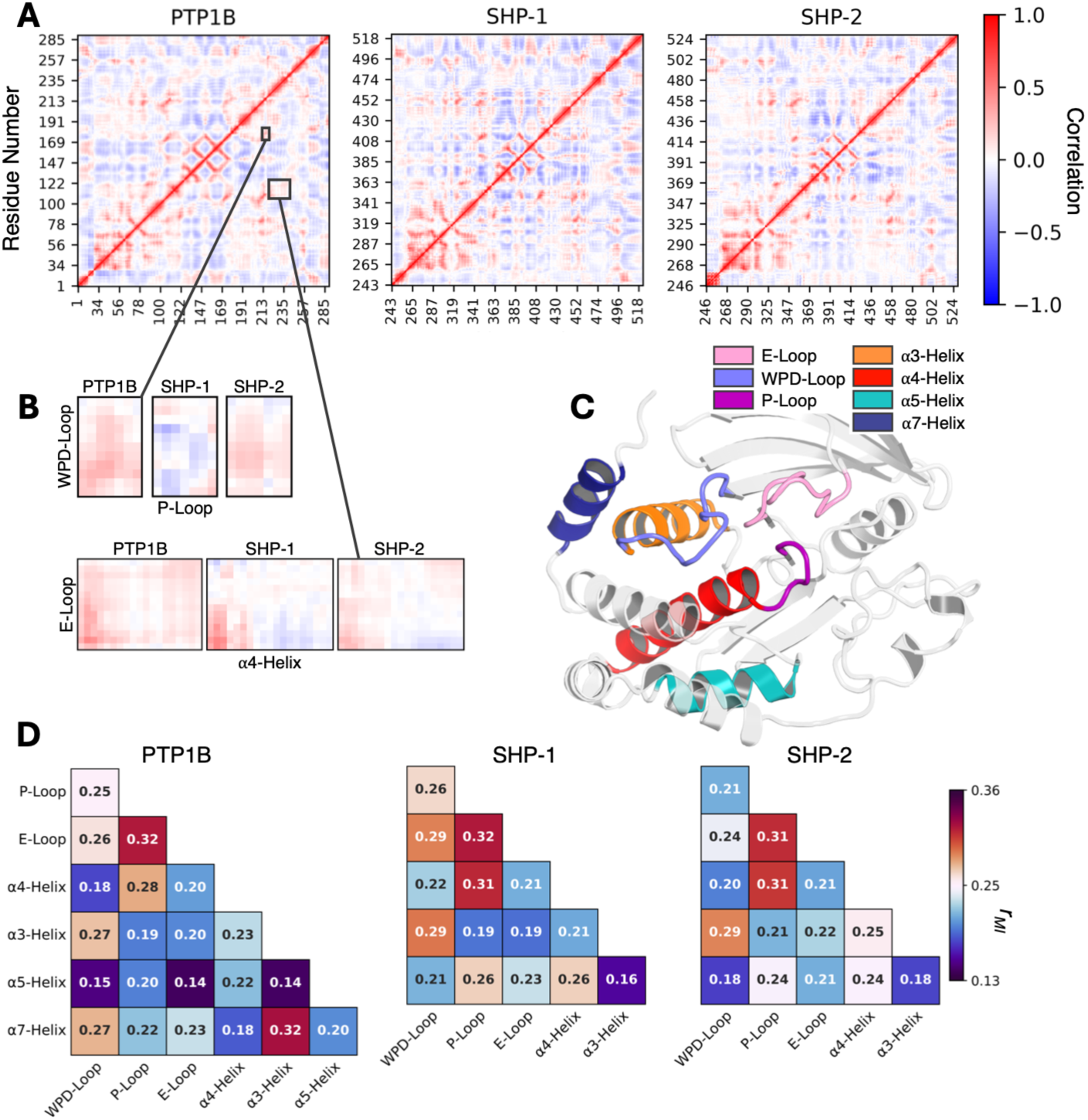
(**A**) Dynamic cross correlation matrices (DCCM) calculated based on simulations of PTP1B, and the SHP-1 and SHP-2 catalytic domains. (**B**) Highlighted DCCM plots for P-Loop *vs*. WPD-loop and α4-helix *vs*. E-loop. (C) Overview of the structural features being considered in this analysis. (**D)** Νormalized linear mutual information scores (*r*_MI_) computed among regions of interest for PTP1B, SHP-1, and SHP-2. The full list of calculated *r*_MI_ results is provided in **Table S1**.

Calculation of normalized linear mutual information (*r*_MI_) to quantify non-linear thermodynamic communication across the catalytic domain (**Figure 3** and **Table S1)** further indicates a coupled core between the P-loop and E-loop, and between the P-loop and the adjacent α4-helix (*r*_MI_ ∼ 0.3 across systems). In contrast, the extent to which distal elements integrate with WPD-loop motion varies across enzymes. In unliganded SHP-1, mutual information between the WPD-loop and E-loop is elevated (*r*_MI_ = 0.29 ± 0.02) relative to SHP-2 (*r*_MI_ = 0.24 ± 0.01), as is dynamic coupling between the WPD-loop and the α4-helix (*r*_MI_ = 0.22 ± 0.01). This enhanced global information transfer qualitatively correlates with the unique propensity of unliganded SHP-1 to sample WPD-loop closed states (**Figure 2**). In contrast, PTP1B exhibits a distinct and highly localized allosteric network, due to the strong coupling of its unique C-terminal α7-helix with the α3-helix (*r*_MI_ = 0.32 ± 0.01) and WPD-loop (*r*_MI_ = 0.27 ± 0.02). This is significant because although we simulated WPD-loop open PTP1B with an ordered α7-helix for simplicity, this strong coupling is consistent with the tendency of this helix to undergo order-disorder transitions with WPD-loop motion.^53, 66, 67^ Finally, and notably, SHP-2 is distinctive because beyond the flanking α3-helix, the WPD-loop is largely decoupled from distal structural elements (*r*_MI_ 0.18 – 0.24), contributing to the open ensemble observed in our conventional MD simulations (**Figure 2**).

These distinct dynamic differences in the catalytic domains of the three PTPs are important for two reasons: (1) the shifted conformational ensembles of the WPD-loops of SHP-1 *vs*. SHP-2 provide structural rationale for the orthogonal effects of the flanking SH2 domains on activity,^30, 48^ and (2) although SHP-1 and SHP-2 have evolved as an evolutionary pair,^19^ their catalytic domains have substantive differences in dynamical behavior, with PTP1B unexpectedly acting as a dynamical intermediate between the two enzymes.

### Impact of the SHP-1 and SHP-2 Tandem SH2 Domains on WPD-Loop Dynamics

As shown in **Figures 1** and **S1**, SHP-1 and SHP-2 both carry tandem SH2 domains, and are in fact the only cytosolic PTPs to do so.^44^ Both proteins exhibit an autoinhibition mechanism whereby in the catalytically active ‘open’ state, the PTP domain is free to bind to substrate, while in the catalytically inactive ‘closed’ state, the active site is effectively blocked by the SH2 domains (**Figure S1**). Despite the similar mechanisms of autoinhibition of SHP-1 and SHP-2,^27, 28, 45–47^ however, the SH2 domains exert orthogonal effects on enzyme kinetics. Specifically, comparisons of the activities of full length and truncated (catalytic domain) versions of SHP-1 and SHP-2 have shown that in SHP-1, the presence of the SH2 domains primarily impacts *K*_M_ with *k*_cat_ largely unaffected,^30^ whereas in SHP-2, the *k*_cat_ of full-length SHP2 is reduced to 4.8% of the activity of the truncated catalytic domain.^48^ Given the importance of WPD-loop motion to catalysis and turnover in PTPs,^19, 22^ this suggests that the presence of the SH2 domains may have minimal impact on WPD-loop dynamics in SHP-1 (compared to the corresponding dynamics in only the truncated catalytic domain), whereas they likely have a substantive impact on WPD-loop dynamics in full length SHP2. This hypothesis is supported by differences in intrinsic WPD-loop dynamics observed in our analysis of the SHP-1 and SHP-2 catalytic domains (**Figures 2** and **3**).

The focus of this section is on comparing dynamics of full length SHP-1 and SHP-2 in the SH2-domain open conformations (their active conformations with an accessible catalytic site), with the intrinsic dynamics of the catalytic domain only. While the autoinhibition of these SHP phosphatases has been extensively studied computationally.^24, 71–83^ WPD-loop motion has largely been understudied (although some prior work has touched on it, *e.g.*, ^84^). However, while allosteric regulation of access to the catalytic domain is clearly one important component of the activity of these enzymes, WPD-loop motion controls the subsequent catalytic activity of the activated PTPs, and thus is an equally important component. We note that while there exist full-length structures of SHP-1 in both catalytically active and autoinhibited conformations of the SH2 domains (PDB IDs: 3PS5^27^ and 2B3O,^28^ respectively), the only structures of wild-type full-length SHP-2 are in its autoinhibited conformation (PDB ID: 4DGP^47^). Thus, as our simulation starting points and for consistency, we generated plausible Boltzmann ensembles of open WPD-loop full length SHP-1 and SHP-2 seeded from the corresponding autoinhibited structures using BioEmu,^85^ as described in the **Materials and Methods**.

**Figure 4** shows a comparison of the conformational ensembles sampled by full-length and truncated (catalytic domain only) SHP-1 and SHP-2 during our respective simulations. tICA analysis was again performed using the combined pairwise distances between the WPD- and P-loop C_α_-atoms, and the sine/cosine values of both the *χ*_1_ and *χ*_2_ angles of the WPD-loop Trp side chain. Here, we observable notable differences between the full-length SHP-1 and SHP-2 conformational ensembles. Specifically, in the isolated catalytic domain, SHP-1 already shows a tendency to sample more closed WPD-loop conformations even in the unliganded states (**Figure 2**); this tendency appears to be amplified by the presence of the SH2 domains. In contrast, the SH2 domains appear to expand the conformational ensemble of the SHP-2 WPD-loop, allowing it to access a range of open, intermediate, and more closed conformations of the WPD-loop, accompanied by a range of conformational states of the WPD-loop Trp, which are not observed in our SHP-1 simulations. Further, we observe that the N-SH2 domain of SHP-2 explores a broader spatial ensemble relative to the catalytic domain than that of SHP-1 (**Figure S5**), in agreement with experiment.^28, 42, 86^ These differences qualitatively align with the observations that the SHP-1 SH2 domains have minimal impact on *k*_cat_, whereas in SHP-2, the impact of the SH2 domains is substantive,^30, 48^ and that the two phosphatases have different biological functions *in vivo*.^44^

**Figure 4.**
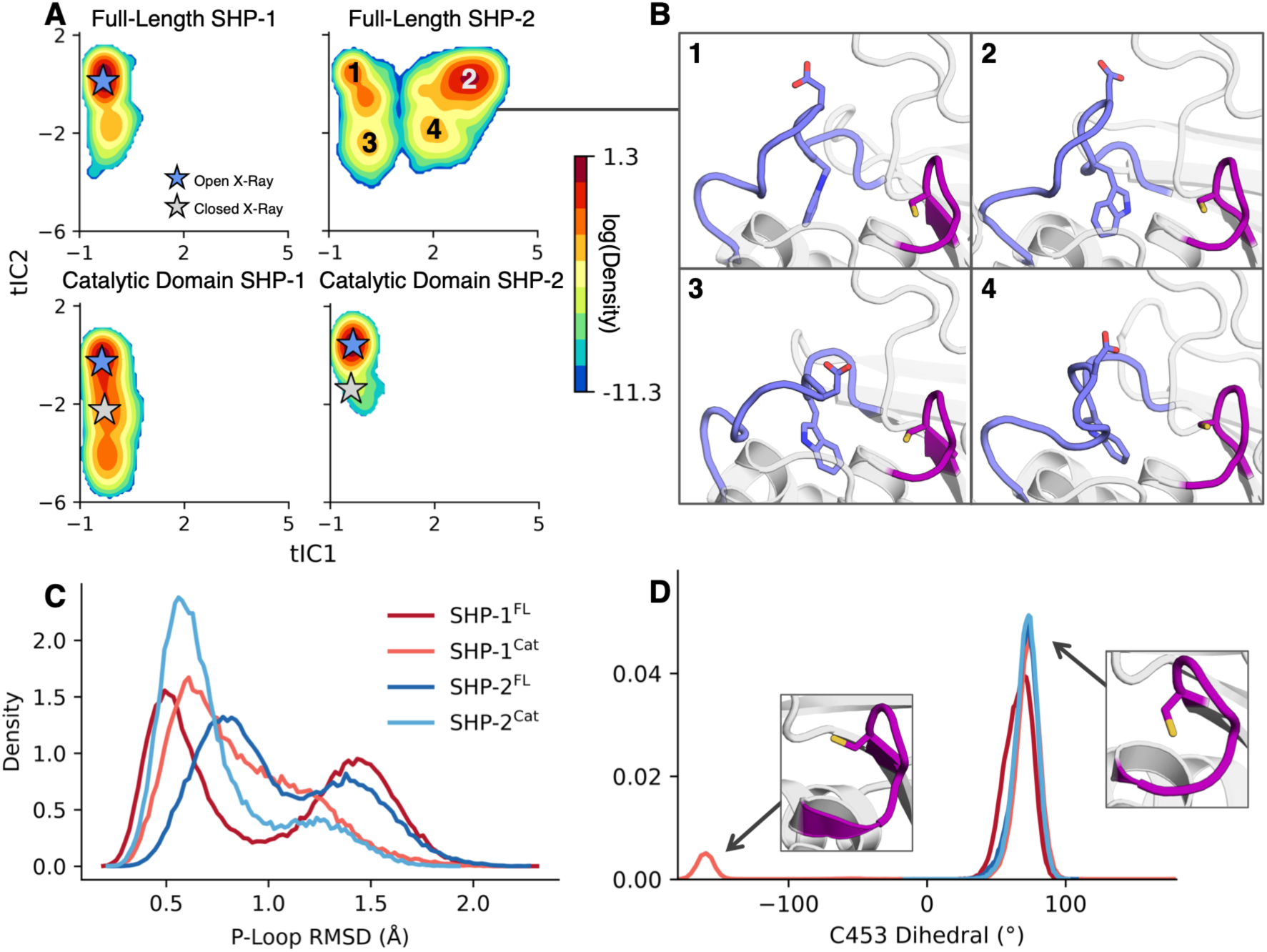
(**A**) Joint probability distributions of the first two components generated from a time-lagged independent component analysis (tICA) fitted on all pairwise C_α_-atom distances between the WPD-loop and P-loops, and sine and cosine values for the W417 (SHP-1 numbering) χ1 and χ2 side chain dihedral angles. (**B**) Representative structures are four selected centroids from K-means clustering (N=6) observed in full-length SHP-2. (**C**) Histogram of the root mean squared deviation (RMSD) of the P-loop Cα-atoms for each of full length SHP-1 and SHP-2, and the SHP-1 and SHP-2 catalytic domains. (**D**) Distribution of the catalytic cysteine (C453 SHP-1 numbering) side chain dihedral angles for each SHP-1 full length, SHP-2 full length, SHP-1 catalytic domain, and SHP-2 catalytic domain.

As a minor point, curiously, our simulations of the full-length SHP phosphatases also indicate enhanced flexibility of the catalytic P-loop (bimodal RMSD distributions shown in **Figure 4C**), suggesting that the presence of the SH2 domains allow these enzymes to sample alternate active-site conformations necessary for substrate engagement. In conjunction with this, the truncated form of SHP-1 appears to occasionally sample unproductive conformations of the catalytic cysteine, (**Figure 4D**), that are dampened in the full-length enzyme. This qualitatively aligns with the observation that the SH2 domains have a greater impact on shielding from cysteine oxidation in SHP-1 compared to SHP-2.^87^

### Disease-Causing Mutations Modulate WPD-Loop Dynamics via Altered Allosteric Networks

There are an increasing range of methods with which to computationally characterize protein allostery.^88–90^ In prior work,^23^ we characterized allosteric communication pathways in both PTP1B and a bacterial PTP, YopH, using the shortest path map approach (SPM).^91^ This analysis demonstrated high conservation of allostery between these enzymes, with structure-based sequence alignment indicating that 35 of 69 SPM residues identified in PTP1B are conserved in YopH, despite only 17.5% sequence similarity between the two enzymes.^23^ PTP1B and SHP-1 have 38% sequence similarity, and the SHP-1 and SHP-2 PTP domains share even higher sequence similarity (60%, calculated using MUSCLE^58^). Projection of SPM residues onto structure-based sequence alignment of all four enzymes (**Figure 5**) shows suggests that, despite the differences in WPD-loop dynamics in these PTPs (**Figure 2**), >50% of SPM residues are conserved between SHP-1 and SHP-2 (23/41), and ∼40-50% of SPM residues are conserved between pairs of SHP-1/SHP-2 and PTP1B (**Table S2**). This supports an evolutionary core conserved across the three enzymes (which is highest among SHP-1 and SHP-2, which are an evolutionary pair^19^), in agreement with other work indicating conserved allostery across PTPs.^23, 92, 93^ Our analysis also identified several SPM residues that are unique to each isoform, as summarized in **Figure S6**. We note that N303 in SHP-1 has been described to help stabilize the active site,^27^ and that modifications at the N308, L261, and L262 position on SHP-2 are known to cause disease.^26, 48, 94–99^ Given that targeting conserved active-site or allosteric core residues across PTPs can lead to off-target cross-reactivity, identifying unique SPM positions can be helpful for the rational design of isoform-selective allosteric inhibitors.

**Figure 5.**
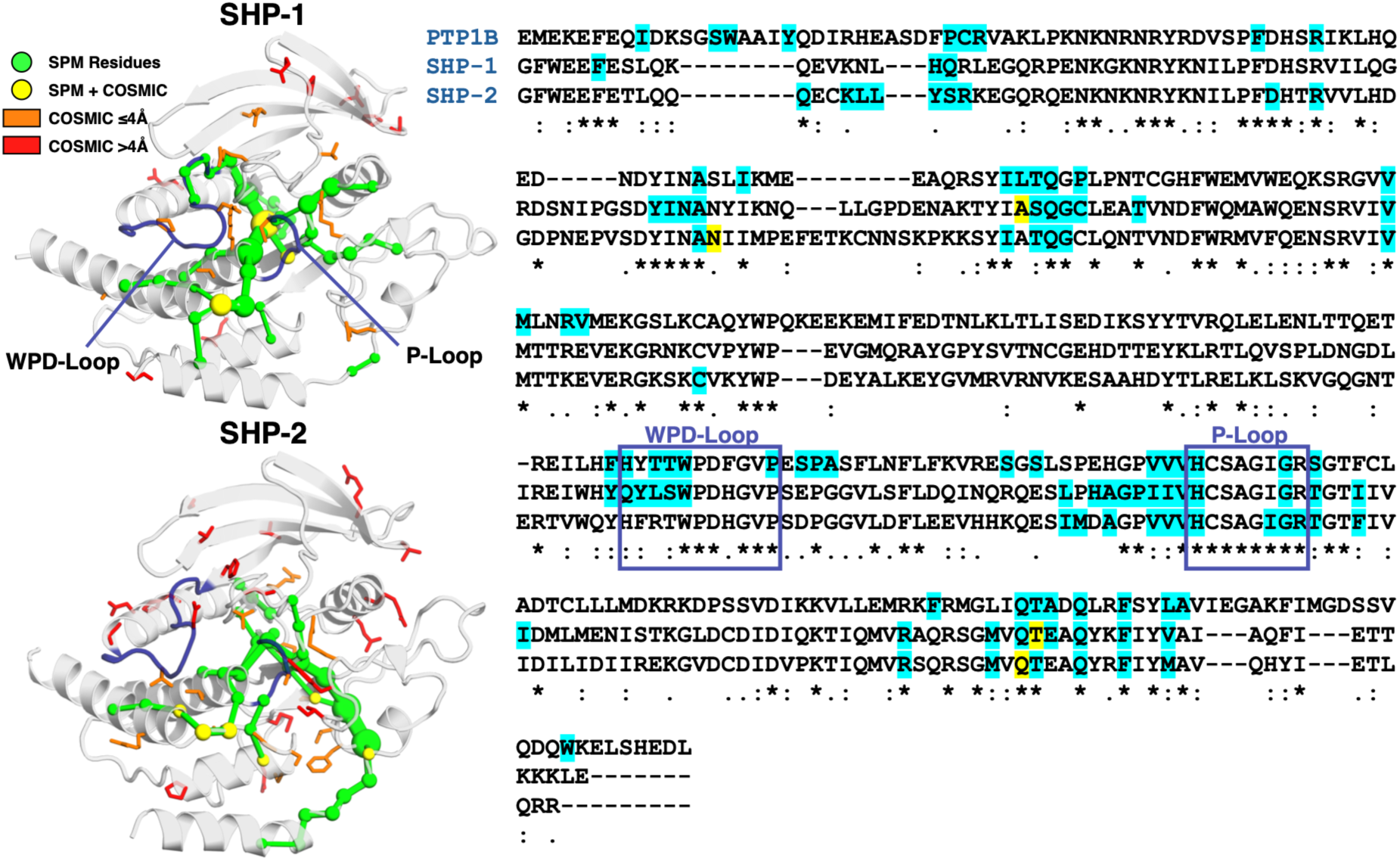
Identification of pathways utilized for allosteric communication in SHP-1 and SHP-2. **(left)** The shortest path map^91^ (SPM) obtained from our conventional MD simulations of SHP-1 and SHP-2. Overlaid onto the graph are the residues with at least 2 known oncogenic mutations (obtained from the COSMIC database^96^) for SHP-1 and SHP-2. Mutations are in yellow if found on the SPM itself, orange if the closest atom distance to an SPM residue is <4 Å, and red the distance is ≥4Å. The positions of mutations selected for further analysis on the scaffold (A323T and T501M in SHP-1, and N308D and Q506P in SHP-2) are annotated. (**right**) Structure-based sequence alignment of PTP1B, SHP-1, and SHP-2, with SPM residues highlighted in blue. Between 41 and 56% of SPM residues are conserved between enzyme pairs (see **Table S2**). The WPD-loop and P-loop sequences are highlighted with boxes. Chosen disease-causing positions A323 and T501 (SHP-1) and N308 and Q506 (SHP-2) are highlighted in yellow.

Following from this, and based on prior work that links oncogenic mutations to WPD-loop mutation in SHP-2,^100^ we compared the calculated allosteric pathways obtained by SPM analysis conducted on our WPD-loop open state conventional MD simulations of SHP-1 and 2 with the locations of known oncogenic PTP-domain amino acid substitutions in the COSMIC database, as of 12^th^ of October 2026.^96^ Based on COSMIC data, we identified 23 positions in the catalytic domain of SHP-1 and 39 positions in the catalytic domain of SHP-2 as being affected by two or more reported oncogenic amino acid substitutions. As can be seen from **Figure 5 and Tables S2** and **S3**, many (13/23 in SHP-1, and 21/39 in SHP-2) of the residues important to oncogenesis in these PTPs are directly or closely involved in an allosteric pathway.

Based on this analysis, we selected the T501M and A323T substitutions in SHP-1, and N308D and Q506P substitutions in SHP-2 as variants of interest for further study, as they are both (1) located on the calculated SPM, and (2) known from either the literature or from the COSMIC database^96^ to be important to SHP-1/SHP-2 function (T501M and A323T in SHP-1 are both associated with adenocarinoma,^38, 96^ and N308D and Q506P in SHP-2 are additionally known to be gain-of-function Noonan/Leopard syndrome substitutions).^48, 97^ The positions of these amino acid substitutions relative to the SHP-1/SHP-2 active sites are shown in **Figures 5** and **S7**. For each variant of interest, we have performed 8 x 1.5µs MD simulations of the truncated unliganded catalytic domain of each PTP, initiated from the open WPD-loop conformation, as in our simulations of the wild-type enzymes (**Figure 2**). Excluding the SH2-domains allows us to test for direct links between these substitutions and WPD-loop motion, without the additional layer of the SH2-autoinhibition mechanism (*i.e.*, it allows us to decouple the two effects).

**Figure 6.**
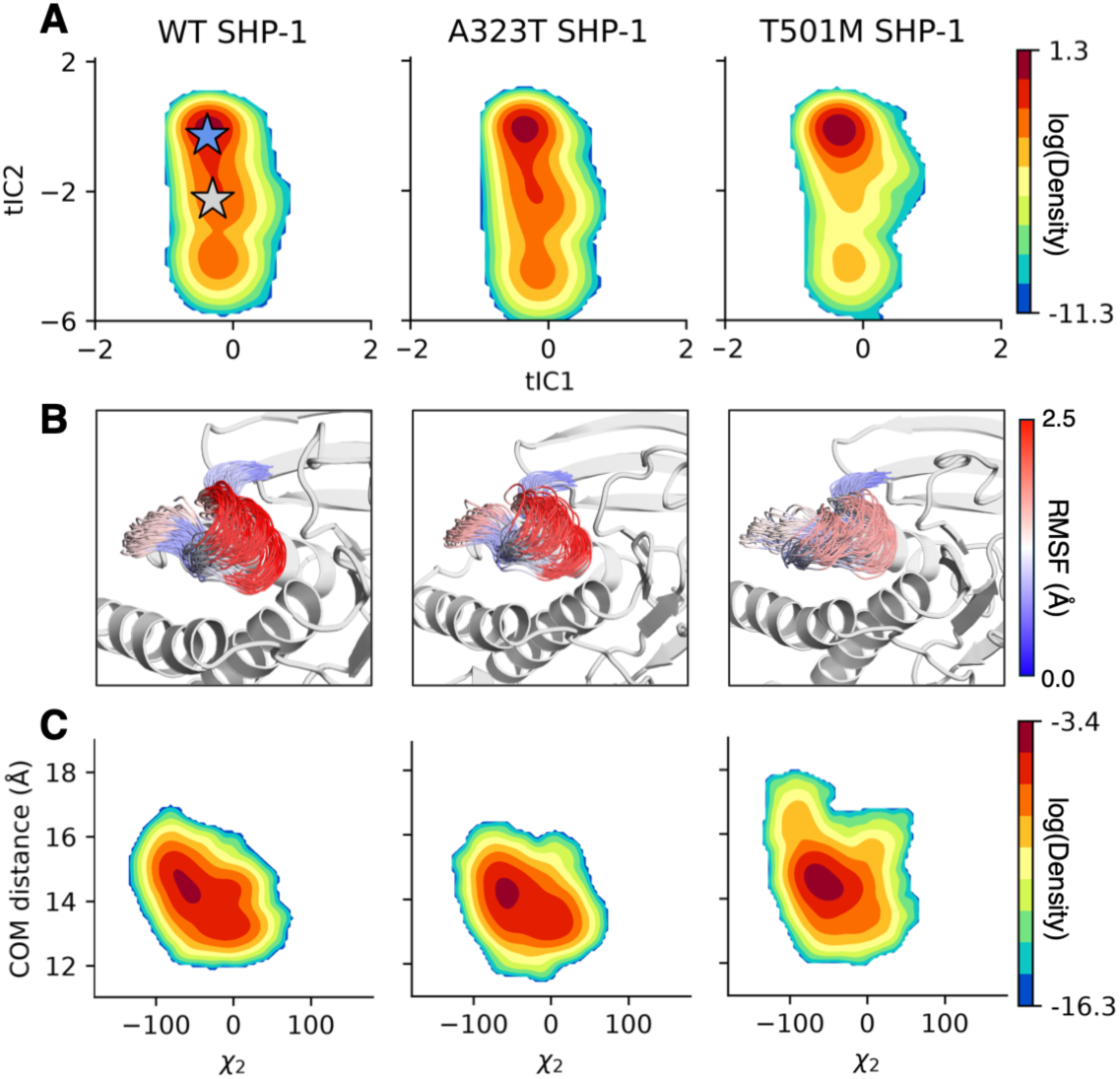
(**A**) Joint probability distributions of the first two time-independent components (tICs) generated from time-lagged independent component analysis (tICA) fitted on all pairwise C_α_-atom distances between the WPD-loop and P-loops, and sine and cosine values for the W417 (SHP-1 numbering) χ_1_ and χ_2_ side chain dihedral angles. The blue star indicates the open crystal structure, and the gray star the closed crystal structure. (**B**) Projection of WPD-loop positions evenly spaced across the range center of mass (COM) distances between the WPD-loop and P-loops. Residues colored according to root mean squared fluctuation (RMSF) values. (**C**) Joint probability distribution of W417 (SHP-1 numbering) χ_2_ side chain dihedral angles *vs*. the COM distance between the WPD-loop and P-loop. See **Figure S4** for the corresponding probability distributions in the wild-type enzyme and **Figure S8** for corresponding distributions of the χ_1_ side chain dihedrals in the variants. All results collected from 8 x 1.5µs molecular dynamics simulations initiated from the WPD-loop open unliganded state.

In the case of SHP-1 (**Figure 6**), the two substitutions appear to have orthogonal impact on the conformational ensemble of the WPD-loop. That is, while A323T broadens the ensemble along tIC2 (describing the conformational transitions of the WPD-loop Trp side chain), the loop largely samples a bimodal distribution of open and closed states, with moderate population redistribution compared to wild-type SHP-1. In contrast, the T501M substitution destabilizes the closed WPD-loop, causing a substantive population shift toward a WPD-loop open ensemble. This is also shown in structural projections of the loop based on our simulations, which demonstrate that while wild-type and A323T sample broad conformational ensembles, T501M samples a much tighter pre-organized ensemble consistent with our tICA analysis. This is at least partially facilitated by the bulky methyl group introduced at position 501, which can sterically preclude WPD-loop closure (**Figure 5**). We further notice that this increase in open WPD-loop states is still associated with one dominant χ_2_ rotamer, whereas the A323T variant is able to sample, to a greater degree than T501M, both the t-90 and t-0 rotamer states in χ_2_ (**Figure 6C**).

In the case of SHP-2, kinetic measurements of the activity of the isolated catalytic domain of N308D and Q506P SHP-2 indicate a ∼2-fold increase in the *k*_cat_ of N308D full-length SHP-2 by alleviating autoinhibition (offset by a reduction in the activity of the isolated SHP-2 catalytic domain^26^), and a ∼10-fold decrease in *k*_cat_ of Q506P SHP-2, compared to wild-type.^26, 48, 94, 101, 102^ Comparison of WPD-loop dynamics in these variants (**Figure 7**) again indicates important differences in WPD-loop dynamics. In the case of the N308D variant, this substitution is located at the interface between the catalytic and N-SH2 domains, and has been suggested to disrupt autoinhibition, rendering the PTP constitutively active.^94, 99, 103, 104^ Given that the catalytic domain remains primarily exposed, it would be expected that differences in WPD-loop dynamics becomes a key driver of activity in this variant, consistent with other PTPs.^22, 105^ Our simulations (**Figure 7A**) indicate that while wild-type SHP-2 undergoes some bimodal sampling of open and closed states, the WPD-loop of N308D SHP-2 becomes trapped in in open conformation. This observation is important, because N308D SHP-2 has been demonstrated to be particularly sensitive to redox regulation, due to the formation of a disulfide bridge between the side chains of Cys367 and Cys359 in this variant (see ref. ^98^). An open WPD-loop would increase solvation of the active site and thus impact redox potential of the nucleophilic cysteine, as discussed in related prior work on ShufPTP.^106^

**Figure 7.**
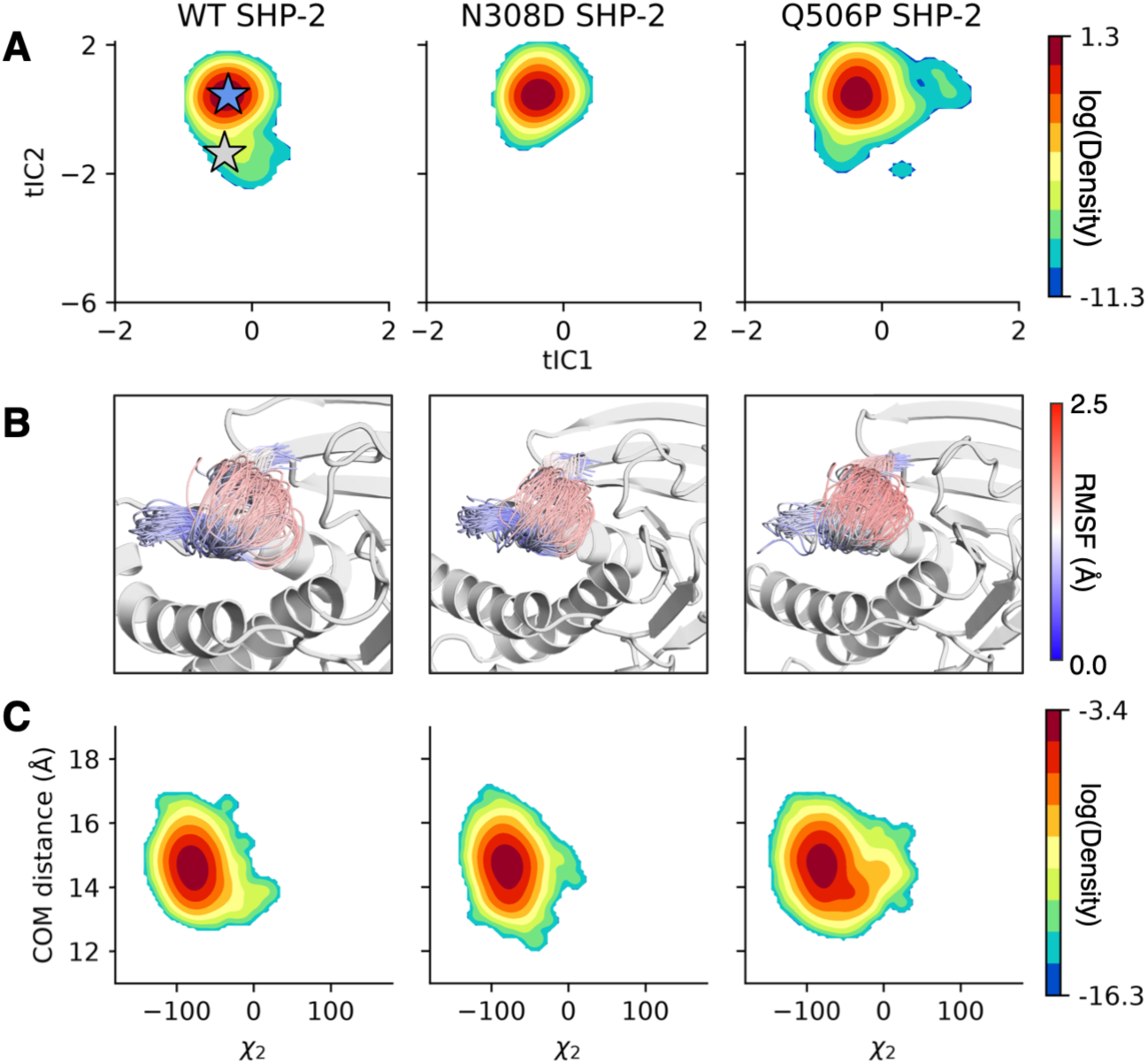
(**A**) Joint probability distributions of the first two time-independent components (tICs) generated from time-lagged independent component analysis (tICA) fitted on all pairwise C_α_-atom distances between the WPD-loop and P-loops and the sine and cosine values for the W417 (SHP-1 numbering) *χ*_1_ and *χ*_2_ side chain dihedral angles. The blue star indicates the open crystal structure, and gray star the closed crystal structure. (**B**) Projection of WPD-loop positions evenly spaced across the range center of mass (COM) distances between the WPD-loop and P-loops. Residues colored according to root mean squared fluctuation (RMSF) values. (**C)** Joint probability distribution of W417 (SHP-1 numbering) χ_2_ side chain dihedral angles *vs*. the COM distance between the WPD-loop and P-loop. See **Figure S4** for the corresponding probability distributions in the wild-type enzyme and **Figure S8** for the corresponding distributions of the χ_1_ side chain dihedrals in the variants. All results collected from 8 x 1.5µs molecular dynamics simulations initiated from the WPD-loop open unliganded state.

In contrast to the N308D substitution, which locks the conformational ensemble of the WPD-loop to open states, the Q506P substitution expands the conformational space of the loop, allowing it to access potentially non-catalytic conformations not observed in our simulations of wild-type or N308D SHP-2 (**Figure 7A**). Finally, we do observe subtly more sampling of the and t-0 rotamer states in χ_2_ for the W417 residue in the Q506 variant as opposed to the N308D variant. This is significant because, while Q506P sits at the interface between the N-SH2 and catalytic domains,^107^ biochemical characterization has suggested that this substitution impacts the substrate specificity of the active site rather than the autoinhibition mechanism,^104^ consistent with also the loss of intrinsic activity observed in the phosphatase/catalytic domain. The broadened WPD-loop ensemble observed in our simulations coupled with altered W417 rotamer preference (**Figure 7**) plausibly includes plausibly catalytically inactive loop conformations, supporting this kinetic observation. Importantly and overall, in both SHP-1 and SHP-2, our simulations indicate that substitutions distal to the active site but located on the allosteric pathways regulating WPD-loop motion have substantive impact on the ensemble of this key catalytic loop, further emphasizing that targeting WPD-loop closure provides a potentially effective strategy for inhibiting both SHP phosphatases.

### Modeling the Impact of Disease-Causing Mutations on the Energetics of WPD-Loop Closure

While our conventional MD simulations focus characterizing WPD-loop ensembles in the open/unliganded states of these phosphatases, an important question is how oncogenic mutations impact the energetics of WPD-loop closure, and the relative free energies of open and closed states. In prior work,^23^ we characterized these transitions in unliganded, Michaelis complex and phosphoenzyme intermediate states of PTP1B and YopH using bias-exchange metadynamics simulations.^108^ Here, we use Markovian milestoning with Voronoi tessellation (MMVT)^109^ as implemented into SEEKR2^110^ interfaced with OpenMM,^111^ to model WPD-loop closure in unliganded (Cys^−^) and phosphoenzyme intermediate (*p*Cys) states of wild-type and pathogenic SHP-1 and SHP-2 (**Figures 8** and **S9** to S**20**). Experimental benchmarks in related PTPs (PTP1B and YopH) established WPD-loop motion in these enzymes to be a μs-to ms-timescale process.^22^ Absolute transition state barriers obtained for complex conformational changes are sensitive to the choice of CVs. Here, we evaluate all systems using identical CV definitions, to isolate how specific disease-causing mutations modulate relative loop-gating energetics (ΔΔG), focusing on both activation barriers and thermodynamic shifts in loop equilibria.

**Figure 8.**
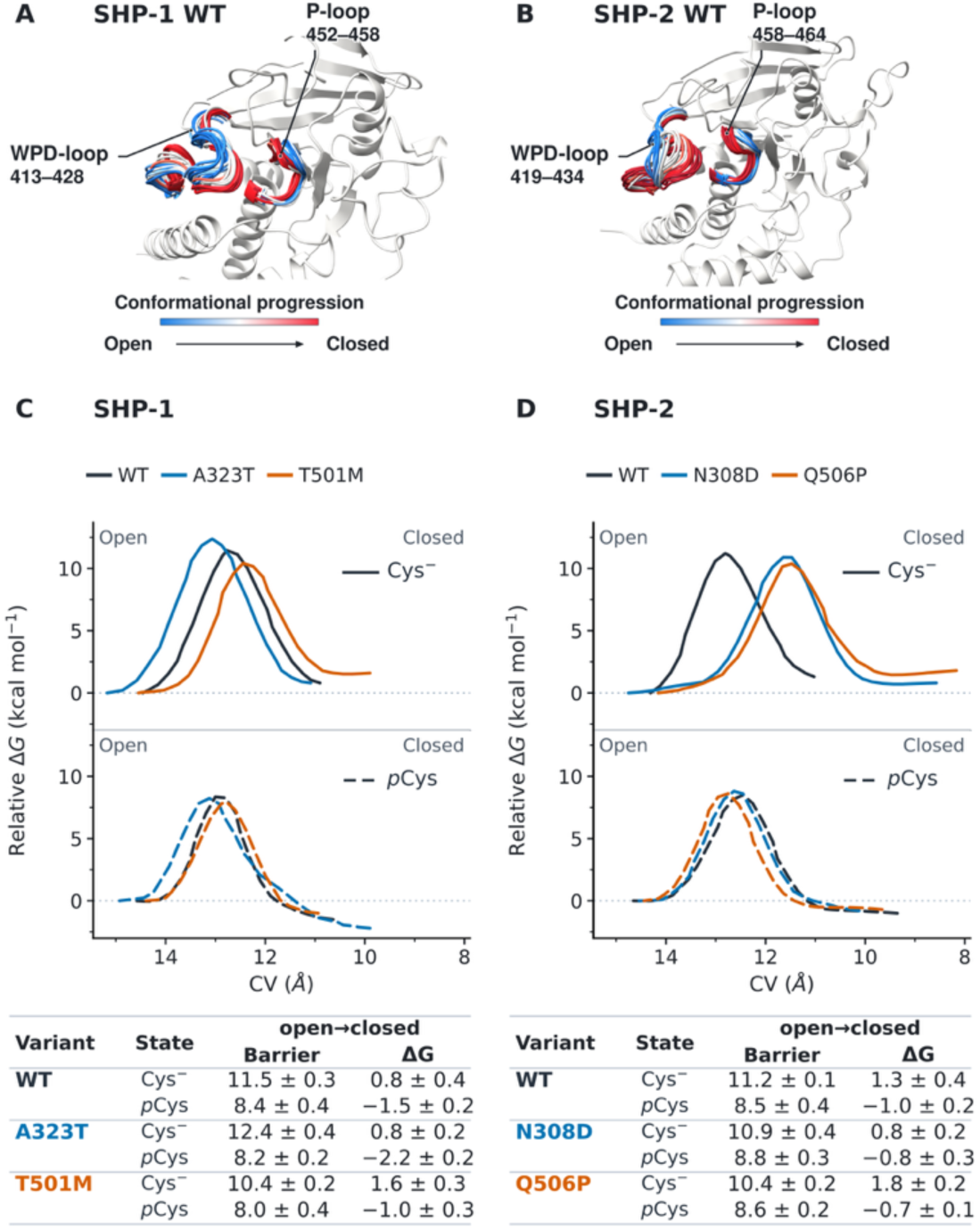
Representative conformations illustrating WPD-loop closure in wild-type **(A)** SHP-1 and **(B)** SHP-2. The WPD- and P-loops are labelled on the figure, and the blue-white-red gradient indicates the progression from open to closed states of the loops. Shown here as also free energy profiles for wild-type and variant **(C)** SHP-1 and **(D)** SHP-2 in the unliganded (Cys⁻; upper plots, solid lines) and phosphoenzyme intermediate (*p*Cys; lower plots, dashed lines) states, calculated using Markovian milestoning with Voronoi tessellation (MMVT)^109^ as implemented into SEEKR2^110^ interfaced with OpenMM.^111^ Wild-type profiles are shown in dark grey, A323T and N308D in blue and T501M and Q506P in orange. The collective variable (CV) used to define reaction progress is the distance between the WPD-loop and P-loop C_α_-atom groups, with decreasing values corresponding to loop closure. Each profile is referenced to its open state minimum. The associated tables in panel (**C**) give the closing barrier and the open-to-closed free energy difference (ΔG_open→closed_). Negative values indicate that the closed minimum is below the open minimum. All energies are in kcal mol⁻¹. Protein structures were visualized using UCSF ChimeraX (version 1.10.1).^112^

Our simulations (**Figure 8**) demonstrate that, for all systems, phosphorylation of the nucleophilic cysteine both reduces the barrier for WPD-loop closure by 1.8–4.2 kcal mol^−1^, while also shifting the thermodynamic ensemble of the loop towards a WPD-loop closed state, through a ligand-gated conformational change triggered by WPD-loop / phosphate interactions (see discussion in ref. ^23^). In SHP-1, T501M has the lowest reported free energy barrier for WPD-loop closure in both the unliganded and *p*Cys states, whereas A323T has the highest barrier in the unliganded state. In the *p*Cys intermediate, the closure barriers range from 8.0 to 8.4 kcal mol–1. Pairwise block-bootstrap comparisons (**Figures S21** and **S22**) found no statistically significant differences between either mutants or the wild-type after correction for multiple comparisons (A323T: *P*_adj_ = 1.000; T501M: *P*_adj_ = 0.145; **Figure S21**). Both mutations, however, significantly affect the reaction free energy for loop closure (A323T: *P*_adj_ < 0.001; T501M: *P*_adj_ = 0.0022; **Figure S22**), which impacts the population of the WPD-loop open and closed states. In the unliganded form of SHP-1, both wild-type and A323T SHP-1 slightly prefer the WPD-loop open state (0.7 kcal mol^-1^), with T501M further shifting the preference for an open loop to 1.6 kcal mol^−1^. In the phosphoenzyme intermediate state, all three variants now show a preference for a closed WPD-loop; however, A323T further stabilizes the closed state compared to wild-type (−2.2 *vs.* −1.5 kcal mol^−1^), whereas T501M reduces the preference of the closed conformation (−1.0 kcal mol^−1^). Thus, while these substitutions do impact loop closure kinetics in the unliganded SHP phosphatase, their primary effect is on the equilibrium between open and closed states of the loop.

In the case of SHP-2, both N308D and Q506P substitutions lower the kinetic barrier to WPD-loop closure in the unliganded enzyme, while very slightly increasing it in the phosphoenzyme intermediate state. Further, in unliganded SHP-2, the N308D and Q506P substitutions have the effect of shifting the equilibrium between the WPD-loop open and closed states in opposite directions (**Figure 8D**). Specifically, the N308D substitution reduces ΔG_open→closed_ from 1.3 kcal mol⁻¹ in the wild-type to 0.8 kcal mol⁻¹, whilst Q506P increases it to 1.8 kcal mol⁻¹, thereby increasing the preference for the open state (consistent with the reduced activity of this variant). In contrast, in the phosphoenzyme intermediate state, both variants very slightly destabilize the loop-open conformation.

These energetic trends are in qualitative agreement with the changes in WPD-loop conformational sampling that we have seen in our MD simulations of the unliganded catalytic domains (**Figures 6** and **7**). In SHP-1, the T501M mutant thermodynamically favors the open conformation which is in accordance with the shift in MD towards open WPD-loop states. A323T on the other hand has a lower thermodynamic barrier separating the two WPD-loop states, and continues to sample both conformations. In SHP-2, the Q506P mutant has a higher relative free energy for the closed conformation which is again consistent with the broadened ensemble of potentially non-productive loop conformations observed in the MD. In conjunction with the identification of these residues on the allosteric communication pathways (**Figure 5**), these findings link substitutions remote from the active site to changes in both the conformational landscape and energetics of the catalytic loop. Taken together, these calculations provide a quantitative framework for understanding the pathogenicity of these mutants, demonstrating the existence of a direct role for altered WPD-loop dynamics in aberrant enzyme activity in these variants.

### Empirical Valence Bond Calculations

Having established that disease-causing SHP phosphatase variants can alter WPD-loop dynamics, we address the question of whether these mutations also impact the chemical step of catalysis. To do so, we performed empirical valence bond (EVB) simulations^57^ on these variants following the same protocol and parameters as in our prior PTP studies,^29, 56, 59, 113^ in order to determine whether the cancer-causing amino acid substitutions affect the rate-limiting hydrolysis step of the PTP mechanism.^21, 30–33^ The resulting calculated reaction barriers are shown in **Figure 9** and **Table S4.** Further, representative structures of key stationary points from EVB simulations of wild-type SHP-1 and SHP-1 are presented in **Figures 9 and S23**.

**Figure 9.**
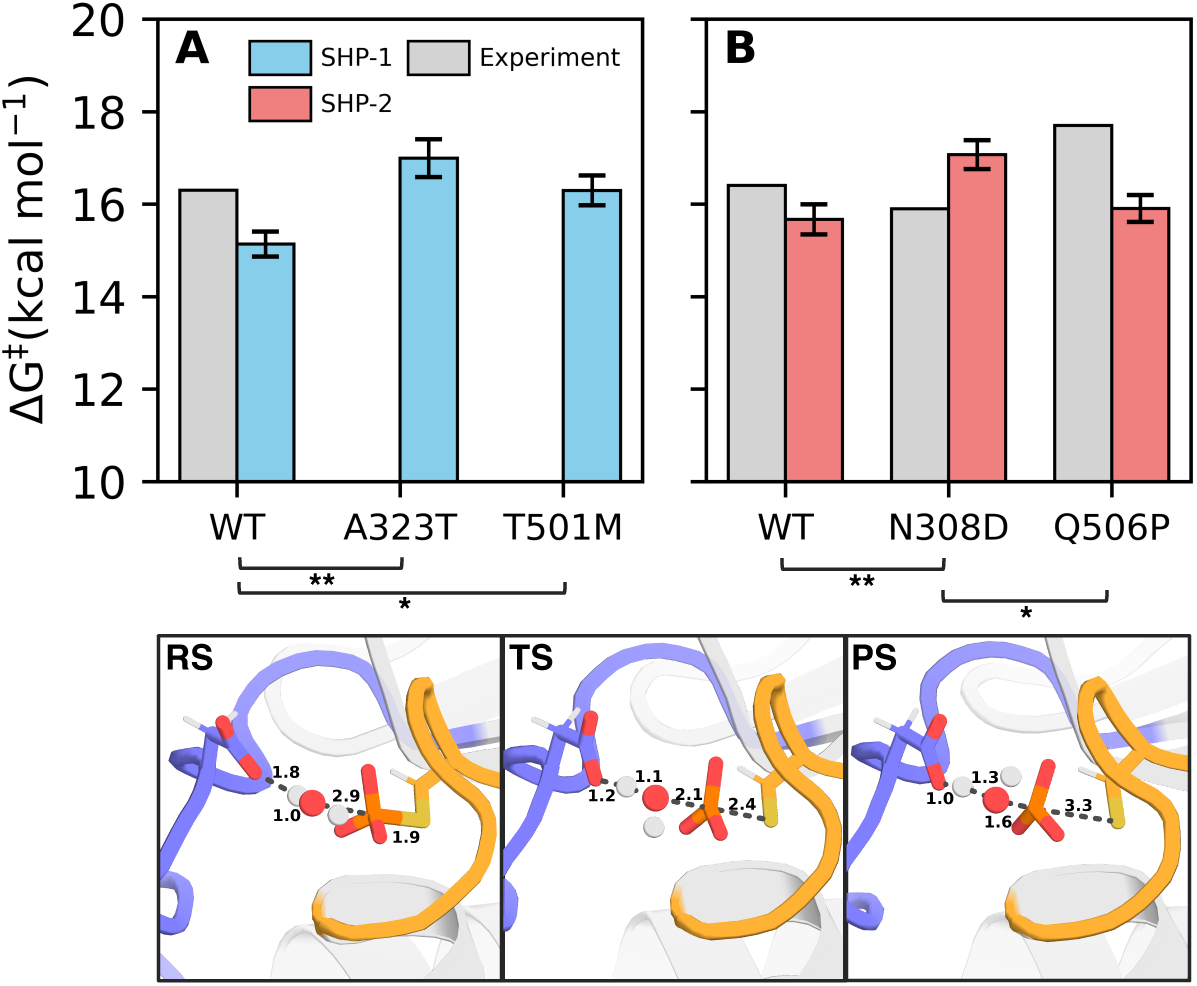
Comparison of experimental (gray) and calculated (blue/red) activation free energies (ΔG^‡^, kcal mol^-1^) for simulations of wild-type (WT) and variant forms of (**A**) SHP-1 and (**B**) SHP-2 initiated from structures extracted from molecular dynamics trajectories. Experimental values were derived from experimentally measured *k*_cat_ values, using transition state theory. The raw data for this figure, including the sources for the experimental data, are provided in **Table S4**. All calculated data are shown as average values and standard error of the mean over 30 independent empirical valence bond simulations. Stars indicate values which are statistically different (*p<0.05, **p<0.01) as calculated from a one-way ANOVA test followed by a Tukey Honestly Significant Difference (HSD) test.^114, 115^ Shown here are also representative reactant (RS), transition (TS) and product (PS) state structures from our EVB trajectories, extracted from the dominant cluster obtained from hierarchical agglomerative clustering on reacting heavy atoms during empirical valence bond (EVB)^57^ simulations of wild-type SHP-1, using CPPTRAJ.^69^ The equivalent structures for SHP-2 are shown in **Figure S23** and the corresponding reactive distances are shown in **Table S5.**

Based on these data, it can be seen that, in line with our prior EVB simulations of a range of PTPs including SHP-1,^23, 29, 56, 59^ our EVB calculations of wild-type SHP-1 and SHP-2 are in good agreement with the corresponding experimental values,^21, 30, 48, 94, 116–118^ especially given that it is plausible that WPD-loop motion is at least partially rate-limiting as in PTP1B.^22, 23^ In the case of SHP-1, we see an increase in activation free energy upon introduction of the A323T (p<0.01) and p<0.05) variants, in agreement with a reduction in reactive conformations upon disrupting the dynamics of the WPD-loop (**Figure S24**). This disruption is more pronounced in A232T than in T501M SHP-1, following the trend in activation free energies, although we note that all four SHP phosphatase variants considered in this work reduce the % of reactive conformations observed in our simulations. In the case of SHP-2, we again observe an activation free energy increase in the N308D variant (p<0.01). We note that this contradicts kinetic measurements of the intrinsic activity N308D SHP-2 catalytic domain measured using a peptide substrate , which suggest a slight *increase* in activity in this variant.^94^ However, more recent deep scanning mutagenesis data using the fluorogenic substrate 6,8-difluoro-4-methylumbelliferyl phosphate (DiFMUP) has suggested that mutations at position N308 in SHP-2 decrease intrinsic phosphatase activity, in line with our EVB calculations.^26^ Finally, we observe no notable difference in activation barrier between the SHP-2 wild-type and Q506P variant (p>0.05), indicating that experimentally observed loss of activity^48, 94^ is likely driven by altered loop dynamics rather than a chemical effect. Statistical significance was calculated using a one-way ANOVA test, followed by a Tukey Honestly Significant Difference (HSD) test.^114, 115^

Overall, these observations are in agreement with prior work,^29, 56, 59, 113^ where calculated activation energy differences between systems were small, as is to be expected given the nearly superimposable active sites of these enzymes. However, as shown in **Figure S24**, these mutations disrupt the fraction of reactive conformations sampled by these enzymes, which is in turn catalytically disruptive. Taken together, these data indicate that as in our prior studies,^29, 56, 59, 113^ effects on the chemical steps can be negligible compared to the impact of loop motion on catalysis, due to the essential role of catalytic loop of PTPs in positioning a key catalytic residue in the PTP active site (**Figure 1**).

## Conclusions

The protein tyrosine phosphatases SHP-1 and SHP-2 are attractive but challenging drug targets for the treatment of cancer and a number of rare diseases,^10, 36, 38, 40, 41^ making them important topics of study. These enzymes have evolved as an evolutionary pair,^19^ and are the only known PTPs to carry tandem SH2 domains (**Figure 1**),^44^ which are involved in the autoinhibition of both SHP PTPs.^27, 28, 45–47^ Given the biomedical importance of these enzymes, the dynamics of SHP PTPs have been the subject of significant computational studies.^24, 72–83, 119–121^ However, the majority of these have focused largely on the autoinhibition of these SHPs rather than the regulation of WPD-loop motion, despite the intimate link between loop dynamics and chemistry in PTPs.^22, 23^ In particular, a landmark study of by Fortin and coworkers showed that allosteric inhibition of SHP-2 by an orally bioavailable small molecule inhibitor, SHP099, stabilizes SHP-2 in an autoinhibited conformation and inhibits cancers driven by receptor protein kinases.^122^ Recent work, has, however, showcased that oncogenic mutations and allosteric drug inhibition impact WPD-loop motion in SHP-2.^26, 84^ We present here a detailed computational study of truncated catalytic domain and full length SHP-1 and SHP-2 (both wild-type enzymes and selected oncogenic variants), focusing on the impact of the SH2 domains and on oncogenic substitutions on WPD-loop dynamics and chemistry.

Our simulations demonstrate that despite the high sequence identity of (60% across catalytic domains) and active site structure conservation between SHP-1 and SHP-2,^14^ these enzymes exhibit fundamentally distinct WPD-loop dynamics in their catalytic domains. Specifically, unliganded SHP-1 is able to sample intermediate and closed states of the WPD-loop even when initiating simulations from a WPD-loop open conformation, unliganded SHP-2 remains tightly localized in a WPD-loop open state, and PTP1B acts as a dynamical intermediate between the two SHP phosphatases (**Figure 2**). As SHP-1 and SHP-2 are enzymes that evolved as a pair through gene duplication,^19^ these clear differences in dynamics showcase how differences in sequence can allow for dramatically different function.

Further, our simulations of full-length active SHP phosphatases indicates that the presence of the tandem SH2 domains impacts WPD-loop dynamics in both enzymes. However, this happens in an orthogonal fashion in the two enzymes specifically, our simulations show that in full-length SHP-2, the SH2 domains expand the conformational ensemble of the WPD-loop and associated rotamer sampling of the WPD-loop Trp side chain compared to the isolated catalytic domain (**Figure 4**). In contrast, in SHP-1, the presence of the SH2 domains narrows the WPD-loop conformational ensemble, focusing sampling on the WPD-loop open state, while still allowing for some sampling of a more closed state (**Figure 4**). This is consistent with experimental kinetic observations demonstrating a much larger impact of autoinhibition on *k*_cat_ in SHP-2 than in SHP-1, whereas in SHP-1 the impact of the SH2 domains is primarily on *K*_M_ rather than *k*_cat_.^30, 48^ We also note that our simulations indicate that full-length SHP-2 exhibits much larger spatial mobility of the N-SH2 domain relative to the catalytic core of this PTP compared to SHP-1 (**Figure S5**).

Following from this, shortest path map (SPM) calculations reveal a conserved allosteric network across SHP-1, SHP-2 and PTP1B (up to 50% conservation of allosteric positions), consistent with prior work on PTPs more broadly (**Figure 5**).^23, 92, 93^ We further observe high congruence between SPM-predicted residues and oncogenic/disease associated mutations listed in the COSMIC database,^96^ the majority of which either lie directly on or are within <4Å of the proposed allosteric networks. This links these distal disease-causing mutations to regulation of catalytic loop motion. To validate this, we considered in detail four known disease-causing variants of these enzymes: T501M/A323T in SHP-1, and N308D/Q506P in SHP-2.^38, 48, 96, 97^ Conventional MD (**Figure 6**) and Markovian milestoning with Voronoi tessellation (MMVT, (**Figure 8**))^109^ simulations show that these disease-causing mutations (**Figure S6**) both shift the conformational ensemble of the WPD-loop and fine-tune the free-energy landscape associated with this motion. Despite being distal to the active site, these mutations also disruptive active site dynamics, leading to reduced sampling of catalytically productive conformations (**Figure S24**), while the activation free energies for rate-limiting phosphoenzyme hydrolysis^21, 30–33^ remain largely conserved across variants (**Figure 9**). This shows that the mutations are primarily impacting WPD-loop motion and active site preorganization rather than the intrinsic chemical reactivity of these enzymes.

Taken together, our computational work provides detailed molecular insight into catalytic loop motion in these biomedically important PTPs, and highlights key differences between the dynamical profiles of the two enzymes. Prior work has emphasized the importance of the flexibility of the SH2 domains of these SHP phosphatases,^28, 42, 86^ the importance of the autoinhibition mechanism,^27, 28, 45–47^ the ability of disease causing mutations to shift the equilibrium between inactive and active states of SHP-2 with respect to its SH2 domains towards an inactive conformation,^71^ and also highlighted the role of oxidative regulation in the case of N308D SHP-2.^98^ Our data demonstrate that, while allosteric regulation of the autoinhibition mechanism is an attractive drug target in these enzymes (see *e.g.*, refs. ^122–127^, among others) WPD-loop motion is also important, and our data suggest that both SHP-1 and SHP-2 can likely be targeted by allosteric inhibitors focused on blocking WPD-loop closure, as has been achieved with other PTPs.^24–26^ That is, correct function of these SHP phosphatases is a multi-layer process, involving (1) shifting the equilibration of the full complex towards formation of an active state, (2) ensuring stable association of the SH2 domains with the catalytic domain,^71^ and ultimately (3) regulation of WPD-loop motion, which facilitates the actual chemical step of catalysis. If any of these steps are disrupted, this in turn has disease-causing potential. Given that the impact of the SH2 domains on WPD-loop motion and catalysis differs between the two PTPs, this expands our repertoire of strategies to develop isoform selective inhibitors for these critical but elusive anti-cancer drug targets.

## Materials and Methods

### Structure Selection and Preparation

While there exist structures of SHP-1 in both catalytically active and autoinhibited conformations of the SH2 domains (PDB IDs: 3PS5^27^ and 2B3O,^28^ respectively), the structure of full-length SHP-2 has only been solved in its autoinhibited conformation (PDB ID: 4DGP^47^). Historically, the structure of the constitutively active E76K variant has been used as a model for active SHP-2.^47, 128^ However, there have been concerns raised in the literature about how realistic a model this is for active SHP-2,^76, 129^ and NMR studies have shown that the E76K variant samples an atypical open state of SHP-2.^84^ Further, given the focus of this work on the conformational ensemble of the WPD-loop, it is plausible that E76K would alter this ensemble compared to the wild-type counterpart. To address this issue, we leveraged BioEmu^85^ to generate plausible Boltzmann ensembles of WPD-loop open SHP-1 and SHP-2, seeding from the corresponding autoinhibited structures, (PDB IDs: 2B3O^28^ for SHP-1, and 4DGP^102^ for SHP-2). We then extracted an SH2-open active structure from the ensembles to use as a starting point of our simulations. While an active state crystal structure exists for SHP-1 (PDB ID: 3PS5^27^), using BioEmu^85^ to generate starting structures for both PTPs allows for consistency in our simulations.

Conformational sampling was performed using BioEmu version 1.3.1, *via* its Python sampling interface.^85^ Side chains were reconstructed and relaxed using the BioEmu sidechain_relax module, with its molecular dynamics equilibration step enabled. Individual conformations were extracted from the reconstructed trajectories in PDB format, using CPPTRAJ^69^ for the subsequent structural comparison. Once we obtained reasonable conformational ensembles, we determined the structure most likely to be in a biologically relevant active state by extracting the structure which exhibited the lowest RMSD to the active SHP-1 crystal structure (PDB ID: 3PS5^27^). An alignment of the SHP-1 BioEmu structure used for our simulations and the corresponding available crystal structure is shown in **Figure S25**. Finally, we note that there exist variations in residue numbering for SHP-1 based on different interpretations of sequence.^29, 116, 130, 131^ Throughout this work, we use the residue numbering used in the COSMIC database,^96^ among other sources, for consistency.

Following from this, there exist a number of crystal structures of the SHP-1 catalytic domain with alternate positions of the N-terminal helix (see PDB IDs 4HJP,^130^ 4GRZ^131^ and 8YHI^132^). In full-length SHP-1, this helix bridges the C-SH2 and PTP domains,^28^ and has been suggested to play a cooperative role in substrate recognition.^133^ An analogous terminal helix in PTP1B is ordered in the WPD-loop closed conformation of the PTP, and disordered in open/inhibited conformations of the enzyme.^25, 65, 67, 134^ Since it was unclear which helix position was preferable for catalysis, we conducted benchmarks to determine the impact of helix position on WPD-loop motion. We used the Morph2 webserver (https://www.bioinformatics.org/pdbtools/morph2) to construct twelve SHP-1 starting structures, smoothly varying the starting position of the N-terminal helix for SHP-1 in both the closed WPD-loop state (PDB ID: 4GRZ^131^) and in the open WPD-loop state (PDB ID: 4HJP^130^) (24 simulations total), which we ran using the same equilibration procedure described below each followed by 500ns of production MD using GPU-accelerated GROMACS version 2024.2. Results indicated that helix positions located further from the main scaffold (such as those observed in the 4HJP^130^ and 8YHI^132^ structures) can be unstable and can affect WPD-loop motion *via* disrupted active site residue interactions, as the highly mobile helix forces the WPD-loop shut *via* interactions among a selection of N-terminal helix Glu residues and the active site residues. This motion, in turn, deforms the structure of the N-terminal helix. (**Figure S26)**. Based on this observation, we grafted the starting coordinates of the α_0_-helix in the 4GRZ^131^, which is closer to the protein scaffold, onto the 4HJP^130^ open-loop structure for our simulations. We note that throughout our simulations, this helix samples a broad distribution of conformations, in contrast to the predominant localized conformation of the corresponding helix in SHP-2, accompanied by minor sampling of conformations further from the core scaffold (**Figure S26**).

SHP-2 catalytic domain only simulations were performed using PDB ID: 4RDD,^135^ as this is the highest resolution (1.6Å) available structure, with the cumulative best quality scores. However, as this structure lacks density for the N-terminal helix, this helix was grafted onto the scaffold using coordinates from PDB ID: 6CMQ,^84^ and removing any clashing water molecules. All other missing regions were fixed by using the Modeller^136^ extension of ChimeraX,^112^ generating 30 possible structures, and taking the structure with the lowest energy score for further simulations. In each unliganded PTP, the catalytic Asp and Cys residues (**Figure 1**) were kept in their deprotonated forms respectively. Protons were added to the structures using MolProbity,^137^ and PROPKA 3.1^138^ was utilized to identify any anomalous protonation states of ionizable residues, followed by extensive visual inspection. Based on this analysis, all other residues except the catalytic cysteine in unliganded PTPs were kept in their standard protonation states at physiological pH. We finally solvated each protein in a truncated octahedral water box extending 10Å away from the protein in all directions, and neutralized our systems with. Na^+^/Cl^-^ counterions. Parameters describing the phosphorylated cysteine along with all starting structures of PTP1B were taken from prior work.^23, 113^ For all oncogenic variants, amino acid substitutions were made in PyMOL^139^ using the Dunbrack rotamer library,^62^ and the best rotamers were chosen from visual inspection of possible clashes. Residues 313-315 in PDB ID: 4HJP^130^ (SHP-1 catalytic domain, WPD-loop open) were manually added using PDB ID: 4GRZ^131^ (SHP-1 in complex with PO_4_, WPD -loop closed) as a template.

### Molecular Dynamics Simulations

All conventional molecular dynamics simulations were performed using GPU-accelerated GROMACS^140^ version 2026.1. In all cases, the AMBER ff14SB^141^ forcefield was used along with the TIP3P water model,^142^ and simulation starting structures were generated using AmberTools22,^143^ utilizing the tLEaP and ParmEd libraries. A table of crystal structures used in this paper is shown in **Table S6**. Simulations were performed in the unliganded forms of each protein. Simulations were performed using the same protocol as described in prior work.^29^ In brief, the system was heated to 300K over 50ns using velocity rescaling,^144^ followed by 50ns NPT production at 300K and 1 atm using a Parinello-Rahman barostat^145^ before initiating 1.5 μs of production simulations in 8 replicas for each of wild-type SHP-1, SHP-2, oncogenic variants, and PTP1B. All simulations were performed in each of the WPD-loop open unliganded state. This led to a total cumulative simulation time of 216 μs across all systems. Simulation convergence is shown in **Figures S27** – **S30.** Additional simulation details are found in ref. ^29^.

Simulation analyses were carried out using CPPTRAJ^69^ for creating dynamic cross-correlation matrices (DCCM), deeptime for tICA analysis,^146^ and MDAnalysis^147^ for all other analyses. We utilized the EMBL-EBI Job Dispatcher^148^ to conduct sequence alignments using MUSCLE,^58^ and used the Shortest Path Map method^91^ in order to compute allosteric pathways. All protein structures presented in this manuscript, unless otherwise noted, were visualized using PyMOL.^139^

### Markovian Milestoning Calculations of WPD-Loop Motion

The WPD-loop opening and closing were studied using Markovian milestoning with Voronoi tessellations (MMVT),^109^ as carried out in SEEKR2^110^ in conjunction with OpenMM.^111^ The calculations were carried out for wild-type SHP-1 (full-length and catalytic domain) and its A323T and T501M mutants, and for wild-type SHP-2 (full-length and catalytic domain) and its N308D and Q506P mutants, in both the unliganded cysteine (Cys^−^) and phosphocysteine intermediate (*p*Cys) states. MMVT is able to reconstruct global thermodynamic and kinetic properties from molecular dynamics simulations that are carried out within each Voronoi cell.^109, 110, 149^

The loop transition was described by means of a one-dimensional collective variable (CV) based on the pairwise distance between the groups of C_α_-atoms located at the centers of the WPD-loop and the P-loop. For SHP-1, the WPD-loop comprised residues 413−428 and the P-loop residues 452−458; for SHP-2, the WPD-loop comprised residues 419−434 and the P-loop residues 458−464. These residue ranges refer to the numbering used in the simulation models. Smaller coordinate values reflect closed-like conformations, whereas larger values indicate open conformations. The same definitions of the residue groups were used consistently in all the variants and chemical states of each protein.

The initial configurations covering the transition were obtained from steered molecular dynamics (sMD) trajectories performed using GROMACS 2025^140^ patched with Plumed 2.10.^150^ Each system followed a standard minimization as described in the section details for our molecular dynamics systems and shortened equilibration procedure (125ps of heating, followed by three stages of 1ns equilibration, before 50ns of steering). Each structure was initiated from the WPD-loop open starting structure and steered closed by applying a time-dependent moving harmonic restraint, targeting a dRMSD of 0 with respect to the PTP1B closed WPD-loop crystal structure, including all C_α_-atoms of the WPD-loop and P-loop, and the C_γ_-atom of the general acid (D181 in PTP1B numbering). To steer slowly, the *κ* value was ramped from 0 to 3 kcal^−1^ mol/Å^−2^ over the course of the steering. Anchors were chosen at intervals of 0.3 Å along the CV. The anchors determined the Voronoi cells, and the common boundaries between adjacent cells formed the milestones. The positions of the physical milestones that were recorded in the SEEKR2^110^ models generated were then used in the later analysis.

The anchor structures were created in explicit TIP3P water molecules,^142^ with counterions added to neutralize each system. Additional NaCl was introduced at a nominal concentration of 0.15 mol L⁻¹ to approximate physiological salt conditions. Na⁺ and Cl⁻ were described using the TIP3P^142^ – compatible ion parameters of Joung and Cheatham,^151^ as distributed with the AMBER force field used in the OpenMM simulations (AMBER ff14SB^141^). *p*Cys parameters were taken from prior work.^23^ The MMVT simulations were carried out in the canonical ensemble at 303.15 K, with a time step of 2 fs and a nonbonded cutoff of 1.0 nm. The production run consisted of 5 × 10^6^ steps, which amounted to 10 ns per anchor, and the coordinates were saved every 2 ps. Reflecting boundary conditions kept each trajectory within its designated cell, and the times at which boundary encounters occurred, together with the encounters themselves, were recorded for use in the milestoning analysis.

The first 25% of the MMVT sampling of each anchor was discarded as equilibration; only the last 75% was kept for the calculation of the transition statistics, the residence times, the equilibrium populations, and the resulting free-energy and kinetic quantities. The cutoff used for equilibration was applied individually to each anchor, and both the number of events and the total sampling time were assessed during the retained period.

The event statistics obtained after equilibration were analyzed using the standard SEEKR2 MMVT reconstruction method, in order to determine the equilibrium populations and the transition rates between milestones.^152–155^ The relative free energies were calculated from the normalized equilibrium populations, *P_i_*, according to:

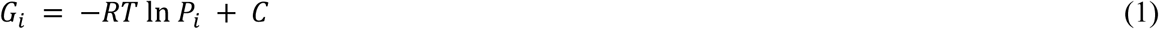

where *R* is the ideal gas constant, *T* is the simulation temperature, and *C* is an arbitrary reference constant. The relative stability of the open and closed states was expressed as

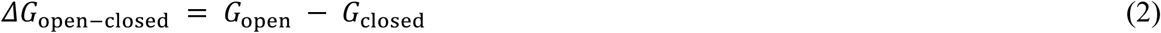

The apparent closing barrier was determined from the free-energy maximum separating the two basins relative to the open-state minimum.

### Empirical Valence Bond Simulations

We performed empirical valence bond (EVB)^57^ simulations of WT SHP-1, SHP-2, and oncogenic variants at the phosphoenzyme intermediate state, since the corresponding hydrolysis reaction is expected to be the rate-limiting step of this phosphatase-mediated reaction.^21, 30–33^ All simulations were performed starting from snapshots taken from molecular dynamics simulations initiated at the WPD-closed conformation (catalytic domain only) conformation.

For the initial molecular dynamics simulations, the 6CMQ^84^ closed conformation was used for SHP-2, removing the N-SH2 domain resolved in that structure and the 4GRZ^131^ structure used for SHP-1, following the same setup outlined above, using GROMACS version 2022.5.^140^ Note that the differences in GROMACS version are due to independent simulations performed across distinct high-performance computing (HPC) facilities, utilizing architecture optimized GROMACS builds. For each PTP, the system was heated to 300K over 100ps using velocity rescaling,^144^ followed by 100 ps NPT production at 300K and 1 atm using a Parinello-Rahman barostat.^145^ Following this initial equilibration, we performed 100ns of further NPT equilibration before initiating 1.5 μs of production simulations per trajectory. Each system was simulated in 8 replicas for each of wild-type SHP-1, SHP-2, and oncogenic variants.

Snapshots were considered productive if the Asp419(O)-pCys453(P) distance was less than 4.5Å and the Gln500(C)-pCys453(P) distance (SHP-1 numbering) was less than 8Å to ensure the Gln was able to point towards the active site and stabilize the catalytic water molecule (**Figure S24**). All system setup, equilibration and subsequent EVB simulations were performed as described in prior work,^29, 56, 59, 113^ using also the same EVB parameters as for prior studies. All EVB simulations were performed using the OPLS-AA force field.^156^ Each system was solvated in a 23 Å radius sphere of TIP3P water molecules^142^ centered on the phosphate atom of the phosphorylated cysteine, and all residues within the initial 85% of radius from the center of the sphere were ionized following the same procedure as for our standard molecular dynamics simulations. A list of ionization states used for our EVB simulations is presented in **Table S7**. This solvent sphere was described using the Surface Constrained All Atom Solvent Model (SCAAS),^157^ and long-range effects were described using the local reaction field approach.^158^ In each system, the nucleophilic water molecule as placed manually in the active site, aligned in an optimal position for catalysis relative to both nucleophilic attack on the phosphorus atom and deprotonation by the aspartic acid side chain. An identical procedure was used for all systems. Following 35 ns of initial equilibration, the convergence of which is shown in **Figures S31** and **S32**, each EVB trajectory was propagated in 51 EVB mapping windows (*λ*_m_) of 200 ps in length each, with 30 independent EVB trajectories propagated per system. This resulted in 10.2 ns of EVB simulation time per trajectory, 306 ns per system, and 1.8 μs cumulative simulation time across all systems studied in this work.

## Supporting Information

Additional analysis of conformational flexibility, simulation convergence, experimental and calculated activation free energies for EVB simulations, PDB structures used in this work, and ionization states of key residues in our EVB simulations are presented as Supporting Information. All raw data necessary to reproduce our MD, sMD, MMVT and EVB simulations have been uploaded to Zenodo, DOI: 10.5281/zenodo.18817309.

## Acknowledgements

This work was supported by the Swedish Research Council (VR 2025-03686). A.-L. R. B. was supported by InQuBATE grant NIH T32GM142616. We acknowledge the National Academic Infrastructure for Supercomputing in Sweden (NAISS), partially funded by the Swedish Research Council through grant agreement no. 2022-06725, for awarding this project access to the LUMI supercomputer, owned by the EuroHPC Joint Undertaking and hosted by CSC (Finland) and the LUMI consortium. Additional simulations were enabled by resources provided by the National Academic Infrastructure for Supercomputing in Sweden (NAISS), partially funded by the Swedish Research Council through grant agreement no. 2022-06725. We further acknowledge the Georgia Institute of Technology PACE supercomputing resources, as well as the Purdue University Anvil and NCSA Delta supercomputers through allocation BIO250146 from the Advanced Cyberinfrastructure Coordination Ecosystem : Services & Support (ACCESS) program. Finally, we thank Alvan Hengge, Sean Johnson and Patrick Loria for helpful discussion.

---

Table of Contents Graphic

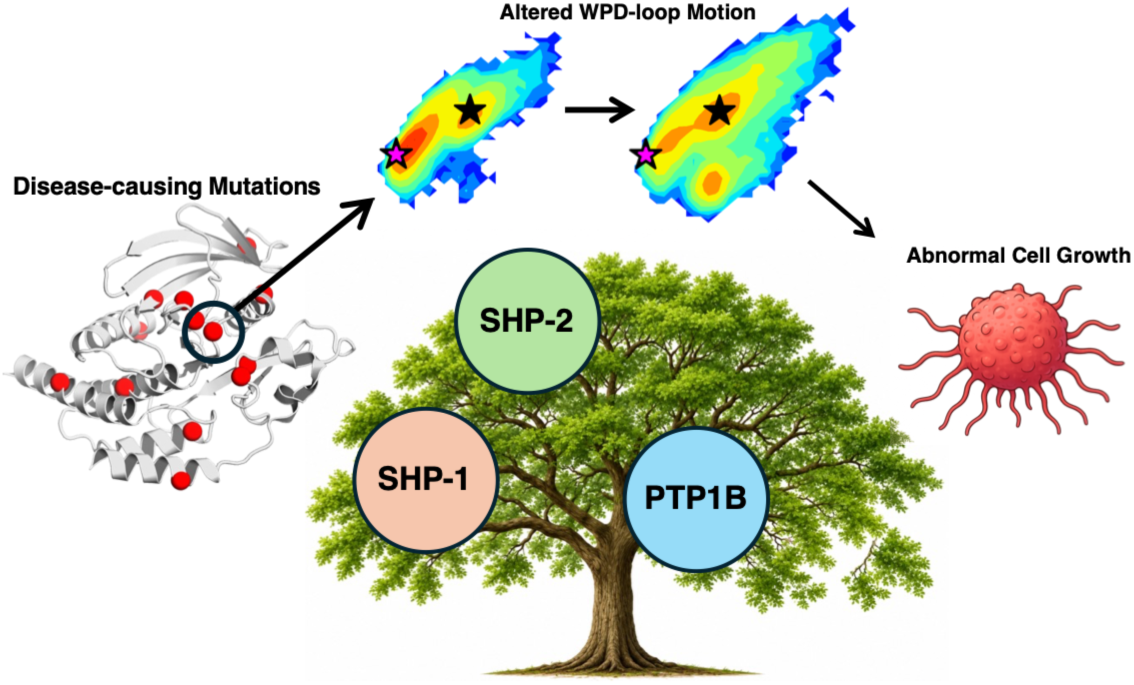

