## Supporting Information for "Disease-Causing Mutations Alter the Interplay Between Loop Dynamics and Catalysis in the Protein Tyrosine Phosphatases SHP-1 and SHP-2"

### Table of Contents

Supplementary Figures ........................................................................................................ S3

Supplementary Tables ........................................................................................................ S47

Supplementary References ........................................................................................................ S54

### Supplementary Figures

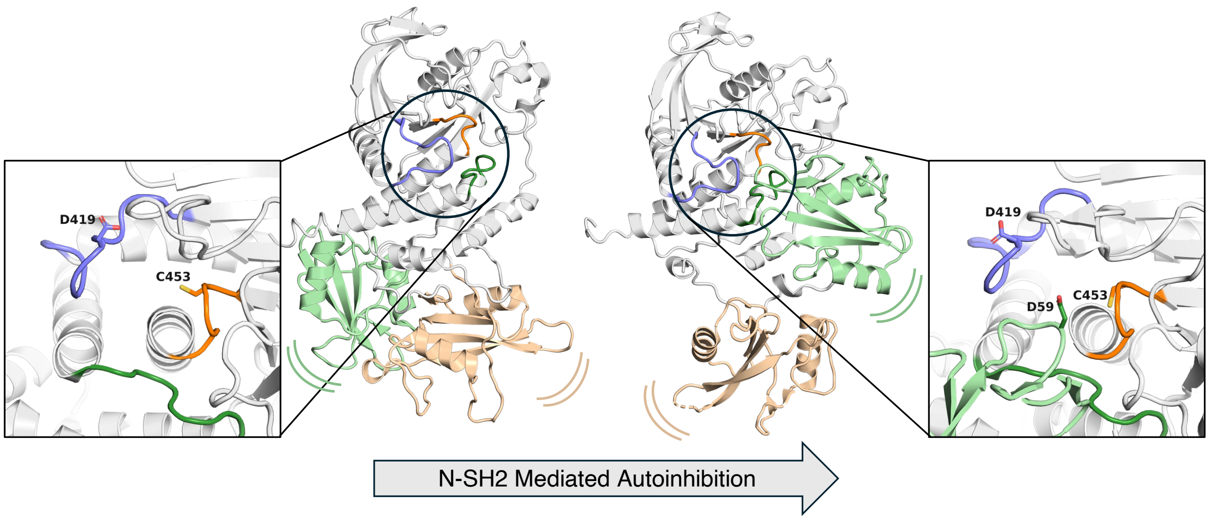


**Figure S1.** Visualization of N-SH2 domain autoinhibition, which is observed in both SHP-1 and SHP-2. In the non-autoinhibited state (left), the WPD-loop (blue) can move towards the P-loop (orange), resulting in a catalytically active conformation. By contrast, when autoinhibited (right), the N-SH2 domain (light green) interacts with the active site, preventing complete closure of the WPD-loop and thus preventing a catalytically active conformation from forming. The Q-loop is shown in dark green, and the C-SH2 domain is shown in tan. Note that structures shown are of SHP-1 in the open-SH2 state (PDB ID: 3PS5[1]), and in autoinhibited state (PDB ID: 2B3O[2]).

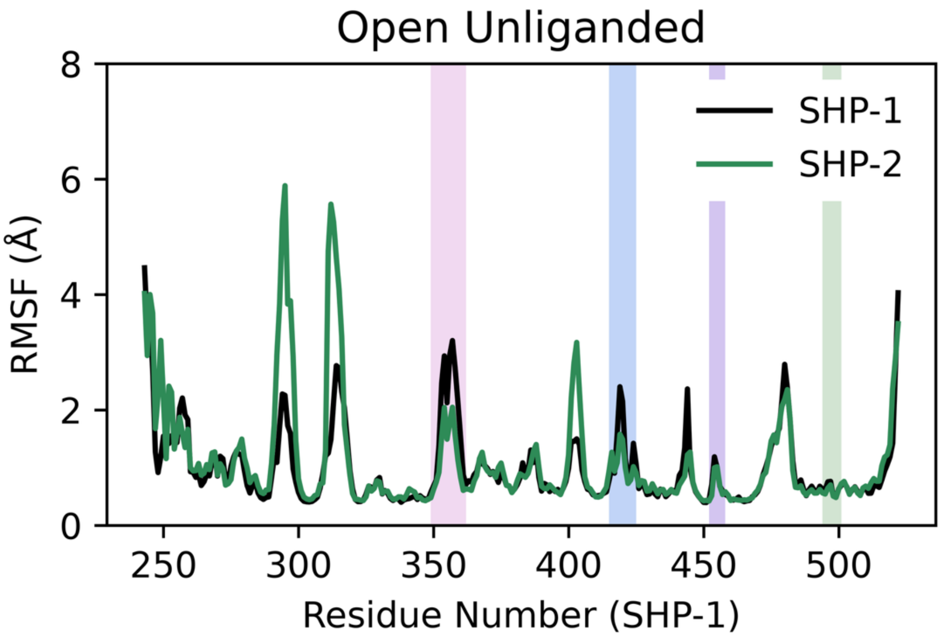


**Figure S2.** Root mean square fluctuations (RMSF, Å) of all Cα-atoms of wild-type SHP-1 and SHP-2 calculated across 8 × 1.5 μs concatenated MD trajectories. Regions of interest were aligned using MUSCLE.[3] Shading indicates key active site loop regions as shown in **Figure 1**: E-loop (pink), WPD-loop (blue), P-loop (purple), Q-loop (green).

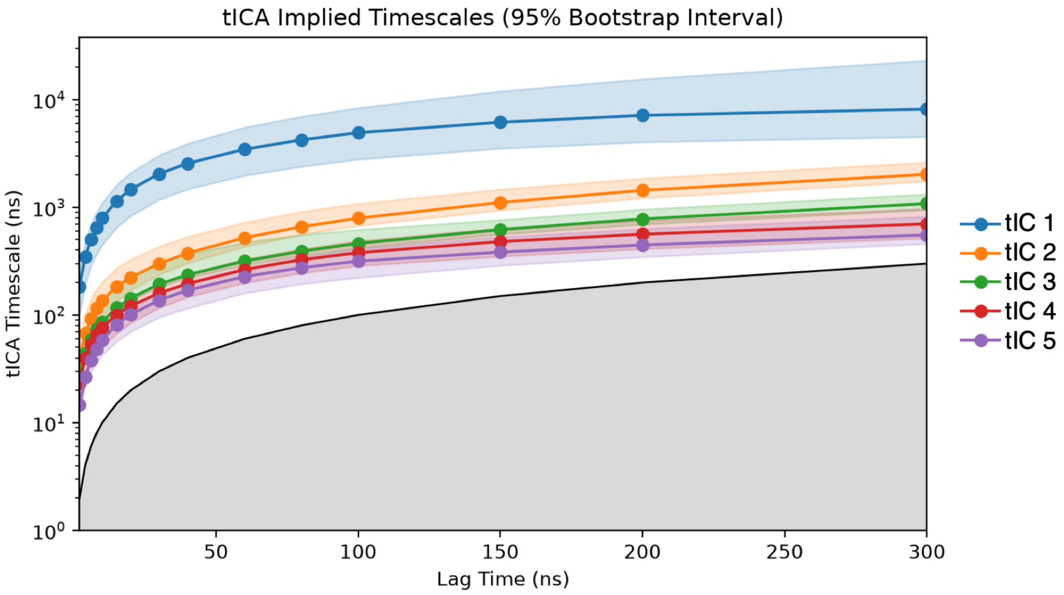


**Figure S3.** Implied timescales of tICA components. Original features included all distances between WPD-loop and P-loop $C_\alpha$-atoms, and sine and cosine values for the W417 (SHP-1 numbering) $\chi_1$ and $\chi_2$ dihedrals collected across 8 x 1.5µs replicas of molecular dynamics simulations for all systems presented in this work at the unliganded open WPD-loop state (see **Materials and Methods**). We note that tICA analysis is being used in this work as a dimensionality reduction technique for conformational analysis, and not for extracting kinetic rate constants. While tIC1 reaches a stationary plateau around $\tau$ = 150ns; a shorter lag time of 20ns was selected for tICA space projection in order to capture sub-state heterogeneity and resolve intermediate states during WPD-loop motion.

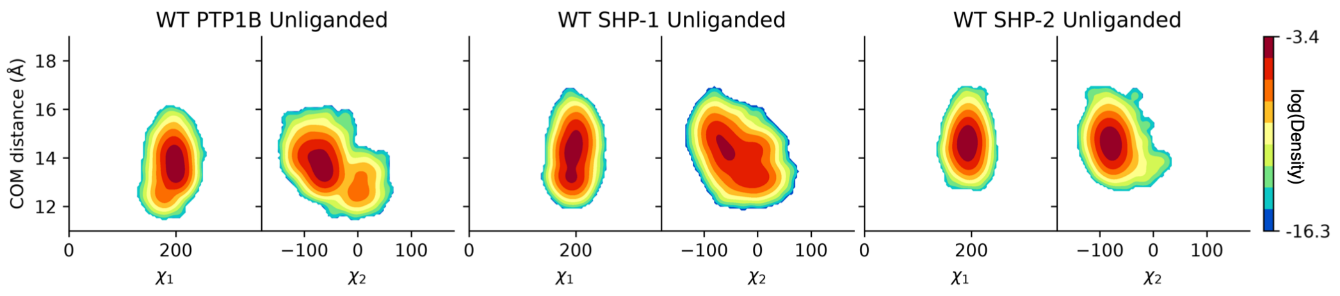


**Figure S4.** Joint probability distribution of W417 (SHP-1 numbering) $\chi_1$ and $\chi_2$ dihedral angles *vs*. the center of mass (COM) distance between the WPD-loop and P-loop. Results collected over 8 x 1.5μs molecular dynamics simulations of PTP1B, and the catalytic domains of SHP-1 and SHP-2.

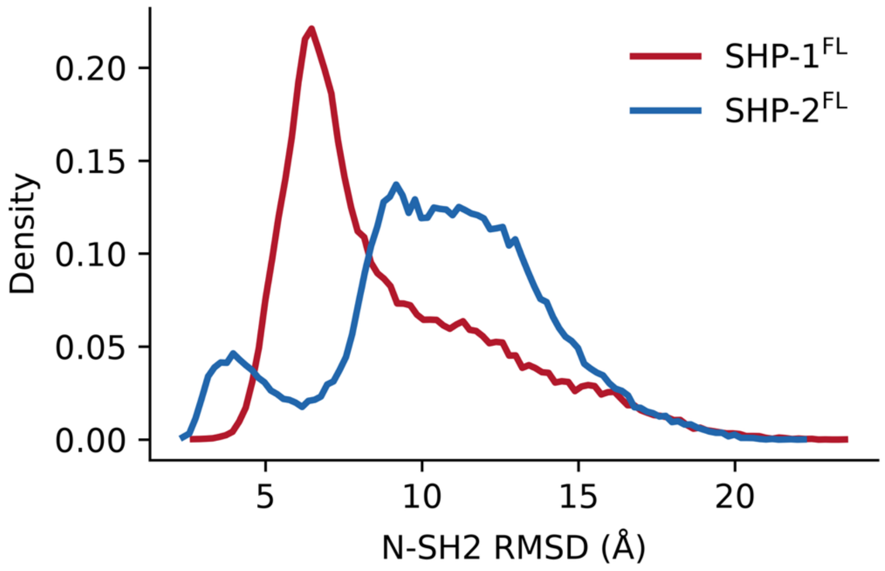


**Figure S5.** Probability distributions of the root-mean-square deviations (RMSD) of the full-length SHP-1 and SHP-2 NSH2 domains. RMSD was calculated on all $C_\alpha$-atoms in the N-SH2 domain, after aligning to the $C_\alpha$-atoms of the catalytic domain.

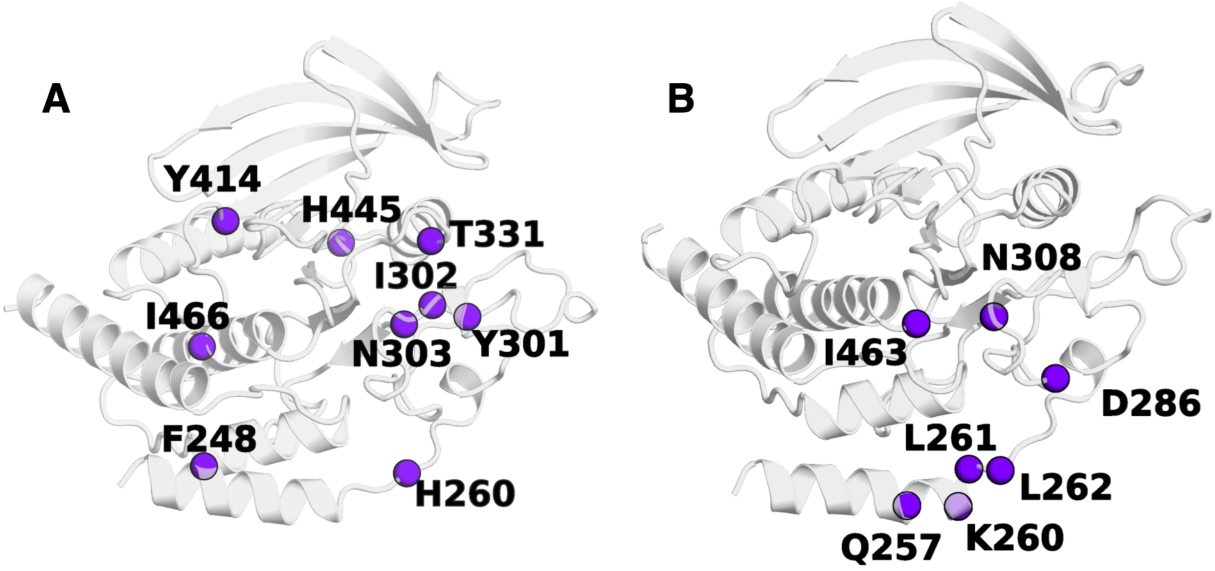


**Figure S6.** Showcasing the positions of allosterically non-conserved residue found on the shortest path map (SPM) pathways calculated for (**A**) SHP-1 and (**B**) SHP-2.

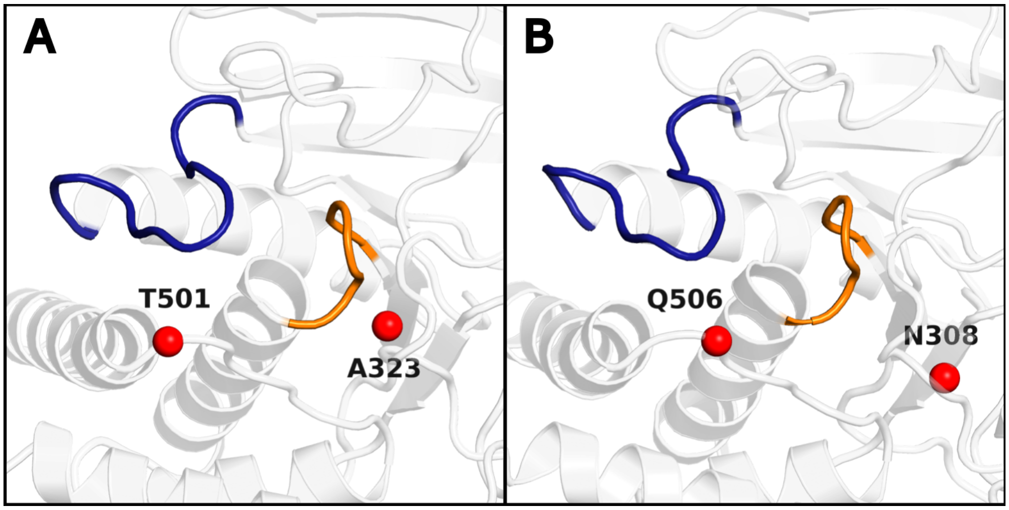


**Figure S7.** Location of the disease-causing amino acid substitutions studied in this work relative to the (**A**) SHP-1 and (**B**) SHP-2 active sites. The WPD-loop is shown in blue, and P-loop is shown in orange. The following SHP variants were considered: A323T, T501M (SHP-1), and Q506P, N308D (SHP-2).

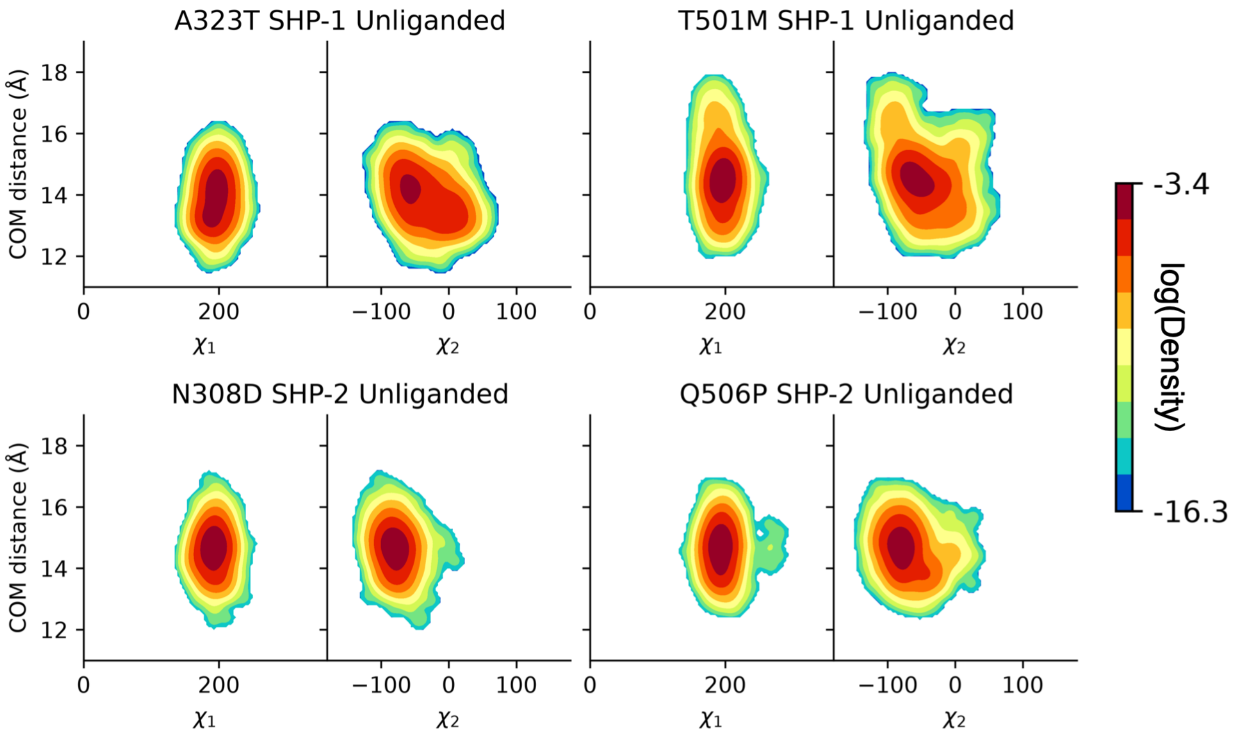


**Figure S8.** Joint probability distribution of W417 (SHP-1 numbering) $\chi_1$ and $\chi_2$ dihedrals versus the center of mass (COM) distance between the WPD-loop and P-loop. Results collected over 8 x 1.5µs molecular dynamics simulations SHP-1 A323T, SHP-1 T501M, SHP-2 N308D, and SHP-2 Q506P variants. The corresponding wild-type SHP phosphatase results are shown in **Figure S4**.

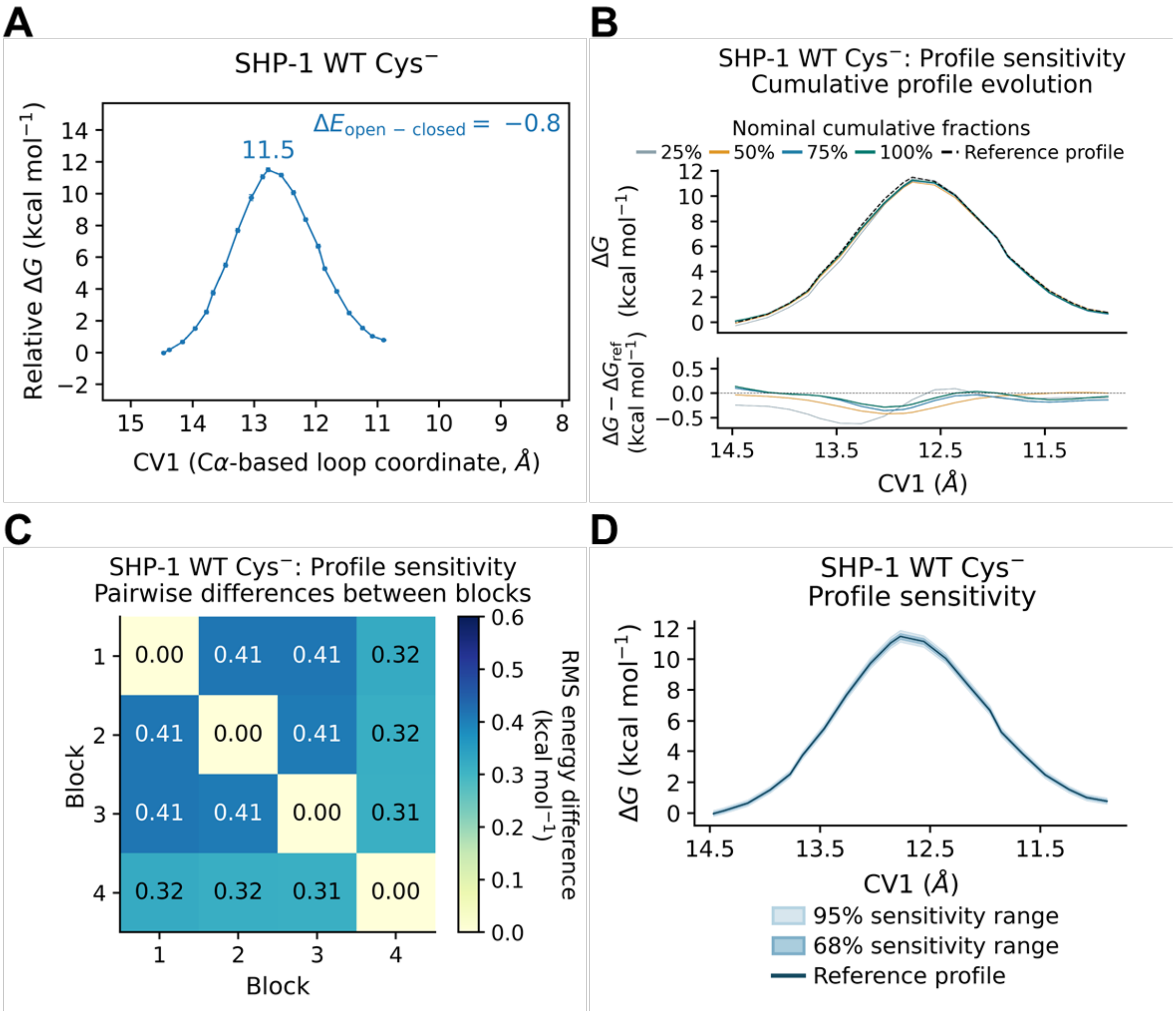


**Figure S9. MMVT free-energy profile, convergence analysis, and uncertainty for simulations of unliganded (Cys⁻) wild-type SHP-1.** The first 25% of each anchor's production trajectory was discarded as equilibration. All sampling fractions refer to the retained data. Profiles are displayed from open to closed WPD-loops, and referenced to the open state. **(A)** Free-energy profile along CV1, the $C_\alpha$-atom-based WPD-loop/P-loop coordinate, reconstructed from the retained MMVT sampling. Annotations indicate the barrier height relative to the lower-energy endpoint and the open–closed energy difference, $\Delta E_{open\text{–}closed} = E_{open} - E_{closed}$. Error bars represent pointwise 68% bootstrap confidence intervals. **(B)** Cumulative free-energy profiles reconstructed using the first 25%, 50%, 75%, and 100% of the retained sampling within each anchor. The dashed black curve represents the profile obtained from the full retained dataset. The lower panel shows deviations from this reference. **(C)** Pairwise root-mean-square (RMS) energy differences between profiles reconstructed from four consecutive, non-overlapping blocks, each containing 25% of the retained sampling within each anchor. Differences were calculated over the common CV1 grid after referencing each profile to its open endpoint. Blocks 1–4 denote disjoint sampling intervals, whereas the percentages in panel B denote cumulative sampling. **(D)** Free-energy profile reconstructed from the full retained sampling, with central 68% (dark shading) and 95% (light shading) pointwise bootstrap confidence intervals. Intervals were obtained by resampling contiguous trajectory blocks within each anchor, repeating the MMVT free-energy reconstruction for each resample, and referencing each reconstructed profile to the open state. All energies and RMS energy differences are reported in kcal mol$^{-1}$.

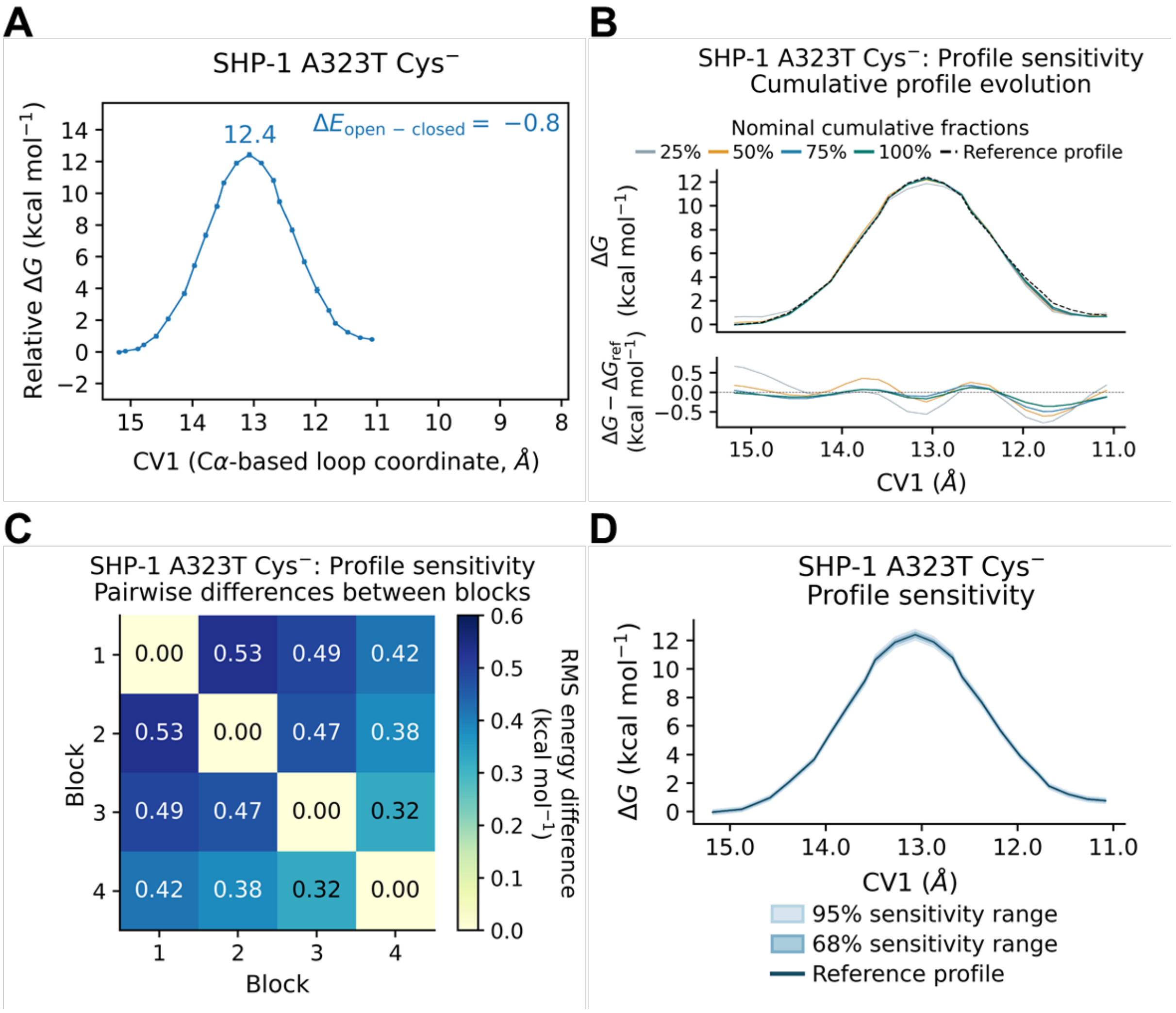


**Figure S10. MMVT free-energy profile, convergence analysis, and uncertainty for simulations of unliganded (Cys⁻) A323T SHP-1.** The first 25% of each anchor's production trajectory was discarded as equilibration. All sampling fractions refer to the retained data. Profiles are displayed from open to closed WPD-loops, and referenced to the open state. **(A)** Free-energy profile along CV1, the $C_\alpha$-atom-based WPD-loop/P-loop coordinate, reconstructed from the retained MMVT sampling. Annotations indicate the barrier height relative to the lower-energy endpoint and the open–closed energy difference, $\Delta E_{open–closed} = E_{open} - E_{closed}$. Error bars represent pointwise 68% bootstrap confidence intervals. **(B)** Cumulative free-energy profiles reconstructed using the first 25%, 50%, 75%, and 100% of the retained sampling within each anchor. The dashed

black curve represents the profile obtained from the full retained dataset. The lower panel shows deviations from this reference. **(C)** Pairwise root-mean-square (RMS) energy differences between profiles reconstructed from four consecutive, non-overlapping blocks, each containing 25% of the retained sampling within each anchor. Differences were calculated over the common CV1 grid after referencing each profile to its open endpoint. Blocks 1–4 denote disjoint sampling intervals, whereas the percentages in panel B denote cumulative sampling. **(D)** Free-energy profile reconstructed from the full retained sampling, with central 68% (dark shading) and 95% (light shading) pointwise bootstrap confidence intervals. Intervals were obtained by resampling contiguous trajectory blocks within each anchor, repeating the MMVT free-energy reconstruction for each resample, and referencing each reconstructed profile to the open state. All energies and RMS energy differences are reported in kcal $mol^{-1}$.

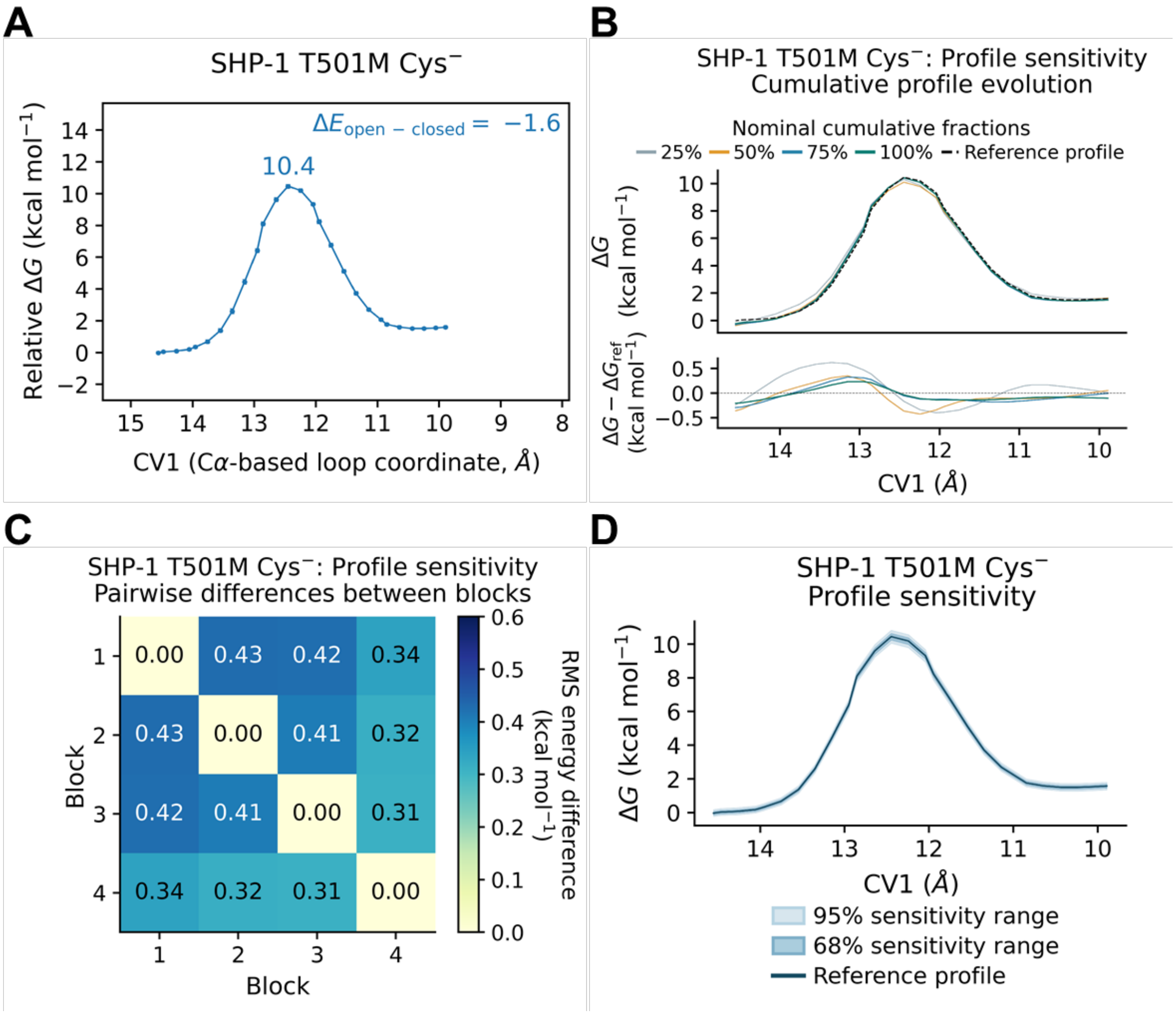


**Figure S11. MMVT free-energy profile, convergence analysis, and uncertainty for simulations of unliganded (Cys⁻) T501M SHP-1.** The first 25% of each anchor's production trajectory was discarded as equilibration. All sampling fractions refer to the retained data. Profiles are displayed from open to closed WPD-loops, and referenced to the open state. **(A)** Free-energy profile along CV1, the $C_\alpha$-atom-based WPD-loop/P-loop coordinate, reconstructed from the retained MMVT sampling. Annotations indicate the barrier height relative to the lower-energy endpoint and the open–closed energy difference, $\Delta E_{open\text{–}closed} = E_{open} - E_{closed}$. Error bars represent pointwise 68% bootstrap confidence intervals. **(B)** Cumulative free-energy profiles reconstructed using the first 25%, 50%, 75%, and 100% of the retained sampling within each anchor. The dashed black curve represents the profile obtained from the full retained dataset. The lower panel shows deviations from this reference. **(C)** Pairwise root-mean-square (RMS) energy differences between profiles reconstructed from four consecutive, non-overlapping blocks, each containing 25% of the retained sampling within each anchor. Differences were calculated over the common CV1 grid after referencing each profile to its open endpoint. Block 1–4 denote disjoint sampling intervals, whereas the percentages in panel B denote cumulative sampling. **(D)** Free-energy profile reconstructed from the full retained sampling, with central 68% (dark shading) and 95% (light shading) pointwise bootstrap confidence intervals. Intervals were obtained by resampling contiguous trajectory blocks within each anchor, repeating the MMVT free-energy reconstruction for each resample, and referencing each reconstructed profile to the open state. All energies and RMS energy differences are reported in kcal $mol^{-1}$.

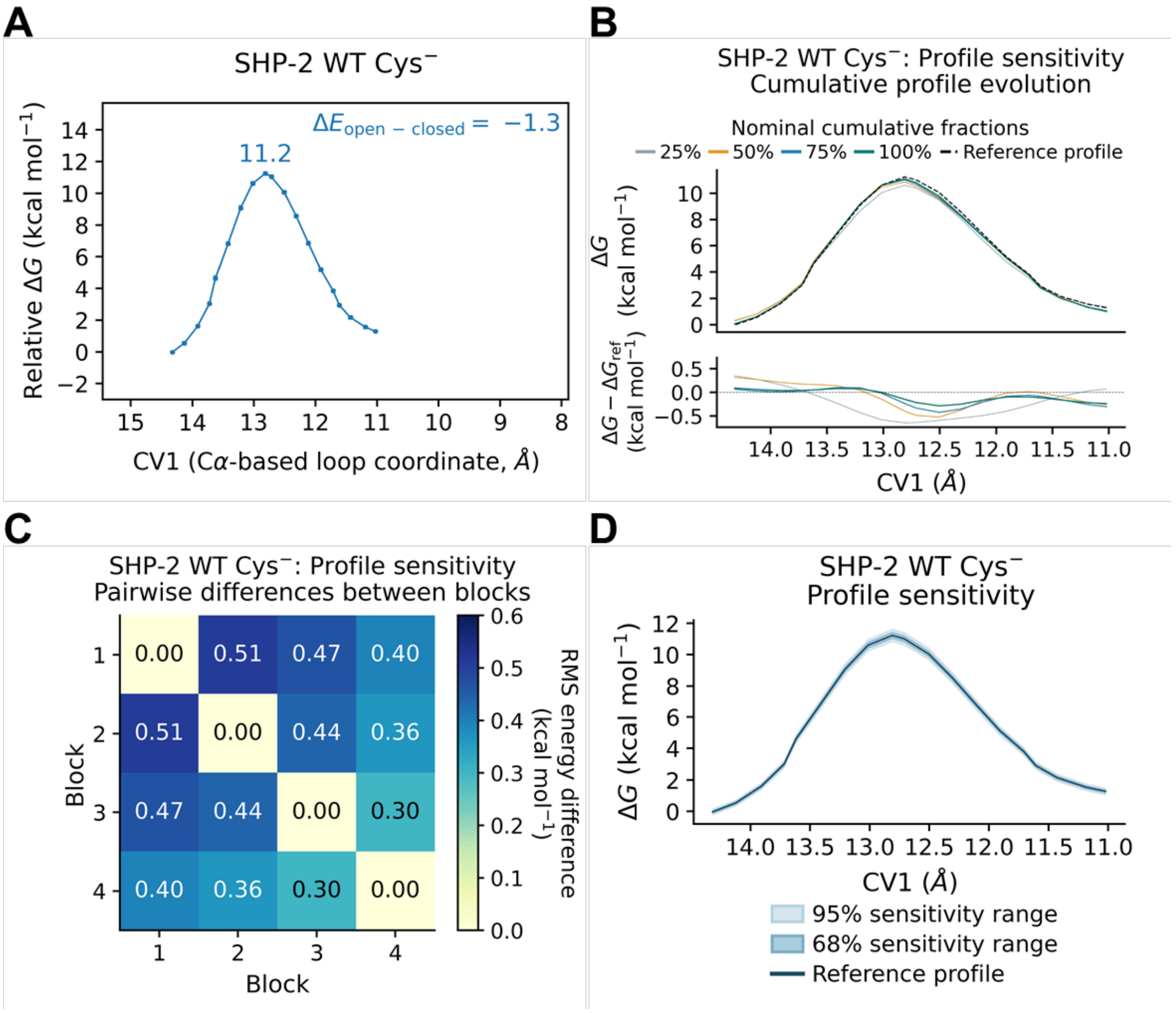


**Figure S12. MMVT free-energy profile, convergence analysis, and uncertainty for simulations of unliganded ($Cys^{-}$) wild-type SHP-2.** The first 25% of each anchor's production trajectory was discarded as equilibration. All sampling fractions refer to the retained data. Profiles are displayed from open to closed and referenced to the open state. **(A)** Free-energy profile along CV1, the $C_{\alpha}$-atom-based WPD-loop/P-loop coordinate, reconstructed from the retained MMVT sampling. Annotations indicate the barrier height relative to the lower-energy endpoint and the open–closed energy difference, $\Delta E_{open\text{–}closed} = E_{open} - E_{closed}$. Error bars represent pointwise 68% bootstrap confidence intervals. **(B)** Cumulative free-energy profiles reconstructed using the first 25%, 50%, 75%, and 100% of the retained sampling within each anchor. The dashed black curve represents the profile obtained from the full retained dataset. The lower panel shows deviations from this reference. **(C)** Pairwise root-mean-square (RMS) energy differences between profiles reconstructed from four consecutive, non-overlapping blocks, each containing 25% of the retained sampling within each anchor. Differences were calculated over the common CV1 grid after referencing each profile to its open endpoint. Blocks 1–4 denote disjoint sampling intervals, whereas the percentages in panel B denote cumulative sampling. **(D)** Free-energy profile reconstructed from the full retained sampling, with central 68% (dark shading) and 95% (light shading) pointwise bootstrap confidence intervals. Intervals were obtained by resampling contiguous trajectory blocks within each anchor, repeating the MMVT free-energy reconstruction for each resample, and referencing each reconstructed profile to the open state. All energies and RMS energy differences are reported in kcal $mol^{-1}$.

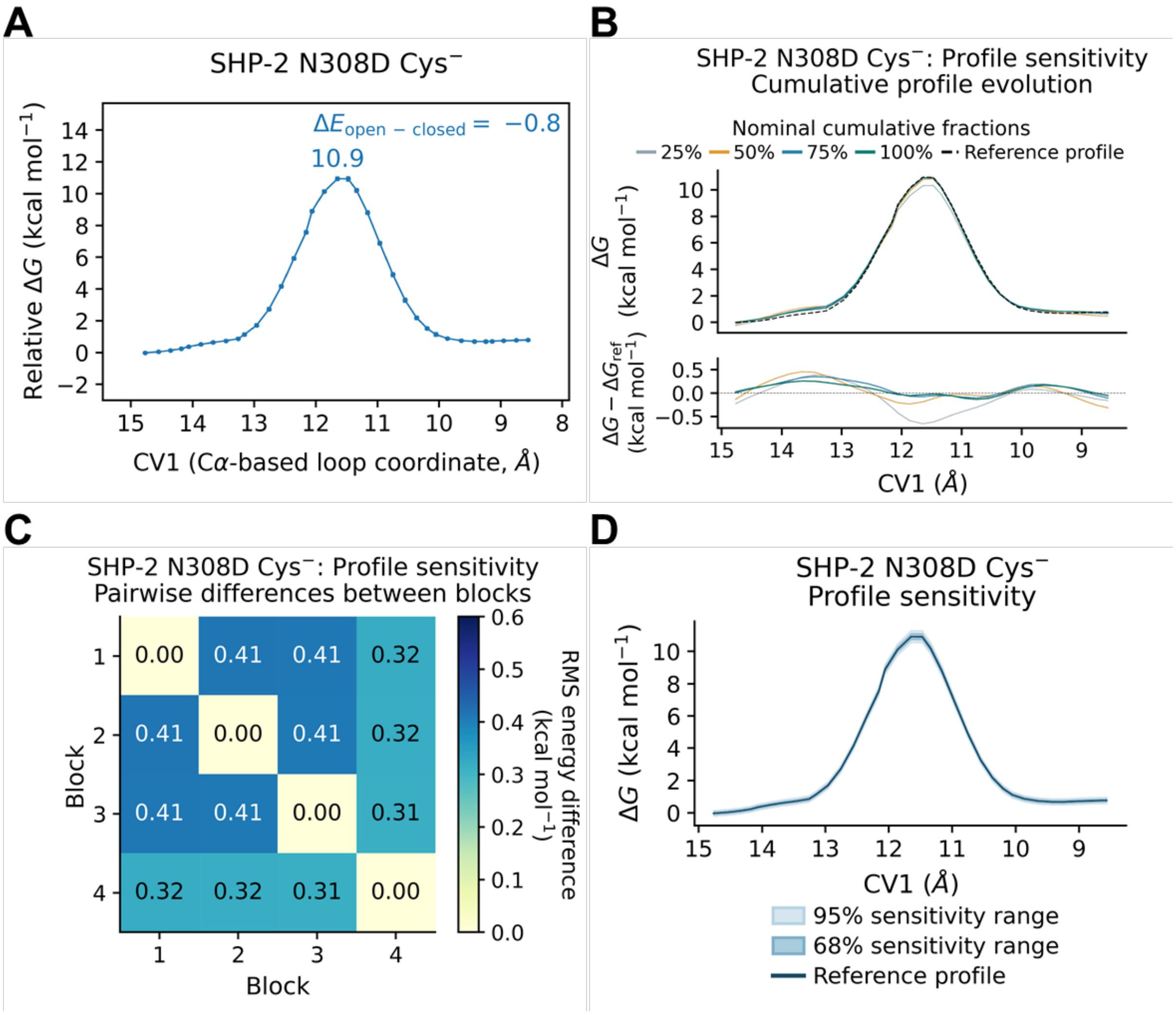


**Figure S13. MMVT free-energy profile, convergence analysis, and uncertainty for simulations of unliganded (Cys⁻) N308D SHP-2.** The first 25% of each anchor's production trajectory was discarded as equilibration. All sampling fractions refer to the retained data. Profiles are displayed from open to closed WPD-loop and referenced to the open state. **(A)** Free-energy profile along CV1, the $C_\alpha$-atom-based WPD-loop/P-loop coordinate, reconstructed from the retained MMVT sampling. Annotations indicate the barrier height relative to the lower-energy endpoint and the open–closed energy difference, $\Delta E_{open–closed} = E_{open} - E_{closed}$. Error bars represent pointwise 68% bootstrap confidence intervals. **(B)** Cumulative free-energy profiles reconstructed using the first 25%, 50%, 75%, and 100% of the retained sampling within each anchor. The dashed

black curve represents the profile obtained from the full retained dataset. The lower panel shows deviations from this reference. **(C)** Pairwise root-mean-square (RMS) energy differences between profiles reconstructed from four consecutive, non-overlapping blocks, each containing 25% of the retained sampling within each anchor. Differences were calculated over the common CV1 grid after referencing each profile to its open endpoint. Blocks 1–4 denote disjoint sampling intervals, whereas the percentages in panel B denote cumulative sampling. **(D)** Free-energy profile reconstructed from the full retained sampling, with central 68% (dark shading) and 95% (light shading) pointwise bootstrap confidence intervals. Intervals were obtained by resampling contiguous trajectory blocks within each anchor, repeating the MMVT free-energy reconstruction for each resample, and referencing each reconstructed profile to the open state. All energies and RMS energy differences are reported in kcal $mol^{-1}$.

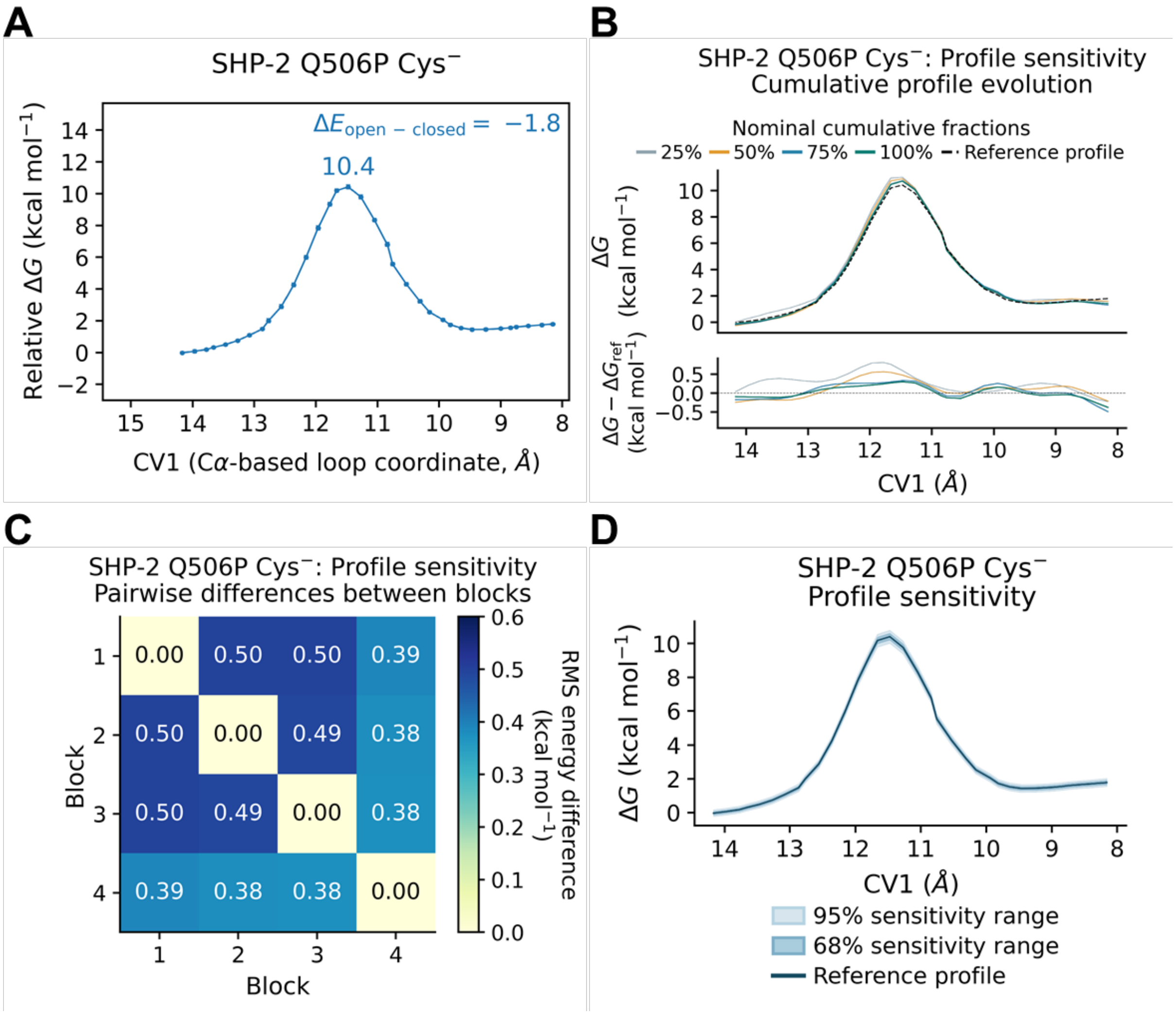


**Figure S14. MMVT free-energy profile, convergence analysis, and uncertainty for simulations of unliganded (Cys⁻) Q506P SHP-2.** The first 25% of each anchor's production trajectory was discarded as equilibration. All sampling fractions refer to the retained data. Profiles are displayed from open to closed WPD-loop and referenced to the open state. **(A)** Free-energy profile along CV1, the $C_\alpha$-atom-based WPD-loop/P-loop coordinate, reconstructed from the retained MMVT sampling. Annotations indicate the barrier height relative to the lower-energy endpoint and the open–closed energy difference, $\Delta E_{open-closed} = E_{open} - E_{closed}$. Error bars represent pointwise 68% bootstrap confidence intervals. **(B)** Cumulative free-energy profiles reconstructed using the first 25%, 50%, 75%, and 100% of the retained sampling within each anchor. The dashed

black curve represents the profile obtained from the full retained dataset. The lower panel shows deviations from this reference. **(C)** Pairwise root-mean-square (RMS) energy differences between profiles reconstructed from four consecutive, non-overlapping blocks, each containing 25% of the retained sampling within each anchor. Differences were calculated over the common CV1 grid after referencing each profile to its open endpoint. Blocks 1–4 denote disjoint sampling intervals, whereas the percentages in panel B denote cumulative sampling. **(D)** Free-energy profile reconstructed from the full retained sampling, with central 68% (dark shading) and 95% (light shading) pointwise bootstrap confidence intervals. Intervals were obtained by resampling contiguous trajectory blocks within each anchor, repeating the MMVT free-energy reconstruction for each resample, and referencing each reconstructed profile to the open state. All energies and RMS energy differences are reported in kcal $mol^{-1}$.

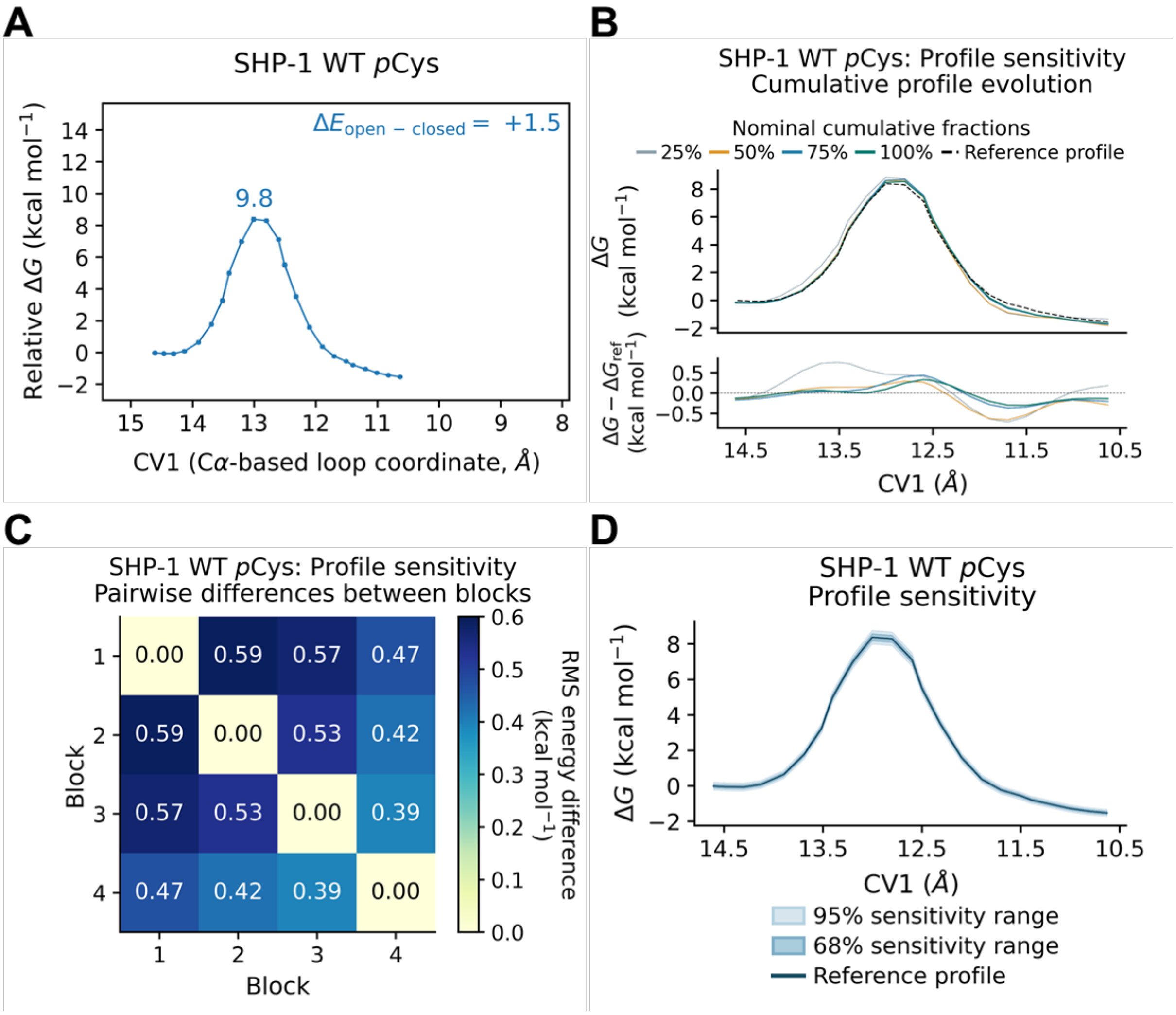


**Figure S15. MMVT free-energy profile, convergence analysis, and uncertainty for wild-type SHP-1 at the phosphoenzyme intermediate state (*p*Cys).** The first 25% of each anchor's production trajectory was discarded as equilibration. All sampling fractions refer to the retained data. Profiles are displayed from open to closed WPD-loop and referenced to the open state. **(A)** Free-energy profile along CV1, the $C_\alpha$-atom-based WPD-loop/P-loop coordinate, reconstructed from the retained MMVT sampling. Annotations indicate the barrier height relative to the lower-energy endpoint and the open–closed energy difference, $\Delta E_{open\text{–}closed} = E_{open} - E_{closed}$. Error bars represent pointwise 68% bootstrap confidence intervals. **(B)** Cumulative free-energy profiles reconstructed using the first 25%, 50%, 75%, and 100% of the retained sampling within each

anchor. The dashed black curve represents the profile obtained from the full retained dataset. The lower panel shows deviations from this reference. **(C)** Pairwise root-mean-square (RMS) energy differences between profiles reconstructed from four consecutive, non-overlapping blocks, each containing 25% of the retained sampling within each anchor. Differences were calculated over the common CV1 grid after referencing each profile to its open endpoint. Blocks 1–4 denote disjoint sampling intervals, whereas the percentages in panel B denote cumulative sampling. **(D)** Free-energy profile reconstructed from the full retained sampling, with central 68% (dark shading) and 95% (light shading) pointwise bootstrap confidence intervals. Intervals were obtained by resampling contiguous trajectory blocks within each anchor, repeating the MMVT free-energy reconstruction for each resample, and referencing each reconstructed profile to the open state. All energies and RMS energy differences are reported in kcal $mol^{-1}$.

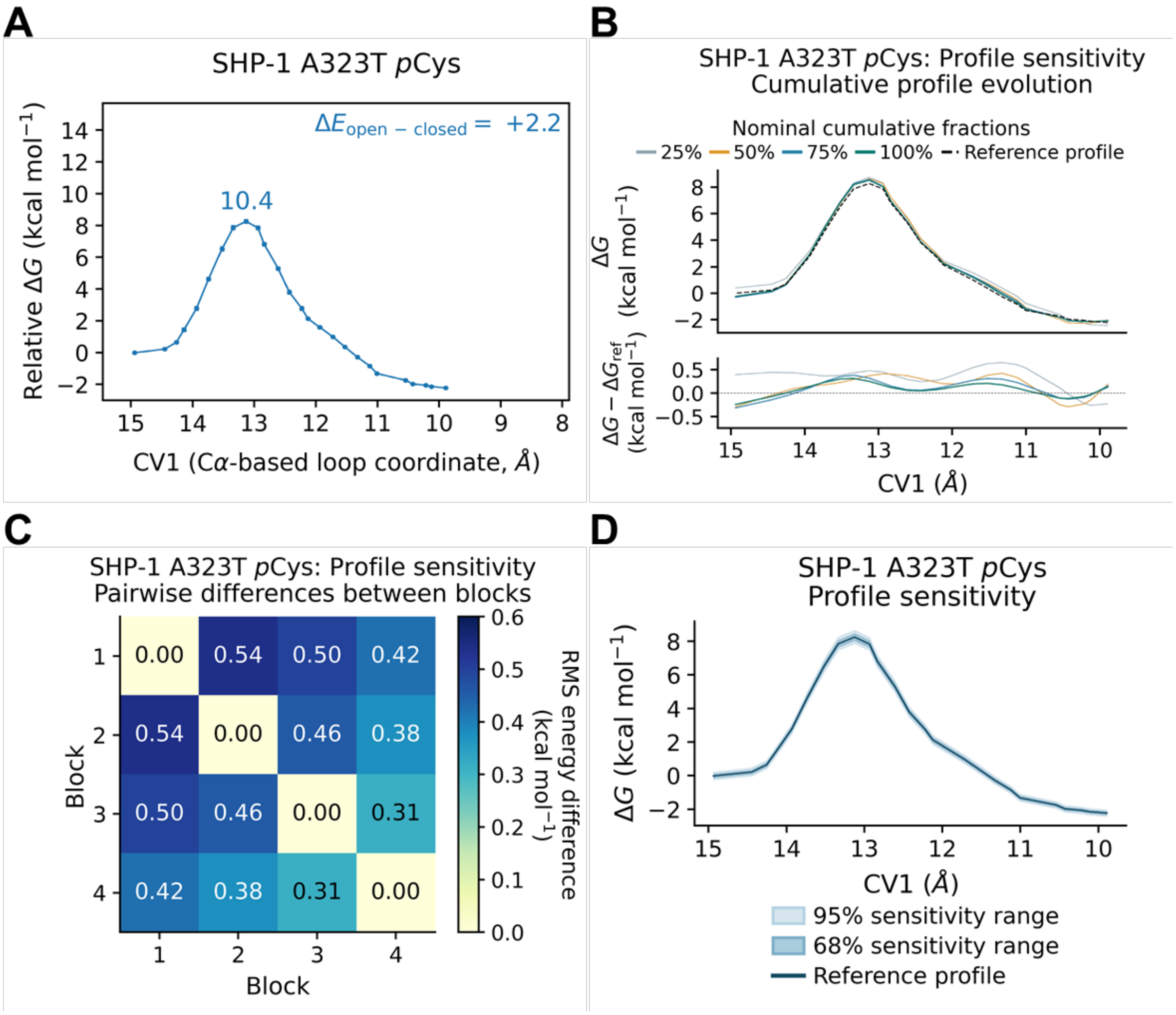


**Figure S16. MMVT free-energy profile, convergence analysis, and uncertainty for A323T SHP-1 at the phosphoenzyme intermediate state (*p*Cys).** The first 25% of each anchor's production trajectory was discarded as equilibration. All sampling fractions refer to the retained data. Profiles are displayed from open to closed WPD-loop and referenced to the open state. **(A)** Free-energy profile along CV1, the $C_\alpha$-atom-based WPD-loop/P-loop coordinate, reconstructed from the retained MMVT sampling. Annotations indicate the barrier height relative to the lower-energy endpoint and the open–closed energy difference, $\Delta E_{open\text{–}closed} = E_{open} - E_{closed}$. Error bars represent pointwise 68% bootstrap confidence intervals. **(B)** Cumulative free-energy profiles reconstructed using the first 25%, 50%, 75%, and 100% of the retained sampling within each

anchor. The dashed black curve represents the profile obtained from the full retained dataset. The lower panel shows deviations from this reference. **(C)** Pairwise root-mean-square (RMS) energy differences between profiles reconstructed from four consecutive, non-overlapping blocks, each containing 25% of the retained sampling within each anchor. Differences were calculated over the common CV1 grid after referencing each profile to its open endpoint. Blocks 1–4 denote disjoint sampling intervals, whereas the percentages in panel B denote cumulative sampling. **(D)** Free-energy profile reconstructed from the full retained sampling, with central 68% (dark shading) and 95% (light shading) pointwise bootstrap confidence intervals. Intervals were obtained by resampling contiguous trajectory blocks within each anchor, repeating the MMVT free-energy reconstruction for each resample, and referencing each reconstructed profile to the open state. All energies and RMS energy differences are reported in kcal $mol^{-1}$.

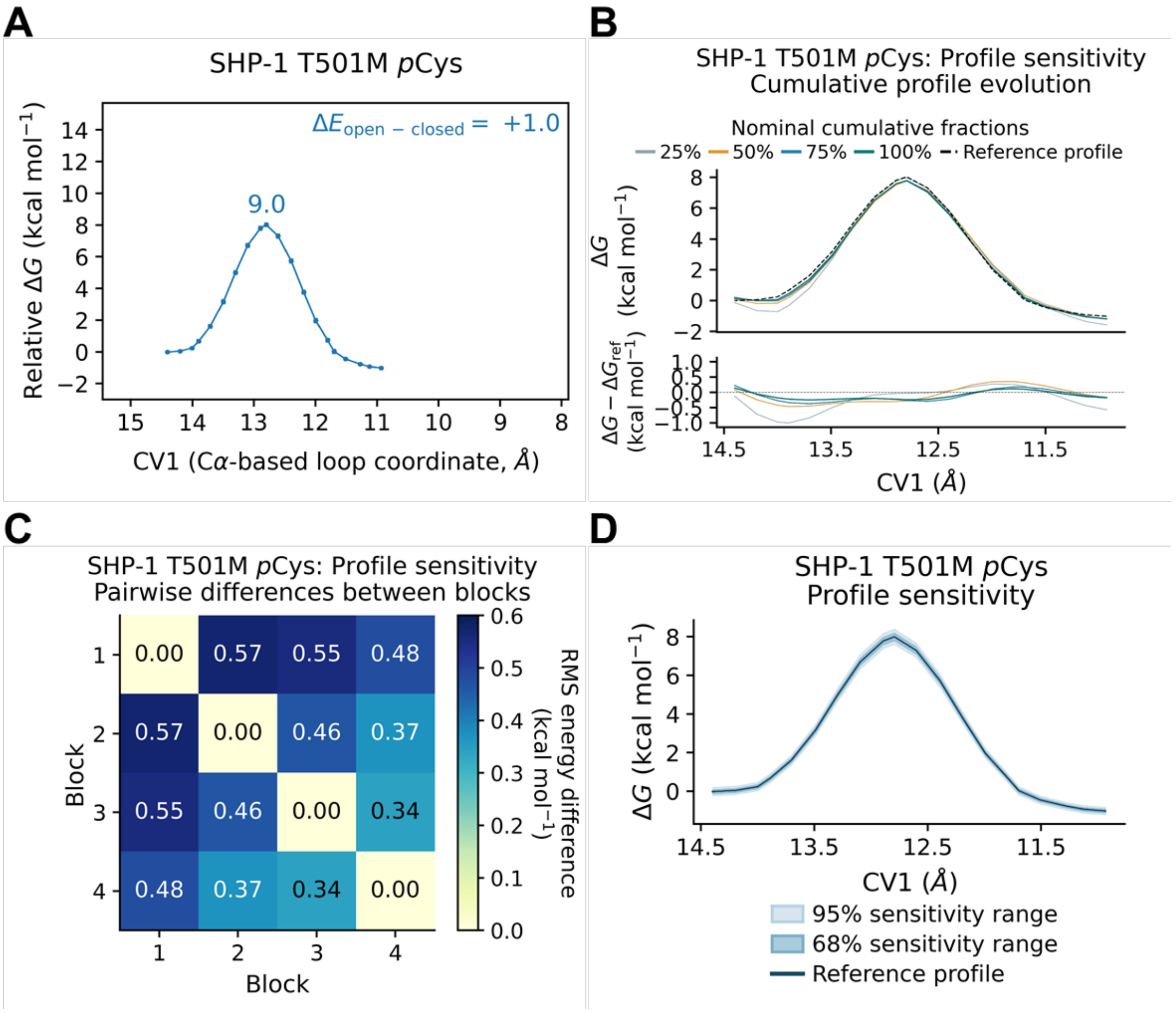


**Figure S17. MMVT free-energy profile, convergence analysis, and uncertainty for T501M SHP-1 at the phosphoenzyme intermediate state (*p*Cys).** The first 25% of each anchor's production trajectory was discarded as equilibration. All sampling fractions refer to the retained data. Profiles are displayed from open to closed WPD-loop and referenced to the open state. **(A)** Free-energy profile along CV1, the $C_\alpha$-atom-based WPD-loop/P-loop coordinate, reconstructed from the retained MMVT sampling. Annotations indicate the barrier height relative to the lower-energy endpoint and the open–closed energy difference, $\Delta E_{open\text{–}closed} = E_{open} - E_{closed}$. Error bars represent pointwise 68% bootstrap confidence intervals. **(B)** Cumulative free-energy profiles reconstructed using the first 25%, 50%, 75%, and 100% of the retained sampling within each

anchor. The dashed black curve represents the profile obtained from the full retained dataset. The lower panel shows deviations from this reference. **(C)** Pairwise root-mean-square (RMS) energy differences between profiles reconstructed from four consecutive, non-overlapping blocks, each containing 25% of the retained sampling within each anchor. Differences were calculated over the common CV1 grid after referencing each profile to its open endpoint. Blocks 1–4 denote disjoint sampling intervals, whereas the percentages in panel B denote cumulative sampling. **(D)** Free-energy profile reconstructed from the full retained sampling, with central 68% (dark shading) and 95% (light shading) pointwise bootstrap confidence intervals. Intervals were obtained by resampling contiguous trajectory blocks within each anchor, repeating the MMVT free-energy reconstruction for each resample, and referencing each reconstructed profile to the open state. All energies and RMS energy differences are reported in kcal $mol^{-1}$.

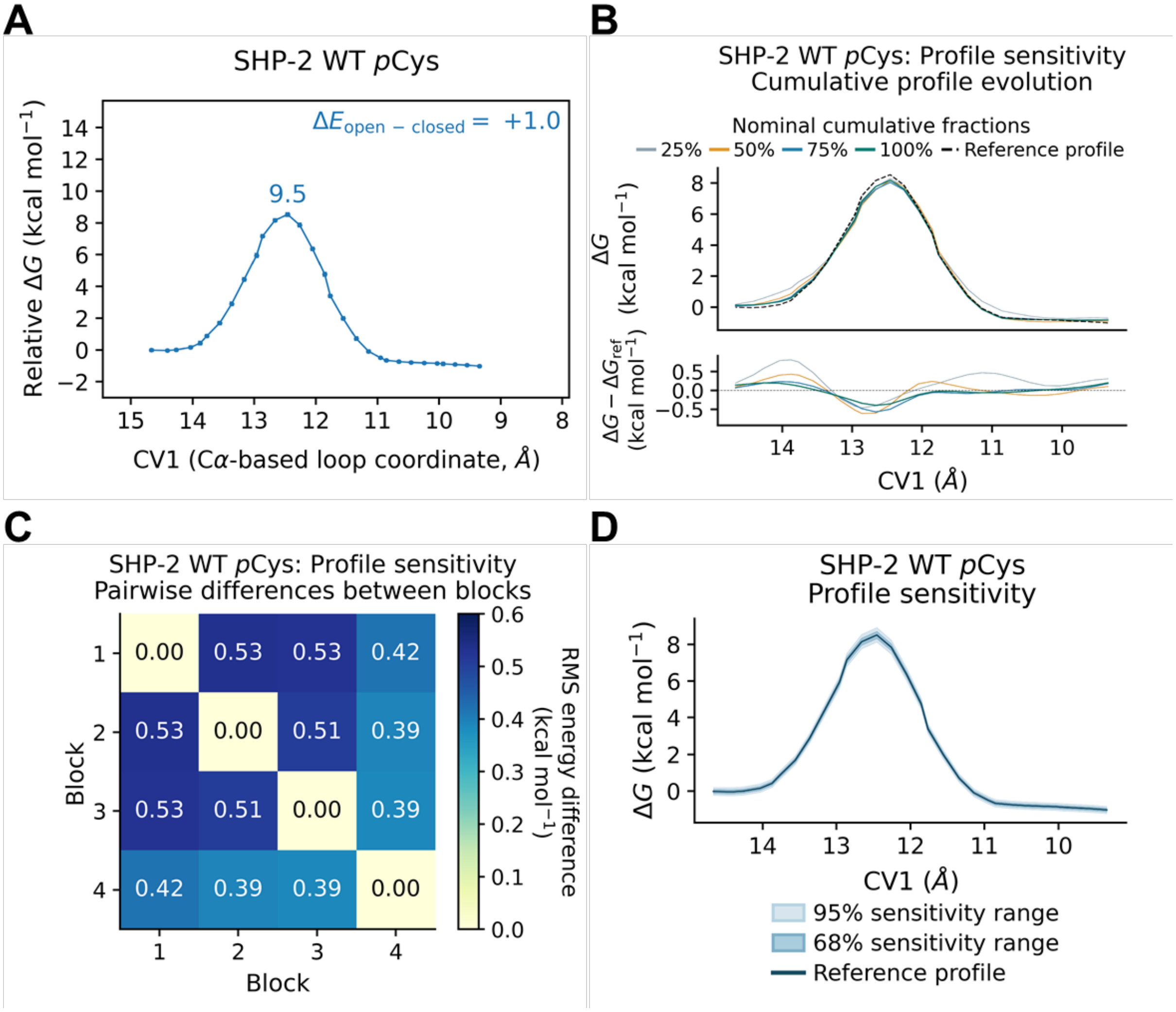


**Figure S18. MMVT free-energy profile, convergence analysis, and uncertainty for wild-type SHP-2 at the phosphoenzyme intermediate state (*p*Cys).** The first 25% of each anchor's production trajectory was discarded as equilibration. All sampling fractions refer to the retained data. Profiles are displayed from open to closed WPD-loop and referenced to the open state. **(A)** Free-energy profile along CV1, the $C_\alpha$-atom-based WPD-loop/P-loop coordinate, reconstructed from the retained MMVT sampling. Annotations indicate the barrier height relative to the lower-energy endpoint and the open–closed energy difference, $\Delta E_{open\text{–}closed} = E_{open} - E_{closed}$. Error bars represent pointwise 68% bootstrap confidence intervals. **(B)** Cumulative free-energy profiles reconstructed using the first 25%, 50%, 75%, and 100% of the retained sampling within each

anchor. The dashed black curve represents the profile obtained from the full retained dataset. The lower panel shows deviations from this reference. **(C)** Pairwise root-mean-square (RMS) energy differences between profiles reconstructed from four consecutive, non-overlapping blocks, each containing 25% of the retained sampling within each anchor. Differences were calculated over the common CV1 grid after referencing each profile to its open endpoint. Blocks 1–4 denote disjoint sampling intervals, whereas the percentages in panel B denote cumulative sampling. **(D)** Free-energy profile reconstructed from the full retained sampling, with central 68% (dark shading) and 95% (light shading) pointwise bootstrap confidence intervals. Intervals were obtained by resampling contiguous trajectory blocks within each anchor, repeating the MMVT free-energy reconstruction for each resample, and referencing each reconstructed profile to the open state. All energies and RMS energy differences are reported in kcal $mol^{-1}$.

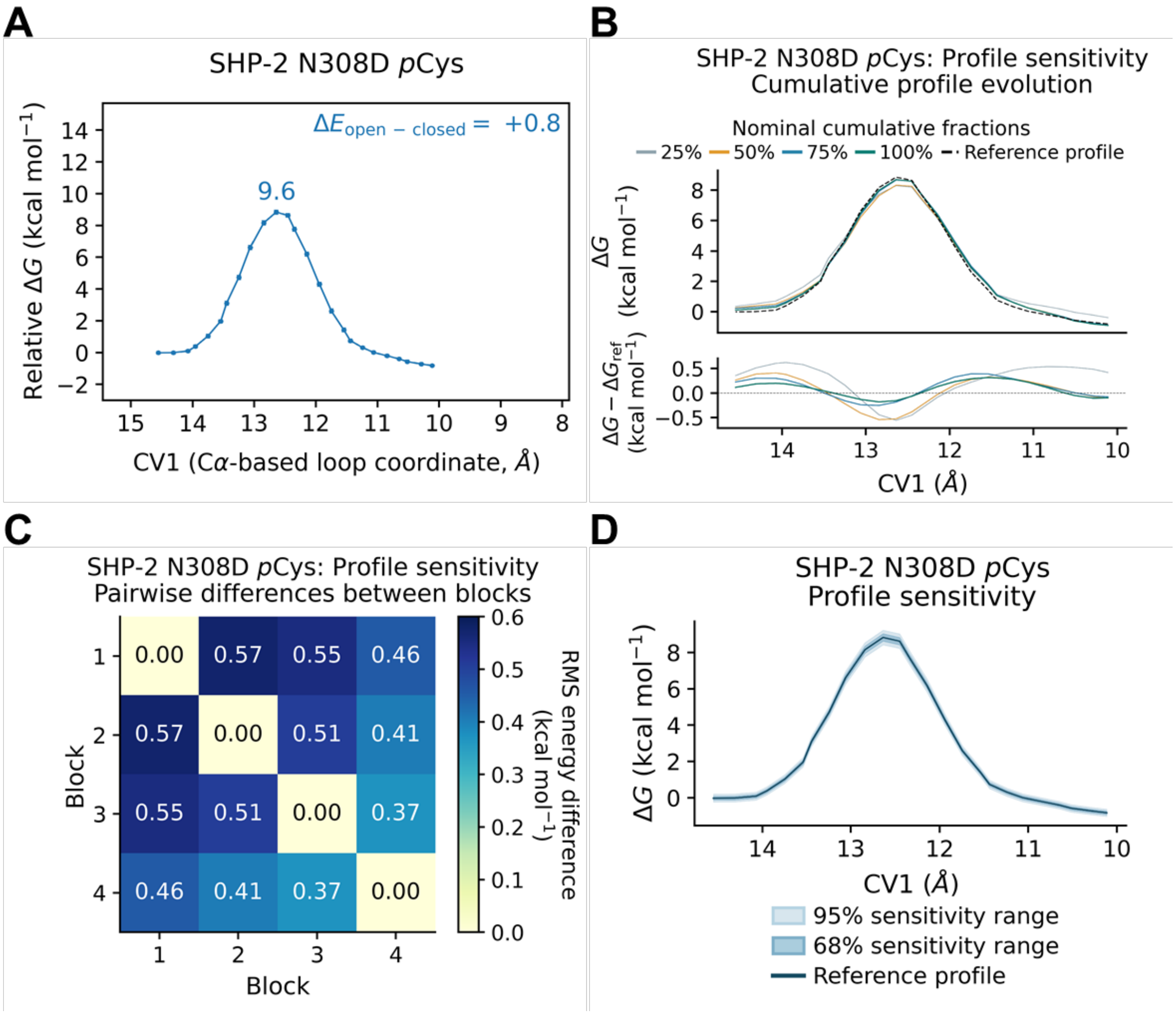


**Figure S19. MMVT free-energy profile, convergence analysis, and uncertainty for N308D SHP-2 at the phosphoenzyme intermediate state (*p*Cys).** The first 25% of each anchor's production trajectory was discarded as equilibration. All sampling fractions refer to the retained data. Profiles are displayed from open to closed WPD-loop and referenced to the open state. **(A)** Free-energy profile along CV1, the $C_\alpha$-atom-based WPD-loop/P-loop coordinate, reconstructed from the retained MMVT sampling. Annotations indicate the barrier height relative to the lower-energy endpoint and the open–closed energy difference, $\Delta E_{open\text{–}closed} = E_{open} - E_{closed}$. Error bars represent pointwise 68% bootstrap confidence intervals. **(B)** Cumulative free-energy profiles reconstructed using the first 25%, 50%, 75%, and 100% of the retained sampling within each

anchor. The dashed black curve represents the profile obtained from the full retained dataset. The lower panel shows deviations from this reference. **(C)** Pairwise root-mean-square (RMS) energy differences between profiles reconstructed from four consecutive, non-overlapping blocks, each containing 25% of the retained sampling within each anchor. Differences were calculated over the common CV1 grid after referencing each profile to its open endpoint. Blocks 1–4 denote disjoint sampling intervals, whereas the percentages in panel B denote cumulative sampling. **(D)** Free-energy profile reconstructed from the full retained sampling, with central 68% (dark shading) and 95% (light shading) pointwise bootstrap confidence intervals. Intervals were obtained by resampling contiguous trajectory blocks within each anchor, repeating the MMVT free-energy reconstruction for each resample, and referencing each reconstructed profile to the open state. All energies and RMS energy differences are reported in kcal $mol^{-1}$.

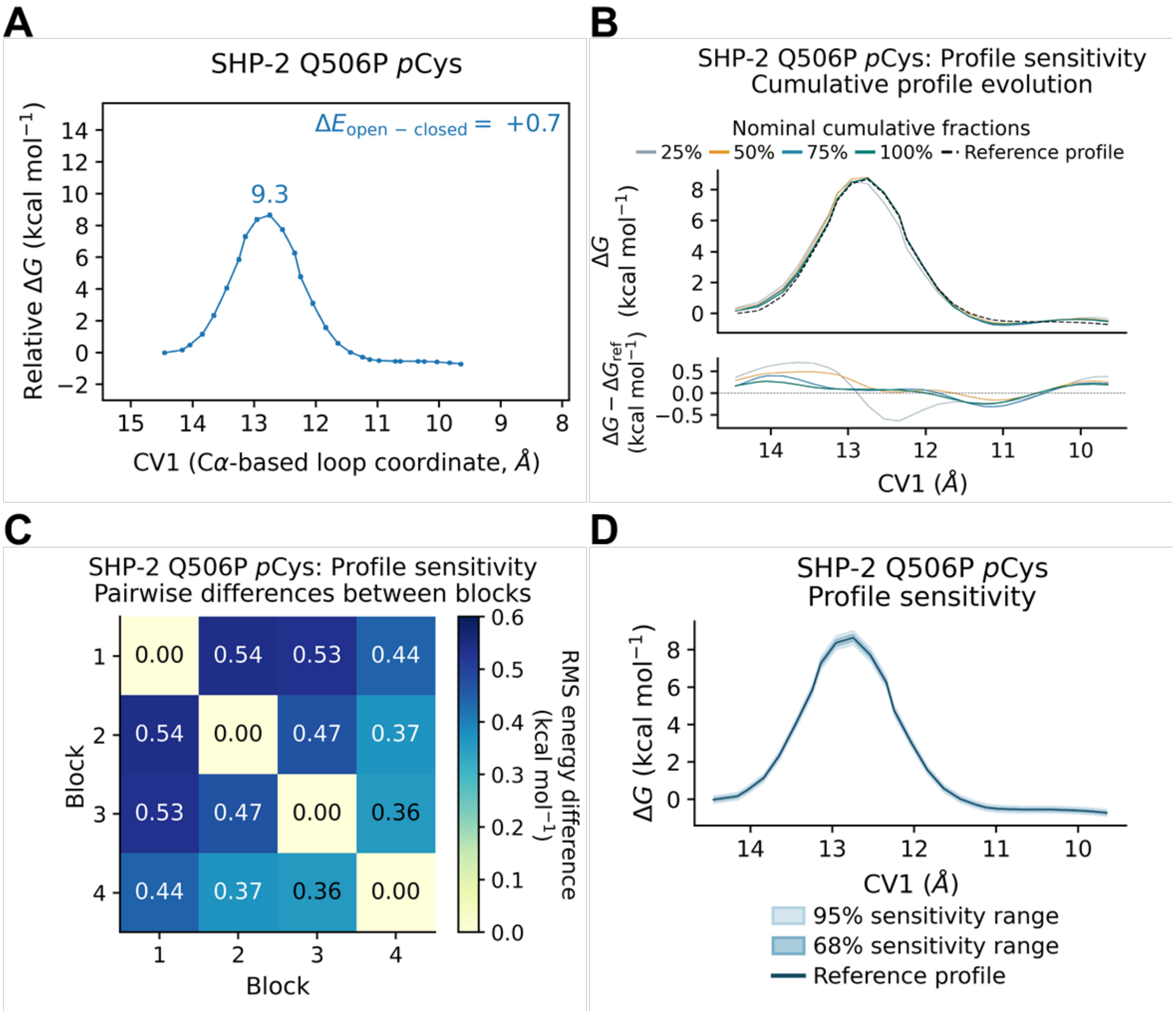


**Figure S20. MMVT free-energy profile, convergence analysis, and uncertainty for Q506P SHP-2 at the phosphoenzyme intermediate state (*p*Cys).** The first 25% of each anchor's production trajectory was discarded as equilibration. All sampling fractions refer to the retained data. Profiles are displayed from open to closed WPD-loop and referenced to the open state. **(A)** Free-energy profile along CV1, the $C_\alpha$-atom-based WPD-loop/P-loop coordinate, reconstructed from the retained MMVT sampling. Annotations indicate the barrier height relative to the lower-energy endpoint and the open–closed energy difference, $\Delta E_{open\text{–}closed} = E_{open} - E_{closed}$. Error bars represent pointwise 68% bootstrap confidence intervals. **(B)** Cumulative free-energy profiles reconstructed using the first 25%, 50%, 75%, and 100% of the retained sampling within each anchor. The dashed black curve represents the profile obtained from the full retained dataset. The lower panel shows deviations from this reference. **(C)** Pairwise root-mean-square (RMS) energy differences between profiles reconstructed from four consecutive, non-overlapping blocks, each containing 25% of the retained sampling within each anchor. Differences were calculated over the common CV1 grid after referencing each profile to its open endpoint. Blocks 1–4 denote disjoint sampling intervals, whereas the percentages in panel B denote cumulative sampling. **(D)** Free-energy profile reconstructed from the full retained sampling, with central 68% (dark shading) and 95% (light shading) pointwise bootstrap confidence intervals. Intervals were obtained by resampling contiguous trajectory blocks within each anchor, repeating the MMVT free-energy reconstruction for each resample, and referencing each reconstructed profile to the open state. All energies and RMS energy differences are reported in kcal mol$^{-1}$."

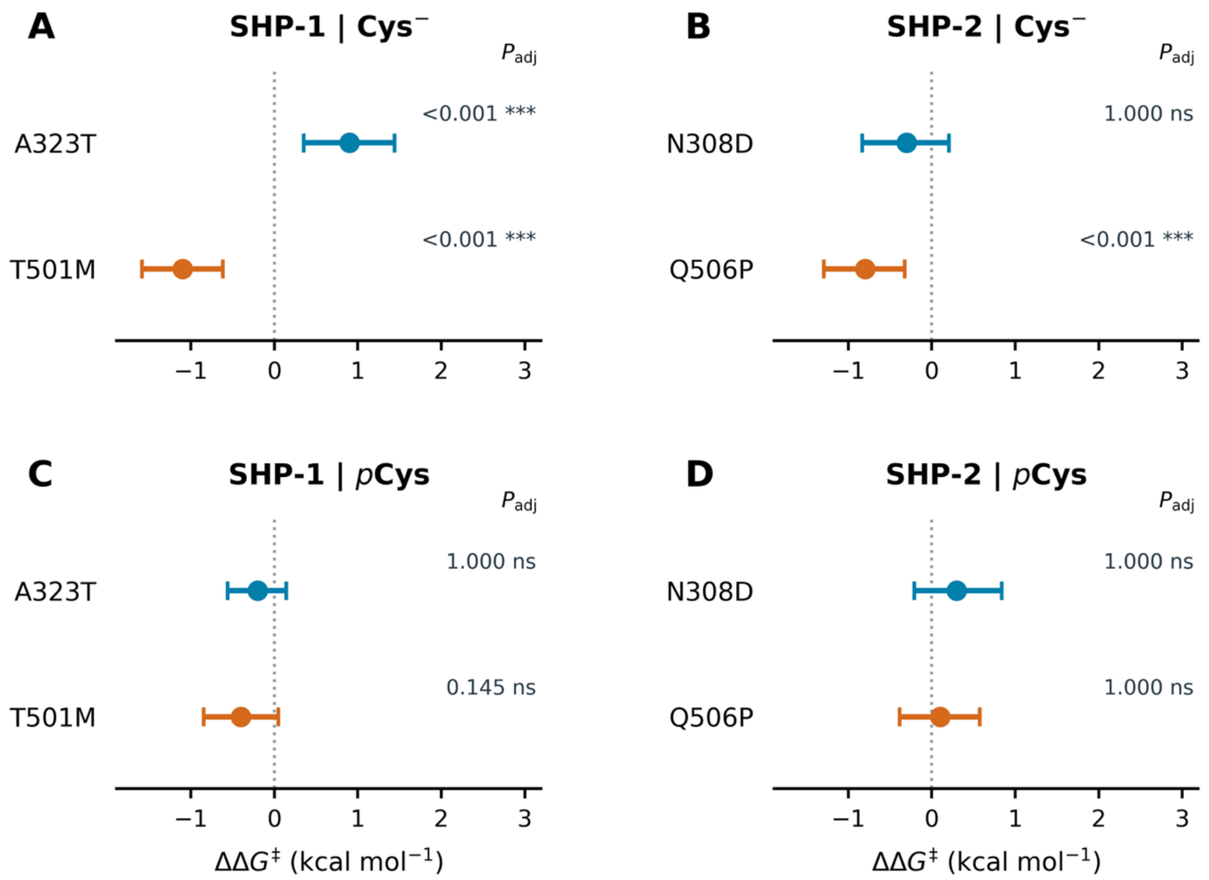


**Figure S21. Comparisons of WPD-loop closing barriers in SHP-1 and SHP-2.** Mutant – wild-type differences are shown for **(A)** unliganded SHP-1, **(B)** unliganded SHP-2, **(C)** phosphoenzyme-intermediate (*p*Cys) SHP-1, and **(D)** *p*Cys SHP-2. Closing barriers are defined relative to the open state: $\Delta G^{\ddagger} = G_{peak} - G_{open}$. Points represent differences between calculated MMVT means: $\Delta\Delta G^{\ddagger} = \Delta G^{\ddagger}(\text{mutant}) - \Delta G^{\ddagger}(\text{wild-type})$, expressed in kcal $mol^{-1}$. Blue denotes A323T/N308D, and orange denotes T501M/Q506P. Horizontal bars show simultaneous 95% basic block-bootstrap confidence intervals. The dotted vertical line indicates no difference from the wild type. Two-sided bootstrap *P* values are Bonferroni-adjusted across comparisons.[4, 5] One, two, and three asterisks indicate adjusted $P < 0.05$, $< 0.01$, and $< 0.001$, respectively; ns, not statistically significant.

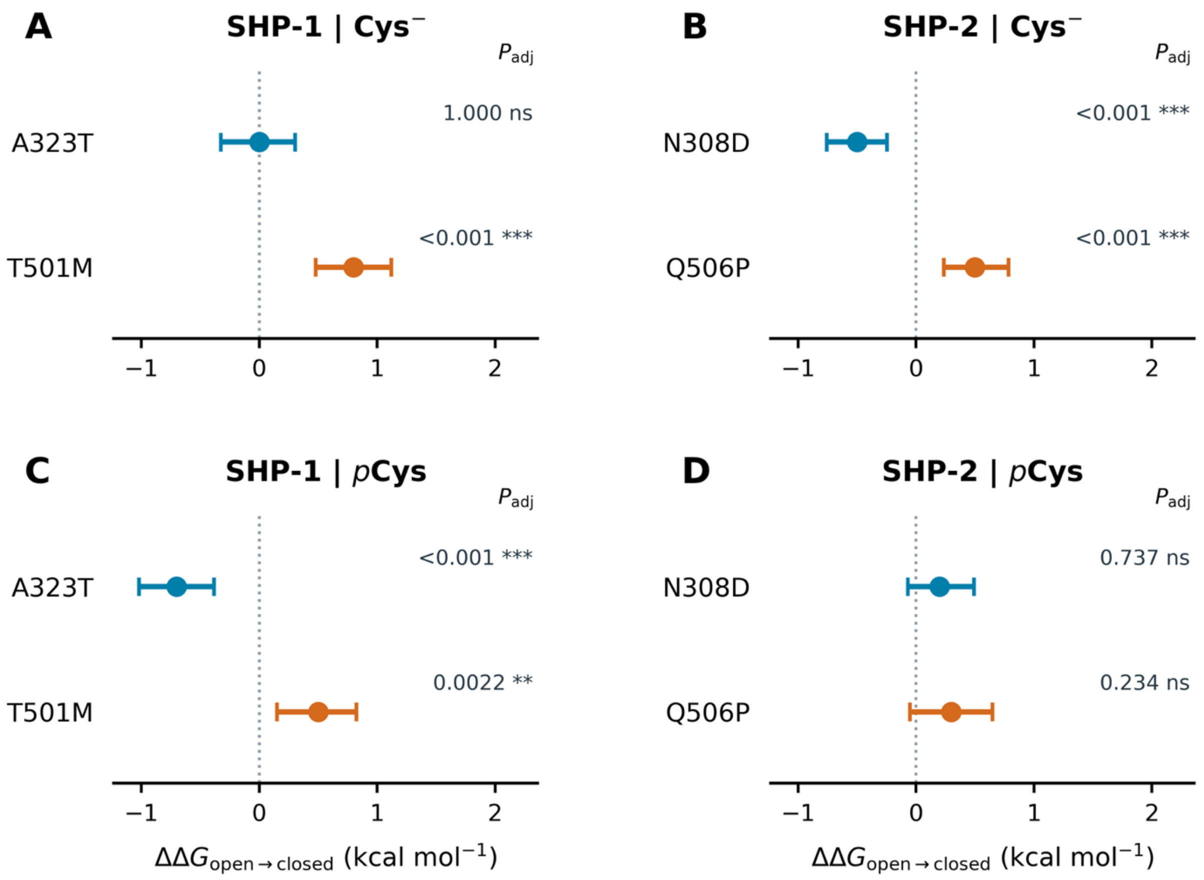


**Figure S22. Comparisons of reaction free energies for WPD-loop closure in SHP-1 and SHP-2.** Mutant – wild-type differences are shown for **(A)** unliganded SHP-1, **(B)** unliganded SHP-2, **(C)** phosphoenzyme-intermediate (*p*Cys) SHP-1, and **(D)** *p*Cys SHP-2. Reaction free energies are defined as $\Delta G_{open \rightarrow closed} = G_{closed} - G_{open}$. Points represent differences between calculated MMVT means: $\Delta\Delta G = \Delta G(\text{mutant}) - \Delta G(\text{wild-type})$, expressed in kcal $mol^{-1}$. Negative values indicate more favorable closure relative to wild-type, whereas positive values indicate less favorable closure. Blue denotes A323T/N308D, and orange denotes T501M/Q506P. Horizontal bars show simultaneous 95% basic block-bootstrap confidence intervals. The dotted vertical line indicates no difference from wild-type. Two-sided bootstrap *P* values are Bonferroni-adjusted across comparisons.[4, 5] One, two, and three asterisks indicate adjusted $P < 0.05$, $< 0.01$, and $< 0.001$, respectively; ns, not statistically significant.

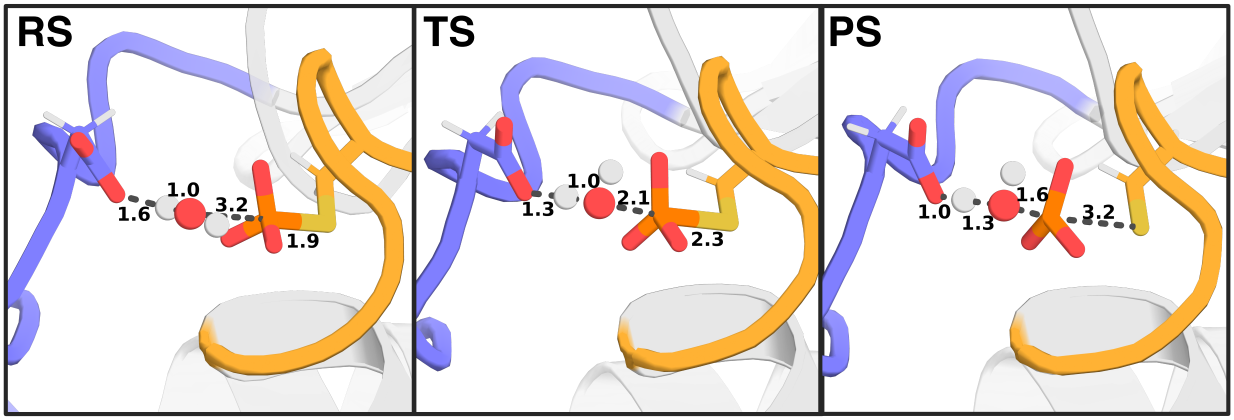


**Figure S23.** Shown here are also representative reactant (RS), transition (TS) and product (PS) state structures from our EVB trajectories, extracted from the dominant cluster obtained from hierarchical agglomerative clustering on reacting heavy atoms during empirical valence bond (EVB)[6] simulations of wild-type SHP-2, using CPPTRAJ.[7] The equivalent structures for SHP-1 are shown in **Figure 9**, and the corresponding reactive distances are shown in **Table S5**.

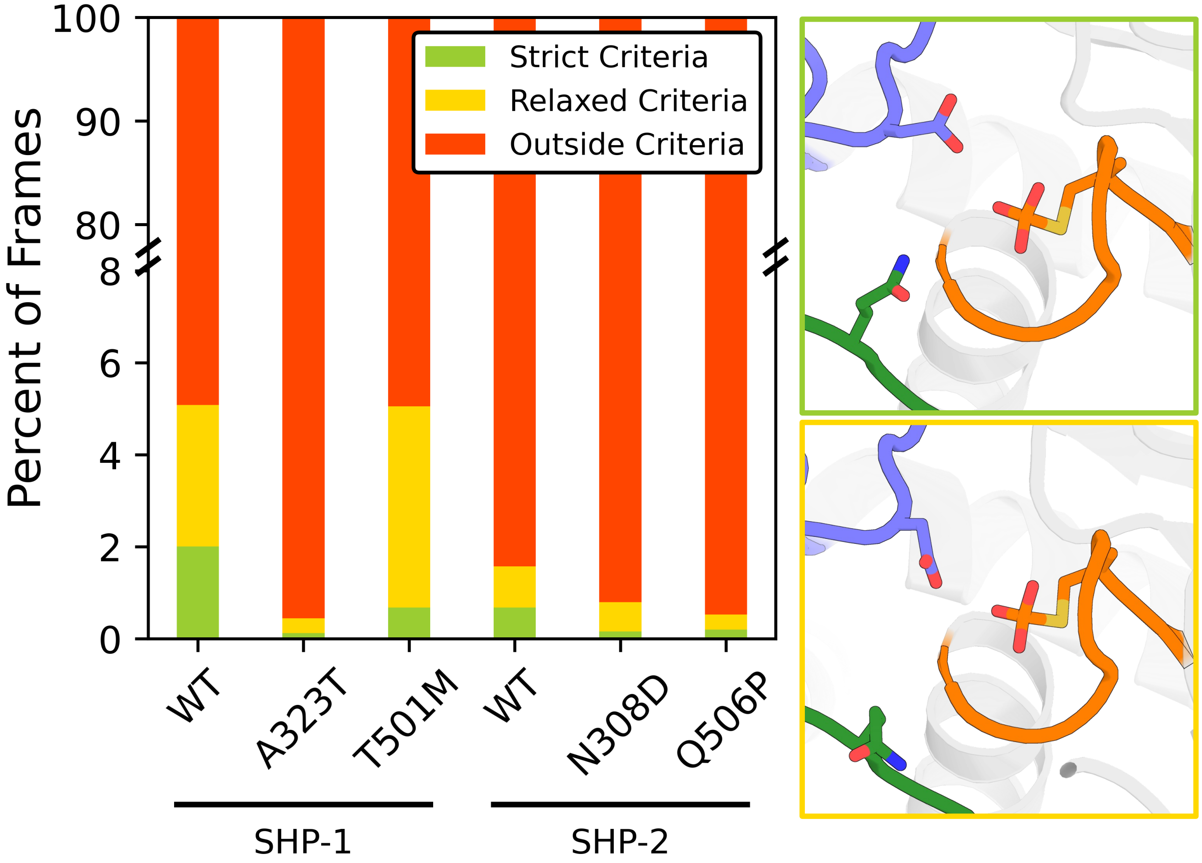


**Figure S24.** Percentage of molecular dynamics simulation frames in which the active site is found in a reactive conformation for hydrolysis, based on distances between the catalytic Asp on the WPD-loop, the Gln on the Q-loop, and the phosphorus atom at the phosphoenzyme intermediate. Structures were classified as obeying “strict criteria” (top right) if the Asp419(O)-pCys453(P) distance was less than 4.5Å and the Gln500(C)-pCys453(P) distance (SHP-1 numbering) was less than 8Å, in order to ensure that the Gln side chain was able to point towards the active site and stabilize the catalytic water molecule. Structures were further classified as obeying “relaxed criteria” (bottom right) if the Asp419(O)-pCys453(P) distance was less than 4.8Å and the Gln500(C)-pCys453(P) distance was less than 12Å.

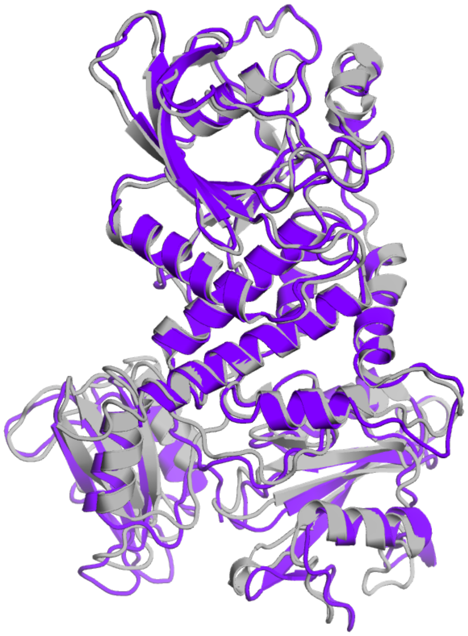

**Figure S25.** Alignment of the BioEmu[8] predicted structure of full-length SHP-1 (purple) with the corresponding crystal structure of active SH2-domain open SHP-1 (gray, PDB ID: 3PS5[1]). The root mean square deviation between the backbone heavy atoms of the two structures is 1.1Å, indicating that BioEmu is capable of predicting plausible active conformations of the full-length SHP PTPs.

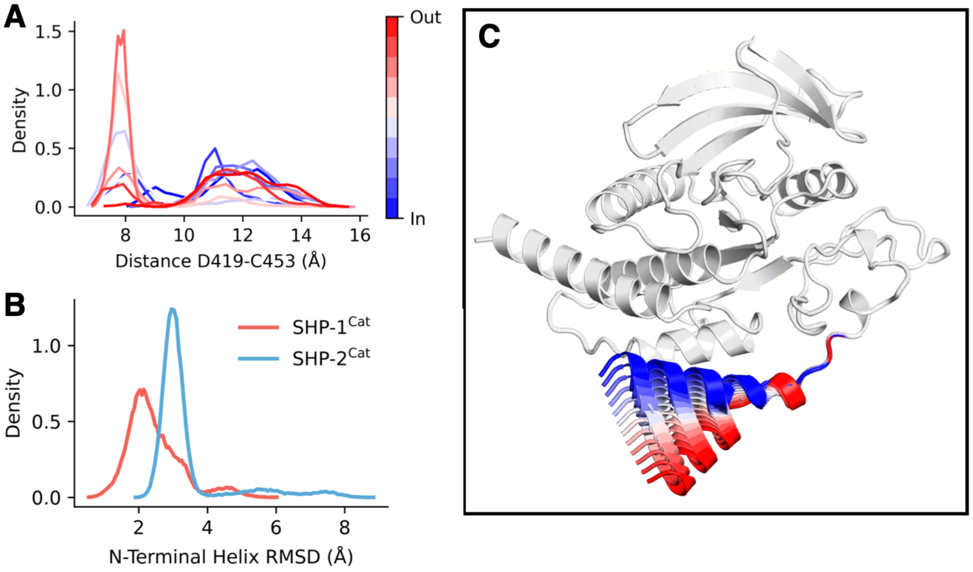


**Figure S26**. Benchmarking the effect of initializing N-terminal helix position on WPD-loop motion in SHP-1. (**A)** Probability distributions of the distance between $C_\alpha$-atom positions between D419 and C453 (Å) as a function of N-terminal helix starting position. (**B**) Probability distributions of the root mean squared deviations (RMSD, Å) of the catalytic domain SHP-1 and SHP-2 N-terminal helices. RMSD was calculated on all $C_\alpha$-atoms in the first 20 residues of SHP-1 and SHP-2 catalytic domain, after aligning on all of the stable structural elements of the catalytic domains. (**C**) Starting snapshots of helix positions were obtained using the Morph2 webserver (available at https://www.bioinformatics.org/pdbtools/morph2), using the helix position as defined from the PDB ID: 4GRZ[9] crystal structure as the 'in' conformation and the position as defined by the PDB ID: 4HJP[10] crystal structure as the 'out' conformation. 500ns simulations were run for each helix position seed.

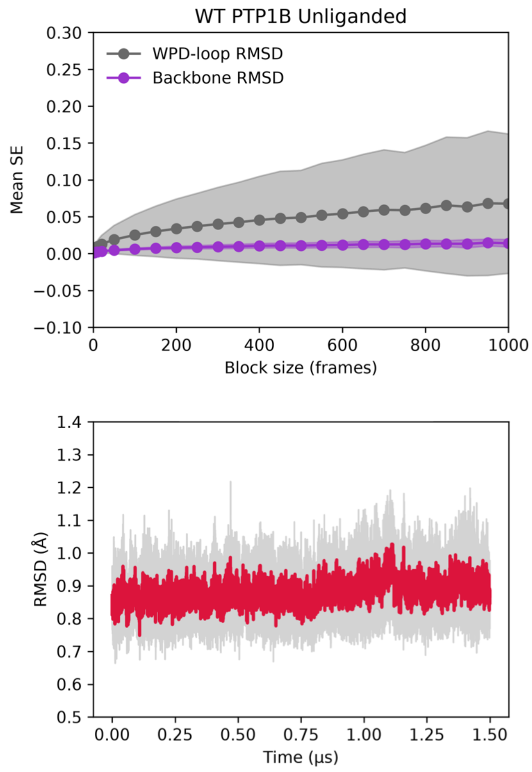


**Figure S27.** (**Top**) Block-averaging analysis for 8 x 1.5μs replica molecular dynamics simulations of unliganded PTP1B. Shown here are the block averaged values for the root mean squared deviation (RMSD) of the mobile WPD-loop, and the backbone $C_{\alpha}$-atoms. (**Bottom**) The RMSD calculated for all $C_{\alpha}$-atoms of the stable secondary structure regions (non-loops) of PTP1B during the simulations. The red lines indicate averages, and gray shading indicates standard deviations calculated over 8 x 1.5μs independent MD replicas.

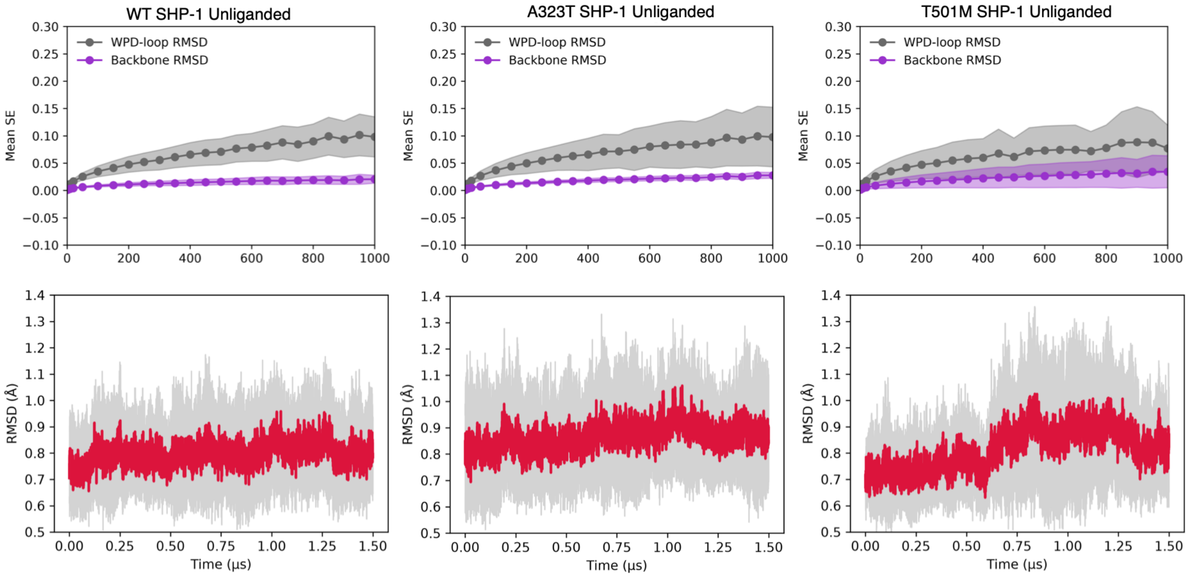


**Figure S28.** (**Top**) Block-averaging analysis for 8 x 1.5µs replica molecular dynamics simulations of unliganded wild-type and mutant forms of SHP-1. Shown here are the block averaged values for the root mean squared deviation (RMSD) of the WPD-loop, and the backbone $C_{\alpha}$-atoms. (**Bottom**) The RMSD calculated for all $C_{\alpha}$-atoms of the stable secondary structure regions (non-loops) of these PTPs during the simulations. The red lines indicate averages, and gray shading indicates standard deviations calculated over 8 x 1.5µs independent MD replicas.

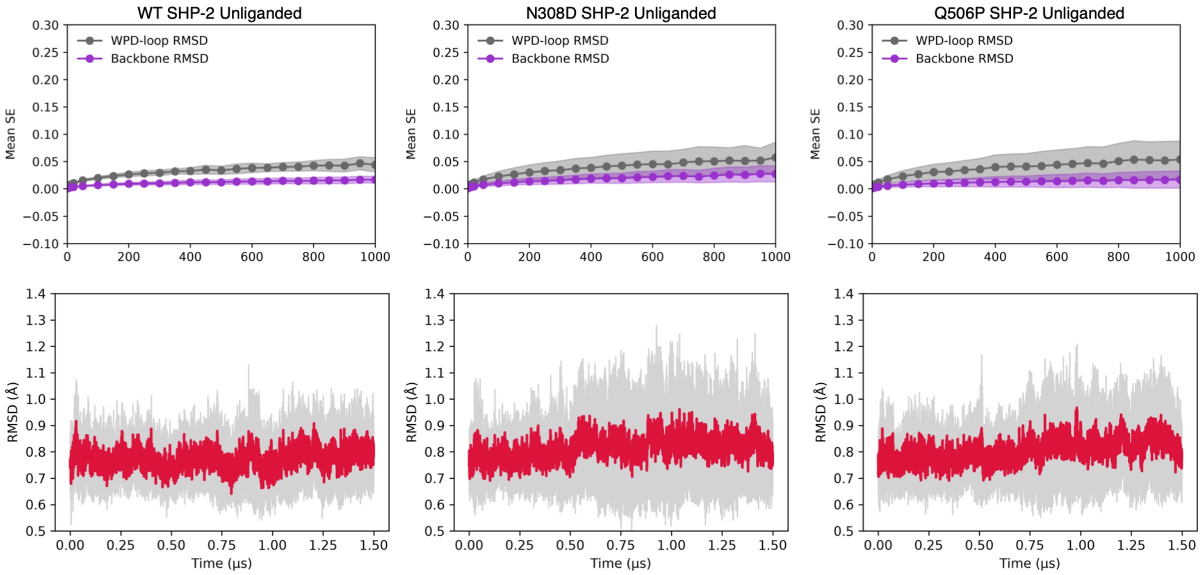


**Figure S29.** (**Top**) Block-averaging analysis for 8 x 1.5μs replica molecular dynamics simulations of unliganded wild-type and mutant forms of SHP-2. Shown here are the block averaged values for the root mean squared deviation (RMSD) of the WPD-loop, and the backbone $C_{\alpha}$-atoms. (**Bottom**) The RMSD calculated for all $C_{\alpha}$-atoms of the stable secondary structure regions (non-loops) of these PTPs during the simulations. The red lines indicate averages, and gray shading indicates standard deviations calculated over 8 x 1.5μs independent MD replicas.

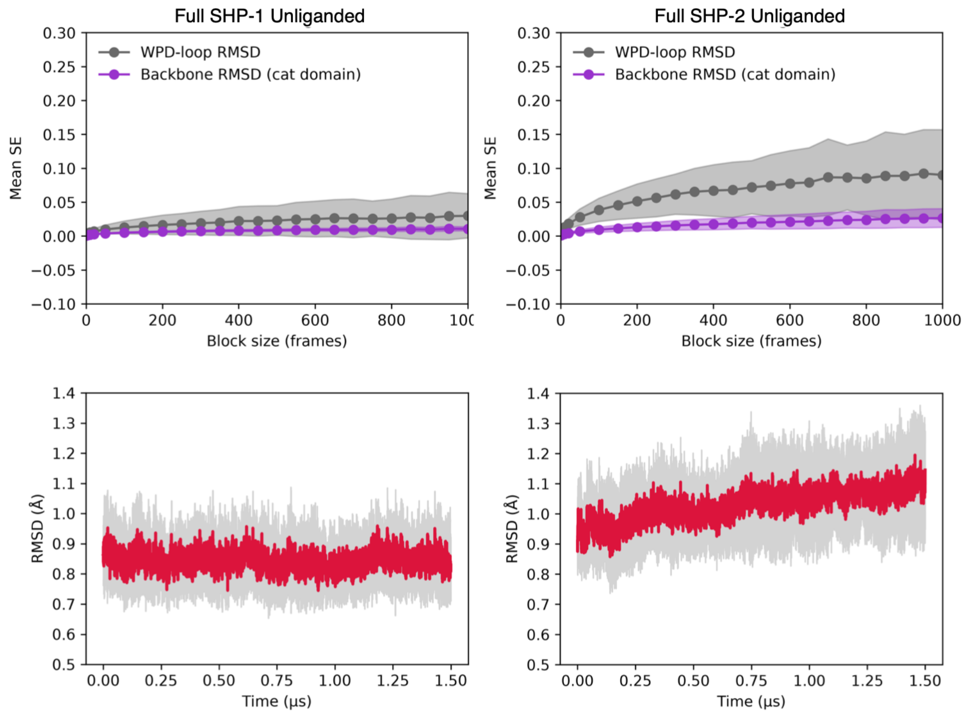


**Figure S30.** (**Top**) Block-averaging analysis for 8 x 1.5μs replica molecular dynamics simulations of unliganded full-length SHP-1 and SHP-2. Shown here are the block averaged values for the root mean squared deviation (RMSD) of the WPD-loop, which is highly mobile, and the backbone $C_\alpha$-atoms. (**Bottom**) The RMSD calculated for all $C_\alpha$-atoms of the stable secondary structure regions (non-loops) of these PTPs during the simulations. The red lines indicate averages, and gray shading indicates standard deviations calculated over 8 x 1.5μs independent MD replicas.

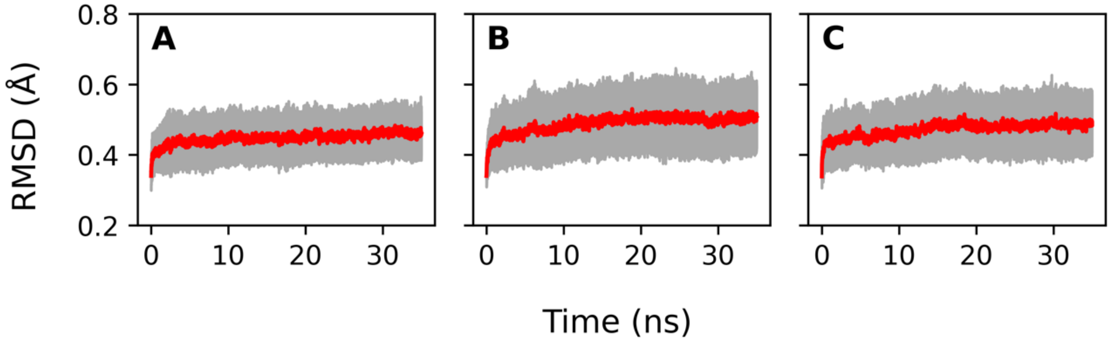


**Figure S31.** Root mean squared deviations (RMSD (Å)) calculated for all $C_\alpha$-atoms of the catalytic domain of wild-type SHP-1 (**A**), A323T variant (**B**), and T501M variant (**C**) over the course of 35ns equilibration before initiating empirical valence bond (EVB) simulations.[6] The red lines indicate averages and gray shading indicates standard deviations calculated over 30 x 35ns independent MD replicas.

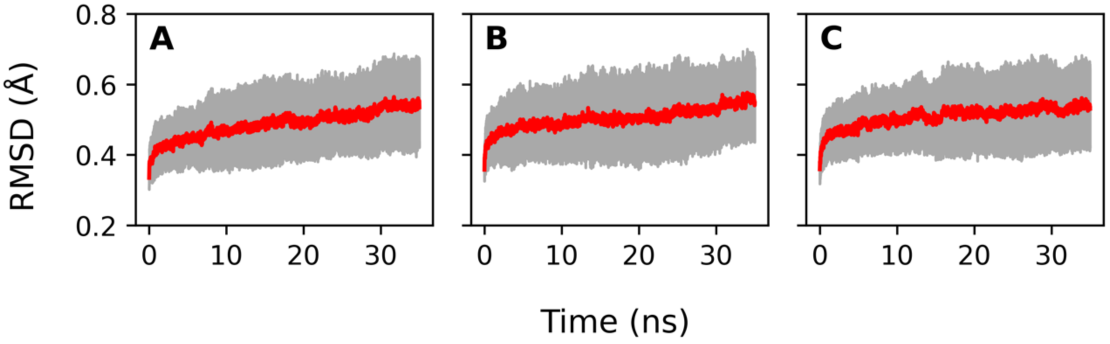


**Figure S32.** Root mean squared deviations (RMSD (Å)) calculated for all $C_\alpha$-atoms of the catalytic domain of wild-type SHP-2 (**A**), N308D variant (**B**), and Q506P variant (**C**) over the course of 35ns equilibration before initiating empirical valence bond (EVB) simulations.[6] The red lines indicate averages and gray shading indicates standard deviations calculated over 30 x 35ns independent MD replicas.

### Supplementary Tables

**Table S1**. Normalized linear mutual information scores between regions of interest identified in Figure 3 for PTP1B, SHP-1, and SHP-2.[a]

| Region pair | PTP1B | SHP-1 | SHP-2 |
|---|---|---|---|
| P-Loop – WPD-Loop | 0.25 ± 0.03 | 0.26 ± 0.03 | 0.21 ± 0.01 |
| E-Loop – WPD-Loop | 0.26 ± 0.02 | 0.29 ± 0.02 | 0.23 ± 0.01 |
| E-Loop – P-Loop | 0.32 ± 0.03 | 0.32 ± 0.02 | 0.31 ± 0.01 |
| α4-Helix – WPD-Loop | 0.18 ± 0.02 | 0.22 ± 0.01 | 0.19 ± 0.01 |
| α4-Helix – P-Loop | 0.27 ± 0.01 | 0.31 ± 0.01 | 0.31 ± 0.01 |
| α4-Helix – E-Loop | 0.20 ± 0.01 | 0.21 ± 0.01 | 0.21 ± 0.01 |
| α3-Helix – WPD-Loop | 0.27 ± 0.01 | 0.29 ± 0.01 | 0.29 ± 0.01 |
| α3-Helix – P-Loop | 0.19 ± 0.01 | 0.19 ± 0.01 | 0.21 ± 0.01 |
| α3-Helix – E-Loop | 0.20 ± 0.01 | 0.19 ± 0.01 | 0.22 ± 0.01 |
| α3-Helix – α4-Helix | 0.23 ± 0.01 | 0.21 ± 0.01 | 0.25 ± 0.01 |
| α5-Helix – WPD-Loop | 0.15 ± 0.02 | 0.21 ± 0.02 | 0.17 ± 0.01 |
| α5-Helix – P-Loop | 0.20 ± 0.01 | 0.26 ± 0.02 | 0.24 ± 0.02 |
| α5-Helix – E-Loop | 0.14 ± 0.01 | 0.23 ± 0.02 | 0.21 ± 0.02 |
| α5-Helix – α4-Helix | 0.21 ± 0.01 | 0.26 ± 0.02 | 0.24 ± 0.01 |
| α5-Helix – α3-Helix | 0.14 ± 0.01 | 0.16 ± 0.01 | 0.18 ± 0.02 |
| α7-Helix – WPD-Loop | 0.27 ± 0.02 | — | — |
| α7-Helix – P-Loop | 0.22 ± 0.01 | — | — |
| α7-Helix – E-Loop | 0.23 ± 0.01 | — | — |
| α7-Helix – α4-Helix | 0.18 ± 0.01 | — | — |
| α7-Helix – α3-Helix | 0.32 ± 0.01 | — | — |
| α7-Helix – α5-Helix | 0.20 ± 0.01 | — | — |

[a] Results are shown as mean values ± standard error of the mean (s.e.m.) across 8 x 1.5μs molecular dynamics simulations initiated from the unliganded state of each PTP, at the WPD-loop open configuration.

**Table S2**. Conservation of allostery between different PTPs considered in this work.[a]

| | **PTP1B** | **SHP-1** | **SHP-2** |
|---|---|---|---|
| **PTP1B** | -- | 52.1% | 41.3% |
| **SHP-1** | 24/46 | -- | 56.1% |
| **SHP-2** | 19/46 | 23/41 | -- |

[a] Fraction of allosteric "hotspot" residues shared between pairs of PTPs, calculated using the shortest path map (SPM)[11] approach, as well as the corresponding % of shared allosteric residues. Fractions are given with respect to the total number of SPM residues at the given column, and percentages are calculated from the corresponding fractions.

**Table S3**. Positions of disease-associated variants from the COSMIC[12] database that lie on or close to the shortest path map (SPM)[13] allosteric networks calculated for SHP-1 and SHP-2.[a]

| | **SHP-1** | **SHP-2** |
|---|---|---|
| **SPM pathway** | 323, 451, 501 | 265, 308, 498, 506, 507, 510 |
| **≤4Å from path** | 262, 275, 352, 362, 393, 419, 423, 453, 459, 462, 513 | 278, 285, 288, 309, 351, 357, 428, 454, 461, 468, 499, 502, 503, 505 512 |
| **>4Å from path** | 243, 316, 366, 367, 375, 399, 438 | 279, 282, 311, 319, 343, 363, 382, 384, 391, 397, 414 425, 433, 437, 491 |

[a] Only residue positions are noted, since there are often many mutations on the same positions which are marked as separate COSMIC entries. "SPM pathway" denotes positions directly on the allosteric networks, and "≤4Å from path" and ">4Å from path" indicate distance from the path.

**Table S4.** Experimental and calculated activation free energies, and corresponding experimental turnover numbers, for wild-type and variant forms of SHP-1 and SHP-2.[a]

| | $\Delta G^{\ddagger}_{calc}$ | $\Delta G^{\ddagger}_{exp}$ | $k_{cat}$ |
|---|---|---|---|
| SHP-1 | | | |
| SHP-1 WT | 15.1 ± 0.3 | 16.3 (pH 7.4), 14.2-15.1 (pH 5.0) | 5.7 (pH 7.4),[14] 57.6-200.9 (pH 5.0)[14-17] |
| A323T | 17.0 ± 0.4 | — | — |
| T501M | 16.3 ± 0.3 | — | — |
| SHP-2 | | | |
| SHP-2 WT | 15.7 ± 0.3 | 16.4 (pH 7.0), 16.1 (pH 7.5), 15.7-16.1 (pH 5.0) | 6.4 (pH 7.0)[18], 9.26 (pH 7.5),[19] 15.9-70.4 (pH 5.0)[15] |
| SHP-2 N308D | 17.1 ± 0.3 | 15.9 (pH 7.5) | 12.82 (pH 7.5)[19] |
| SHP-2 Q506P | 15.9 ± 0.3 | 17.7 (pH 7.0,7.5) | 0.61 (pH 7.5),[19] 0.64 (pH 7.0)[18] |

[a] References to experimental kinetic data are provided in the table. The $k_{cat}$ values are presented in $s^{-1}$, and all energies are presented in kcal $mol^{-1}$. All calculated activation free energies ($\Delta G^{\ddagger}$) are presented as average values and standard error of the mean over 30 independent EVB simulations. "—" indicates experimental data not available. Note that as in some cases, kinetics have been measured under a range of experimental conditions, as indicated in the table.

**Table S5.** Calculated distances at the phosphoenzyme intermediate (RS), transition states (TS), and product states (PS) as obtained from our EVB simulations.[a]

| System | State | $S_{cys}$-P | P-$O_{H2O}$ | $O_{H20}$-H | H-$O_{Asp}$ |
|---|---|---|---|---|---|
| SHP-1 WT | RS | 2.0±0.0 | 3.2±0.2 | 1.0±0.0 | 3.0±0.8 |
| | TS | 2.5±0.6 | 2.1±0.2 | 1.2±0.2 | 1.2±0.1 |
| | PS | 3.2±0.0 | 1.6±0.0 | 1.3±0.0 | 1.0±0.0 |
| SHP-2 A323T | RS | 2.0±0.0 | 3.4±0.3 | 1.0±0.0 | 2.6±0.9 |
| | TS | 2.6±0.7 | 2.0±0.2 | 1.2±0.3 | 1.2±0.1 |
| | PS | 3.3±0.1 | 1.6±0.0 | 1.3±0.0 | 1.0±0.0 |
| SHP-1 T501M | RS | 2.0±0.0 | 3.6±0.6 | 1.0±0.0 | 2.4±1.0 |
| | TS | 2.6±0.5 | 2.0±0.2 | 1.2±0.2 | 1.2±0.1 |
| | PS | 3.3±0.1 | 1.6±0.0 | 1.3±0.0 | 1.0±0.0 |
| SHP-2 WT | RS | 2.0±0.0 | 3.7±0.8 | 1.0±0.0 | 2.8±1.0 |
| | TS | 2.4±0.2 | 2.2±0.1 | 1.2±0.4 | 1.2±0.0 |
| | PS | 3.3±0.1 | 1.6±0.0 | 1.3±0.0 | 1.0±0.0 |
| SHP-2 N308D | RS | 2.0±0.0 | 3.5±0.5 | 1.0±0.0 | 2.8±1.1 |
| | TS | 2.5±0.4 | 2.1±0.1 | 1.1±0.1 | 1.2±0.1 |
| | PS | 3.3±0.1 | 1.6±0.0 | 1.3±0.0 | 1.0±0.0 |
| SHP-2 Q506P | RS | 2.0±0.0 | 3.5±0.6 | 1.0±0.0 | 2.8±1.1 |
| | TS | 2.4±0.2 | 2.1±0.1 | 1.1±0.1 | 1.2±0.0 |
| | PS | 3.3±0.2 | 1.6±0.0 | 1.3±0.0 | 1.0±0.0 |

[a] Distances are measured in Å and are shown as average values ± standard deviation over 30 independent EVB trajectories for each system.

**Table S6.** Overview of crystal structures used in this work to simulate SHP-1 and SHP-2.

| PDB ID | Protein | WPD-loop Conformation | Domains |
|---|---|---|---|
| 4GRZ[9] | SHP-1 | Closed (unliganded and intermediate) | Catalytic domain |
| 4HJP[10] | SHP-1 | Open (unliganded and intermediate) | Catalytic domain |
| 4RDD[20] | SHP-1 | Open | Catalytic domain |
| 6CMQ[21] | SHP-2 | Closed/Open (unliganded and intermediate) | Catalytic domain (N-SH2 removed) |
| 3PS5[1] | SHP-1 | Open (unliganded) | Catalytic domain + N-SH2 + C-SH2 (open) |
| 2B3O[2] | SHP-1 | Open (unliganded) | Catalytic domain + N-SH2 + C-SH2 (autoinhibited) |
| 4DGP[22] | SHP-2 | Open (unliganded) | Catalytic domain + N-SH2 + C-SH2 (autoinhibited) |
| 6B90[23] | PTP1B | Closed/Open (unliganded and open intermediate) | Catalytic domain |
| 3I80[24] | PTP1B | Closed (intermediate) | Catalytic domain |

**Table S7.** Ionization residues and histidine protonation patterns in the empirical valence bond (EVB) simulations of SHP-1 and SHP-2.[a]

| Residue Type | SHP-1 | SHP-2 |
|---|---|---|
| Asp | 334, 386, 419, 434 | 395, 425, 431, 473, 477 |
| Glu | 249, 255, 329, 353, 355, 384, 389, 425, 502 | 258, 359, 361, 390 |
| Arg | 262, 275, 292, 352, 358, 393, 459, 492, 495 | 278, 362, 399, 421, 465, 498, 501 |
| Lys | 273, 277, 356, 360, 391, 506 | 325, 358, 364, 366 |
| His-δ | 411, 452 | 287, 443, 458, 520 |
| His-ε | 260, 284, 385, 420, 445 | 293, 394, 419, 426 |

[a] All other residues were kept in their neutral state, as they fell outside the explicit simulation sphere using the surface constraint all atom solvent model (SCAAS).[25] Equivalent ionization states were used for all oncogenic variants.
